# Central Nervous System axonal regeneration by spatially targeted drug combinations

**DOI:** 10.1101/157099

**Authors:** Mustafa M. Siddiq, Nicholas P. Johnson, Yana Zorina, Arjun Singh Yadaw, Carlos A. Toro, Jens Hansen, Vera Rabinovich, Sarah M. Gregorich, Yuguang Xiong, Rosa E. Tolentino, Sari S. Hannila, Ehud Kaplan, Robert D. Blitzer, Marie T. Filbin, Christopher P. Cardozo, Christopher L. Passaglia, Ravi Iyengar

## Abstract

There are no known drugs or drug combinations that promote substantial central nervous system axonal regeneration after injury. We used systems pharmacology approaches to model pathways underlying axonal growth and identify a four-drug combination that regulates multiple subcellular processes in the cell body and axon using the optic nerve crush model in rats. We intravitreally injected agonists HU-210 (cannabinoid receptor-1) and IL-6 (interleukin 6 receptor) to stimulate retinal ganglion cells for axonal growth. We applied, in gel foam at the site of nerve injury, Taxol to stabilize growing microtubules, and activated protein C to clear the debris field since computational models predicted that this drug combination regulating two subcellular processes at the growth cone produces synergistic growth. Morphology experiments show that the four-drug combination promotes axonal regrowth to the optic chiasm and beyond. Physiologically, drug treatment restored pattern electroretinograms and some of the animals had detectable visual evoked potentials in the brain and behavioral optokinetic responses. We conclude that spatially targeted drug treatment can promote robust axonal regeneration and can restore limited functional recovery.

## Introduction

Injury in the adult CNS is often permanent due to inability of severed axons to regenerate. This inability has two major causes: extracellular factors, including myelin-associated molecules, which inhibit axonal outgrowth^1–5^; and lack of appropriate activation of intracellular signaling pathways. Among these are the MTOR and STAT3 pathways.^6–9^ Simultaneous genome-level activation of these pathways led to robust and sustained axonal regeneration^10^. Likewise, sustained activation of the MTOR pathway by genetic manipulation, combined with visual stimulation, leads to extended regeneration of optic nerve axons to the brain.^11^ Transcriptomic analyses of dorsal root ganglion neurons following peripheral nerve injury have identified the involvement of numerous signaling pathways including neurotrophins, TGFβ cytokine, and JAK-STAT.^11^ Using these transcriptomic data and the Connectivity Map, researchers identified the drug ambroxol, a Na^+^ channel inhibitor that promoted modest axon regeneration after optic nerve crush in mice.^12^ Despite these extensive studies, no drug or drug combinations have been identified that result in substantial axonal regeneration leading to restoration of physiological function have been identified.

We have been studying signaling through the Go/i pathway for over two decades and found that activated Gαo activates STAT3 to promote oncogenic transformation. Studies of Neuro-2A cells treated with a cannabinoid receptor-1 (CB1R) agonist indicated that this receptor — acting through the GTPases Rap, Ral, and Rac — activates Src and STAT3 to stimulate neurite outgrowth.^8,13,14^ An extensive study identified a complex upstream signaling network that controls multiple transcriptional factors including STAT3 and CREB to regulate neurite outgrowth.^15^ These studies indicated that several types of receptors regulate STAT3 to drive neurite outgrowth. IL-6 promotes neurite outgrowth in a PC-12 variant and overcomes myelin-mediated inhibitors to promote neurite outgrowth in primary neurons.^7,16^ Previously, we showed that submaximal, potentially therapeutic concentrations of the CB1R agonist HU-210 with IL-6 can activate STAT3 in a sustained manner and induce neurite outgrowth in both Neuro2A cells and primary rat cortical neurons in an inhibitory environment *in vitro*.^17^ These drugs have never been studied *in vivo*. Initial reasoning indicates that IL-6 might be deleterious as it is proinflammatory. However, at appropriate concentrations its beneficial effects could be resolved from its proinflammatory effects.

We anticipated that stimulating signaling and transcription in the cell body of injured neurons to enable growth in inhibitory environments alone probably would not lead to long-distance axon regeneration to the brain. **Hence, we sought to develop a systems-level combination therapy based on spatial specification of drug action both at the cell body and at the site of injury**. We used our *in vitro* experiments to identify a 2-drug combination that at low concentrations could stimulate biosynthetic processes in the injured neuron *in vivo*. Using predictions from computational models of neurite outgrowth we identified drugs that could selectively modulate subcellular function in the cell body and axon.^18^ We then utilized this spatial information to develop a multi-drug combination with the potential for treating CNS nerve injury by promoting long-range axonal regeneration *in vivo*. Since transcriptional regulation is critical for neurite outgrowth and for axonal regeneration,^15^ we reasoned that stimulation of CB1R and IL-6Rs occurs in the cell body. We then looked for subcellular processes in the axon that could be potential drug targets and utilized computational dynamical models of neurite outgrowth to predict how different subcellular processes could promote axonal regeneration.^18^ Based on these simulations we focused on microtubule growth, which is required for axon regeneration. Microtubules are stabilized by Taxol by *de facto* reducing the intrinsic GTPase turnover rate of tubulin.^19^ Increasing conversion of dynamic to stable microtubules increases axonal length to promote regeneration.^20^ As axonal regeneration is inhibited by cell remnants at the site of injury, we hypothesized that such inhibition could be in part due to regulation of incorporation of membrane vesicles at the growth cone based on the ability of cell debris to bind the growth cone cell surface glycoproteins.^21^ Since computational modeling showed that regulation of vesicle fusion dynamics is an important regulator of neurite length, treating the site of injury to clear it of inhibitory agents and reduce the inflammatory response could also contribute to axonal regeneration. Based on these considerations, we chose a protease that could act locally to reduce the debris field and inhibit inflammation. We selected activated protein C (APC), which is a serine protease endogenous to the coagulation system that is anti-inflammatory and promotes stem cell activity in neuronal tissue in mice.^22,23^ Thus, we identified a four-drug combination of which two had never been tested *in vivo* for axonal regeneration. The overall logic for this four-drug combination is schematically shown in Fig. 1. We tested if a combination of these drugs, two applied at the cell body and two at the axon at the site of injury, promoted long distance regeneration such that visual stimulation led to restoration of physiological function.

**Figure 1.**
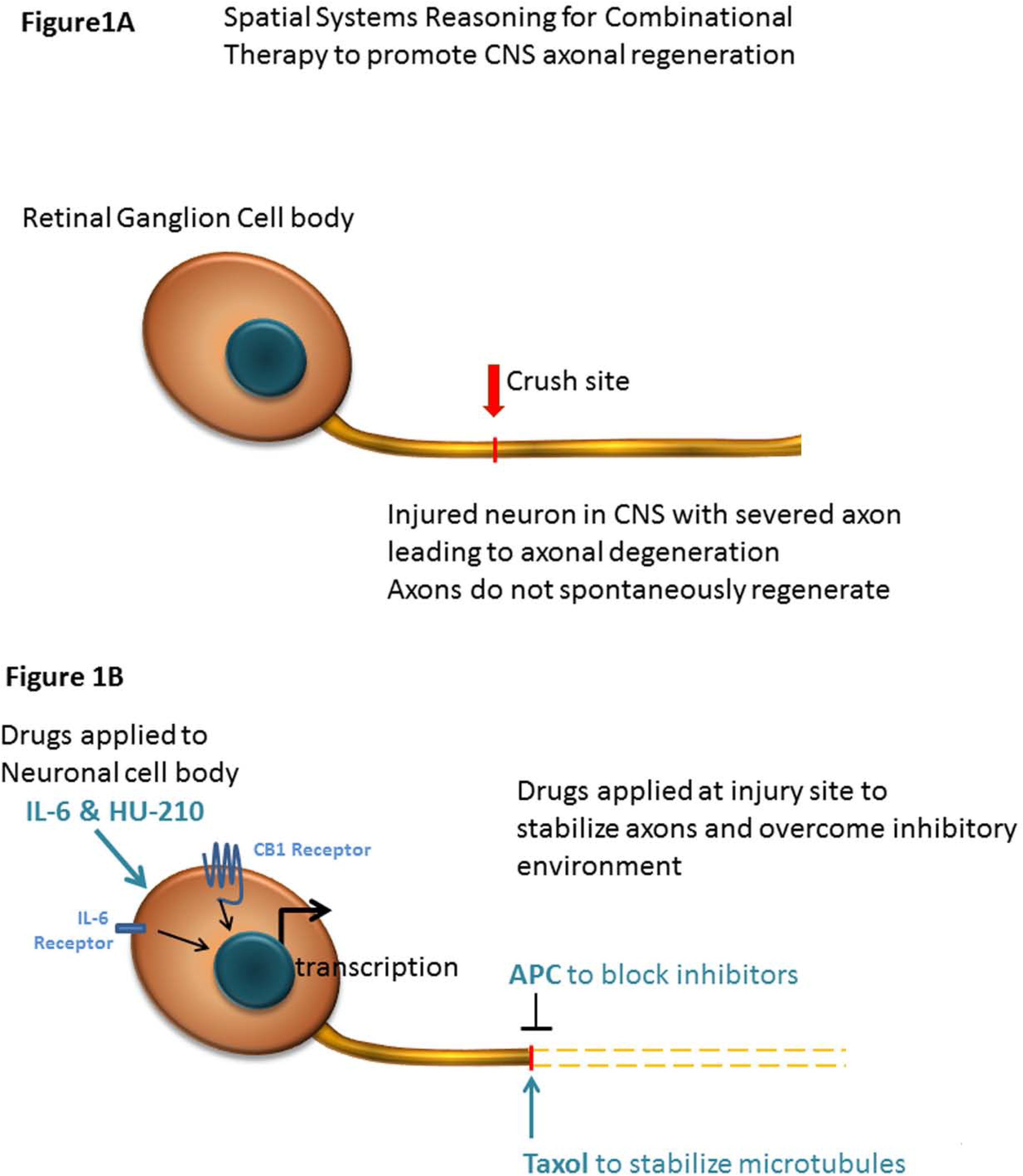
Experimental Summary. Cartoon schematic for spatial systems reasoning for combinational therapy to promote CNS axonal regeneration. Drugs are applied to both the injury site (APC and Taxol) and directly to the RGC cell bodies (IL-6 & HU-210)

## Materials and Methods

### Care and Use of Animals Statement

These experiments were conducted under the guidance of the Institutional Animal Care and Use Committee established in the Icahn School of Medicine at Mount Sinai and in accordance with federal regulations and the guidelines contained in the National Research Council Guide for the Care and Use of Laboratory Animals. To replace the use of animals, when possible, rigorous computational models and *in vitro* experiments were employed prior to *in vivo* experiments. Likewise, the optic nerve crush model was chosen to deliver robust and reproducible results while minimizing the distress animals experience during these experiments. Adult Sprague-Dawley or Long Evans rats (250-280 g, 8-10 weeks old), as well as postnatal day 1 Sprague-Dawley rats, were used in these experiments. Animals were housed in individually ventilated cages under the supervision of the Icahn School of Medicine Center for Comparative Medicine and Surgery and kept on a 12 h light/dark cycle. During surgeries, deep anesthesia was induced using isoflurane (Piramal Pharma Solutions, Sellersville, PA), and post-surgical pain was managed with slow-release buprenophine HCL (Par Pharmaceutical, Woodcliff Lake, NJ).

### Rat primary cortical neuron cultures

Rat primary cortical cultures were dissected from postnatal day 1 Sprague Dawley rat brains. Cortices were incubated twice for 30 min with 0.5 mg/ml papain in plain Neurobasal (NB) media (Invitrogen, Waltham, MA) with DNase (Sigma, St. Louis, MO). Cell suspensions were layered on an Optiprep density gradient (Sigma) and centrifuged at 1900 x *g* for 15 min at room temperature (22°C). The purified neurons were then collected at 1000 x *g* for 5 min and counted.

Suspensions of purified CNS myelin (1-2 μg/well) were plated in chamber slides and desiccated overnight to create a substrate of myelin on eight well chamber slides as previously described.^24,25^ Primary cortical neurons were diluted to 35,000 cells/ml in NB supplemented with B27, L-glutamine, and antibiotics (Thermo Fisher Scientific, Waltham, MA). 300 μl of the cell suspension was seeded to each well and incubated for 24 h. To quantify the outgrowth, we immunostained the neurites using a monoclonal anti βIII tubulin antibody (BioLegend, San Diego, CA) and Alexa Fluor 568-conjugated anti-mouse IgG (Invitrogen). Outgrowth was quantified by measuring the total fluorescence at multiple cross-sections of the image. The zero point on the x-axis represents the right edge of microgrooves. All data points were normalized to the first point of “No MLN.”

7 days in vitro (DIV), axonal injury was modeled by axotomy. The media was aspirated from the axonal chambers with the tip of the aspirator held against the microgrooves. This was repeated three times per chamber. Fresh supplemented NB media containing 10ng/ml IL-6, 200 nM HU-210, or a combination of both, was supplied to the chambers and live imaging was conducted the following day.

For Western Blots, primary neurons were treated with IL-6, HU-210, or both and lysed in radioimmunoprecipitation assay (RIPA) buffer (Cell Signaling Technology, Danvers, MA) supplemented with protease and phosphatase inhibitors (Thermo Fisher Scientific) over ice. After determination of concentration with a standard protein assay kit (Bio-Rad, Hercules, CA), the cell lysates were subjected to immunoblot analysis using standard procedures and visualized by enhanced chemiluminescences (ECL). Polyclonal rabbit antibodies directed against phosphor-STAT3 and total STAT3 were from Cell Signaling Technology Inc.

### Rat optic nerve regeneration *in vivo*

Adult male Sprague-Dawley rats were anesthetized with isoflurane and placed in a stereotaxic frame. The right optic nerve was exposed and crushed with fine forceps for 10 s. For drug treatment, animals received a single 2.5 μl intravitreal (intraocular) injection of either 0.5% DMSO or a combination of IL-6 (5 μg/ml) and HU-210 (300 μM) (see Table 1) immediately after the crush. In later experiments, animals also received gelfoam soaked in DMSO, APC (4.1 mg/ml) (Haematologic Technologies, Essex, VT), and/or Taxol (5 μM) placed over the injury site. Three days later, we confirmed that lens injury was avoided and injected for a second time intravitreally 2.5 μl of DMSO or IL-6 (5 μg/ml) and HU-210 (300 μM). One week prior to sacrificing the animals we labelled the regenerating axons with 5 μl of 1 mg/ml cholera toxin B (CTB, List Biological Lab, Campbell, CA) coupled to Alexa-488 (Invitrogen), which we intravitreally injected. On day twenty-one post-crush, animals were anesthetized with ketamine (100 mg/kg) and xylazine (20 mg/kg) injected intraperitoneally, and the brain was fixed by transcardial perfusion with cold 4% PFA in PBS (pH 7.4). The optic nerves and chiasm attached were dissected out and post-fixed in 4% PFA overnight at 4°C, rinsed for one hour in ice-cold PBS, and then prepared for chemical clearing. Since the advent of the 3DISCO clearing techniques,^26^ we are no longer dependent on sectioning the tissue; this method also eliminates bias associated with artifacts produced by sectioning. The whole nerve was exposed to a graded series of dehydrations with tetrahydrofurane (THF; Sigma) diluted in water, final concentrations THF: 50%, 70%, 80%, 100%, and 100% again. The nerves were incubated in each solution for 20 minutes at room temperature on an orbital shaker, and then incubated in the clearing agent, dibenzyl ether (DBE; Sigma) overnight at room temperature on the orbital shaker. Microscope slides were mounted with Fastwell chambers (Electron Microscopy Sciences, Hatfield, PA) and the cleared sample was placed on the slide and covered with DBE and a No. 1.5 micro coverglass. We imaged the whole sample on an Olympus Multiphoton microscope with a 25X water immersion lens. Optic nerve immunostaining was alternatively conducted using an anti-GAP-43 antibody (Novus Biologicals, Centennial, CO) coupled to Alexa-488 (Invitrogen) or using an anti-GFP antibody (Abcam, Cambridge, UK) coupled to Alexa-488 (Invitrogen). In the final case, AAV8-GFP was injected intravitreally 1 week prior to sacrificing the animals.

For quantification of axonal regeneration in our ONC samples, multiple images of each sample were taken under multiphoton microscopy and combined in photomontages using Adobe Photoshop (Adobe, New York, NY). We used ImageJ software (NIH) to measure the axonal outgrowth determined by immunolabeling in our regenerated nerves in photomontages.^24^ Pixel thresholding was performed to identify labeled axons, and we measured in 250 micron increments from the crush site. We graphed the Intensity of Fluorescent signal (area occupied) over distance (μm) from the crush site using GraphPad Prism Software (GraphPad, San Diego, CA).

### Electrophysiological recordings

All electrophysiology was performed using Long Evans (pigmented) rats. Pattern electroretinogram (pERG) recordings were performed at the Icahn School of Medicine at Mount Sinai. Rats were anesthetized with intraperitoneal ketamine (100 mg/kg) and xylazine (20 mg/kg) 21 days post-surgery and positioned in a device designed for measuring pERGs (UTAS-2000; LKC Technologies, Gaithersburg, MD). Contact lenses embedded with gold electrodes were placed on the corneas, and light-adapted responses of both eyes were recorded to a contrast-reversing checkerboard pattern displayed on a monitor located 11 cm in front of the corneas.^27^ Responses to 300 pattern reversals (spatial frequency: 0.033 cycles/degree; contrast: 100%, temporal frequency: 2 Hz) were averaged to estimate pERGs. Flash ERG (fERG) and visually evoked potential (VEP) recordings were performed at the University of South Florida.^28^ Rats were anesthetized with intraperitoneal ketamine (75 mg/kg) and xylazine (5 mg/kg) and placed on a heating pad. Cannulas were surgically inserted into the femoral vein for intravenous drug delivery and the trachea for mechanical ventilation if necessary. Anesthesia was maintained with intravenous infusion (all dosages are mg/kg/hr) of ketamine (50), xylazine (1.5), dextrose (600), and physiological saline. Heart rate was monitored with ECG electrodes placed in the trunk of the animal, and body temperature was monitored with a rectal thermometer. The eyes were covered with clear contact lenses, and pupils were dilated with mydriatic eye drops (cyclopentolate hydrochloride, Sandoz, Basel, Switzerland). Flash stimuli were produced by a green LED encased in an opaque tube the blocked the escape of light and the tube exit was fitted with a hemispherical diffuser that covered the eye to provide Ganzfeld illumination.^28^ fERGs were recorded with a ring-shaped gold electrode placed on the cornea. Platinum needle electrodes were inserted in the temples and tail to serve as reference and ground, respectively. VEPs were recorded from both hemispheres via two 1.3 mm steel screws inserted through the skull 7 mm posterior to Bregma and ± 2.5 mm lateral to the midline. Recorded signals were differentially amplified (10,000X) and filtered (0.1 – 1,000 Hz) by a multichannel bioamplifier and digitized at 1 kHz. Animals were dark-adapted for 15 minutes prior to data collection, and then baseline fERG and bilateral VEP data were collected simultaneously for a series of 100 brief (10 ms) flashes with an interstimulus interval of 3 s to allow for recovery of visual sensitivity. Average responses were measured for a flash series of increasing light intensity in seven logarithmic steps to a maximum of 1.32 log candelas/m^2^/s (cd·s/m^2^, 100% condition) presented to one eye and then the other. Since approximately 90% of the fibers in the rat optic nerve cross to the opposite side to innervate the brain, when we flashed light and recorded from the right injured optic nerve, we recorded the corresponding VEP on the left side of the brain. All procedures were approved by the IACUCs of the Icahn School of Medicine at Mount Sinai and University of South Florida at Tampa in accordance with the NIH guidelines and AAALAC accreditation.

### Optokinetic Responses

Awake rats were placed on an elevated platform in a virtual reality chamber composed of four computer monitors facing inward (OptoMotry system, Cerebral Mechanics Inc, Lethbridge, AB, Canada). A virtual cylinder covered with a vertical sine wave grating was presented to the rat and a video camera situated above the animal provided real-time feedback to the observer. The spatial frequency of the sine was maintained by manually repositioning the center of the cylinder at the animal’s head. The sine wave drifted clockwise or counterclockwise and the observer judged whether the rat tracked the grating with slow movements of its head and body, with the direction of the stimulus alternating randomly between trials. Spatial frequencies were increased if tracking was observed and decreased if no tracking was observed. Visual thresholds were obtained using a staircase method in which spatial frequency steps were halved after a reversal and terminated after 10 reversals. One staircase was performed in each direction as each direction elicited an independent response from each eye, with the clockwise direction corresponding to the left eye and the counterclockwise direction corresponding to the right eye. The observer was blind to the direction of the stimulus, as well as the spatial frequency and the number of reversals.

### Statistical Analysis

Except where noted, analyses were performed using GraphPad Prism software (GraphPad, San Diego, CA), and data are represented as mean ± SEM. Statistical significance was assessed using paired one-tailed Student’s *t* tests to compare two groups, and one-way ANOVAs with Bonferroni’s *post hoc* tests to compare among three or more groups.

To evaluate the effectiveness of the four-drug treatment in restoring the VEP following injury, we analyzed traces from each animal to detect physiological responses that exceeded RMS noise by a Z-score of at least three. We set this as a criterion to reliably eliminate false positives, as we could not distinguish two signal-free (RMS noise) traces at a Z-score of 3. RMS noise was calculated for each time point using the detrended traces recorded at the three lowest stimulus intensities (0.001%, 0.01%, and 0.1%), since none of the injured nerves showed apparent VEPs at these intensities. For each animal, the mean of these designated noise traces was normalized to 0 mV. We then measured the peak amplitudes of the responses to stimulation at 10%, 50%, and 100%, and these were converted to Z-scores against the mean noise at the corresponding time point. The presence of a VEP was defined by Z-score > 3. Chi-square analysis was used to compare the number of treated and untreated animals that showed VEPs in response to both the 10% and 100% stimuli.

## Results

### IL-6 and HU-210 together stimulates neurite outgrowth in an inhibitory environment

We previously found that submaximal concentrations of IL-6 and HU-210 in combination have an additive effect on neurite outgrowth for rat cortical neurons in primary culture.^17^ We tested if this effect could be observed for regeneration of growing neurites that had been severed *in vitro* and grown in an inhibitory environment. For these experiments, we used microfluidic chambers, where the cell bodies could be compartmentalized from the growing axons (Fig. 2A1-2). We plated primary rat cortical neurons in these chambers and allowed them to grow long neurites. To mimic axotomy, we lesioned all the neurites in the chambers on the right side and then added 200nM HU-210 and 10ng/ml IL-6 to the somal chamber (Fig. 2A2) with the addition of 20μg/ml myelin (MLN), a non-permissive substrate (Supplementary Figure 2) used to inhibit axonal growth. We observed that addition of the combination of IL-6 and HU-210 to the cell bodies promoted longer growth of axotomized processes in the presence of MLN, compared to either treatment alone. Controls and treatments with IL-6 and HU-210 in the absence of MLN are shown in Supplementary Fig. 2A.

**Figure 2.**
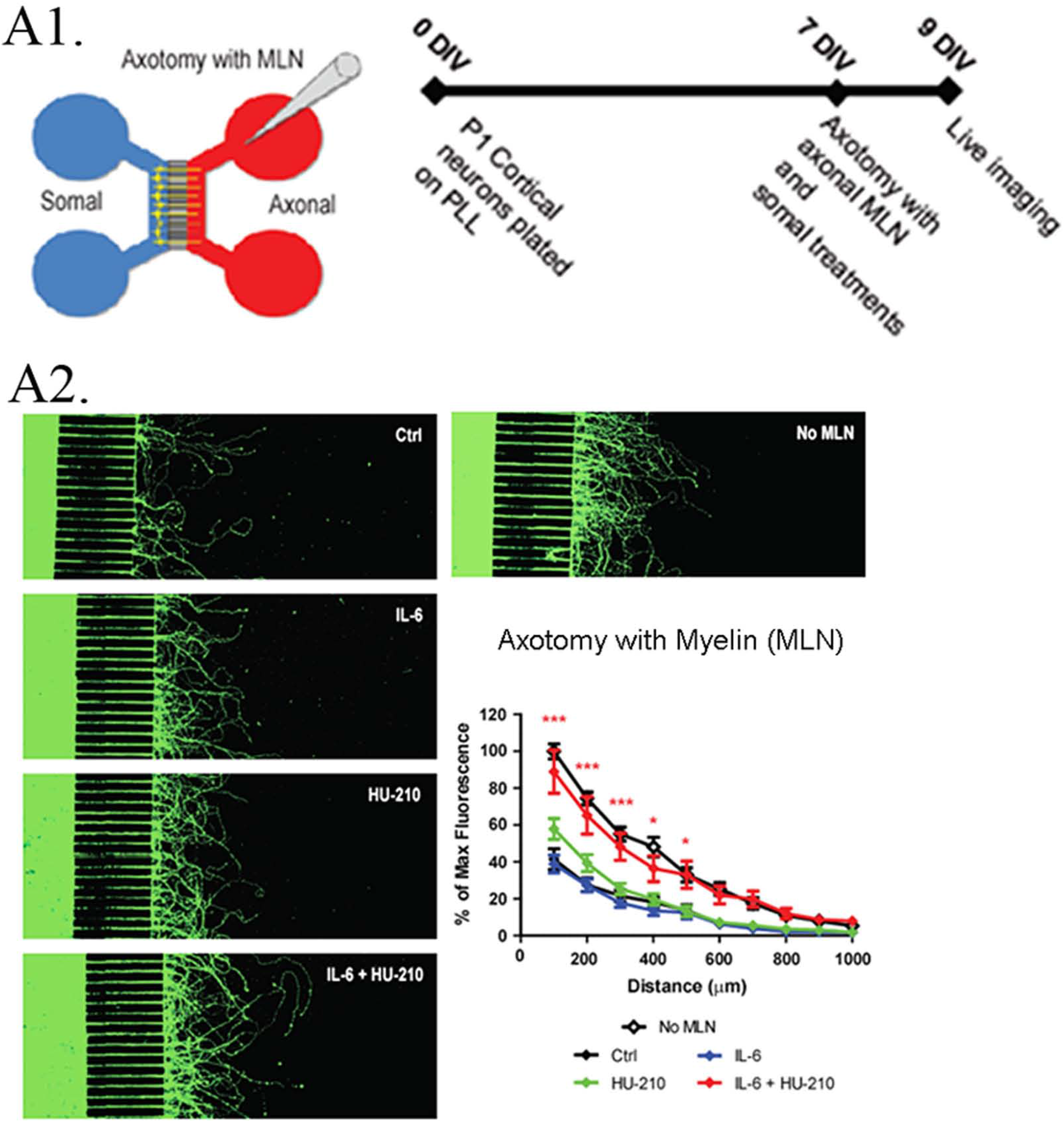
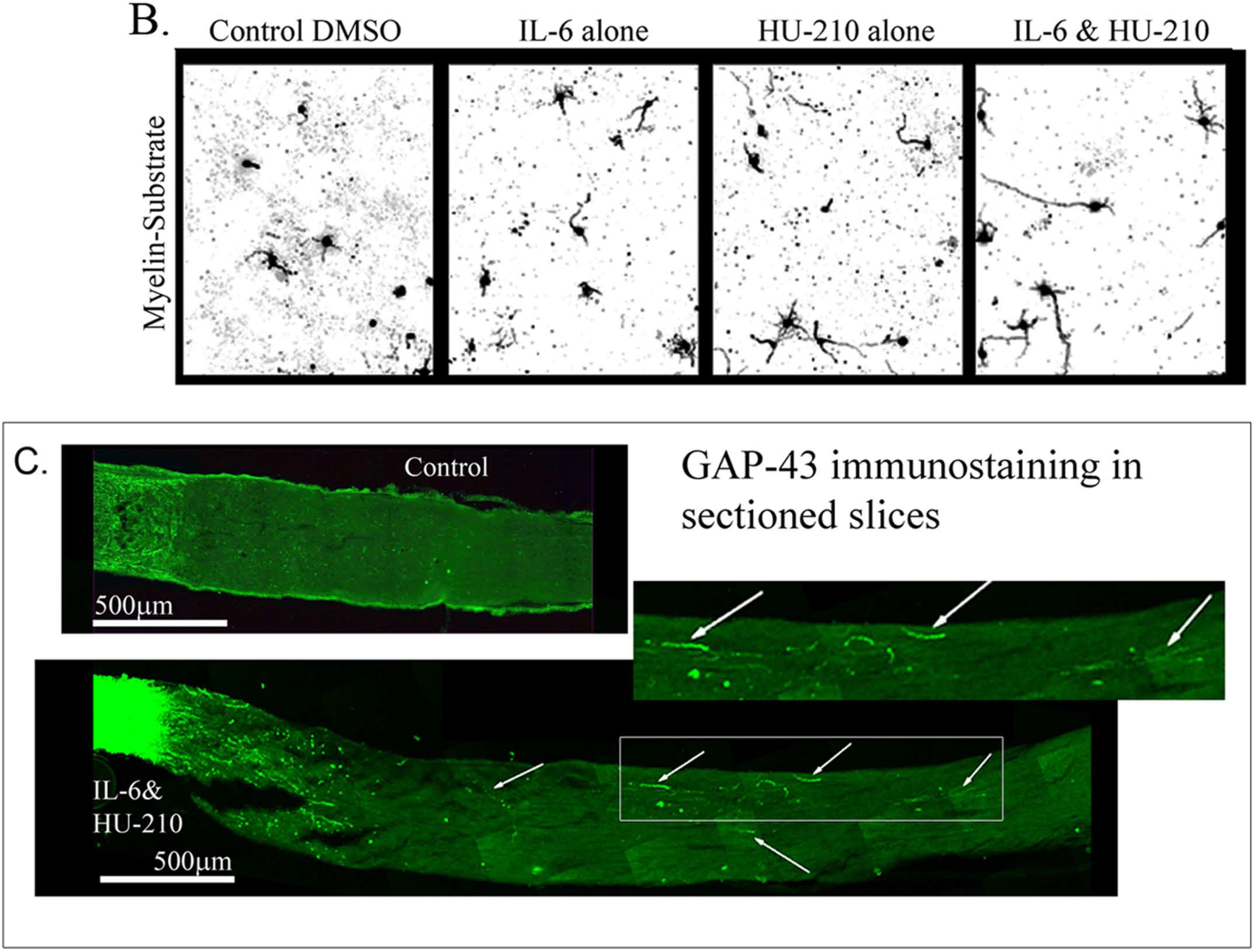
Somal treatment with IL-6 and HU-210 promotes neurite outgrowth on myelin. (A1) Schematic representation of the microfluidic chamber. (A2) Neurons were cultured in chambers on PLL for 7 days *in vitro* (DIV), and axotomy was performed. The axonal compartment was then filled with 20 μg/ml solution of myelin (MLN). 10 ng/ml IL-6 and/or 200nM HU-210 treatments were applied to the somal compartment of each respective chamber. The neurons were imaged live after 48 hrs using Calcein AM. Representative confocal images of neurons following axotomy and treatment. Neurite outgrowth was quantified by measuring the total fluorescence at multiple cross-sections of the image. The zero point on the x-axis represents the right edge of microgrooves. All data points were normalized to the first point of “No MLN” as 100 percent. Statistical differences were calculated from four independent experiments using two-way ANOVA. Asterisks show significance compared to Ctrl treatment at a given distance; *, p < 0.05; ***, p < 0.001. (B)P1 cortical neurons were plated on myelin–coated slides and treated with IL-6 and/or HU-210 over a range of concentrations. The neurons were fixed after 24 hrs and labeled for β-III-tubulin. Representative images of neurons plated on myelin (MLN) substrate treated from top to bottom, with either DMSO, IL-6, HU-210 or IL-6&HU-210 labeled with β-III-tubulin show treatment with IL-6 and HU-210 together promotes neurite outgrowth in an inhibitory environment in a dose-dependent manner. (C) Fischer adult rats had unilateral ONC performed followed by intravitreal injections of either 0.5% DMSO for vehicle control (top) or the combination of IL-6 (5 mg/ml) and HU-210 (300 μM, bottom). Intravitreal injections were repeated on day 3 post-crush. After 3 weeks the nerves were removed, sectioned and immunolabeled with GAP-43. Controls had clear crush sites, but no GAP-43 labeled axons crossing the crush site, while IL-6 and HU-210 had regenerating axons nearly 3 mm away as indicated by arrows.

As we had previously shown for Neuro2A cells,^17^ the combination of IL-6 and HU-210 in cortical neurons stimulated STAT3 by phosphorylation at Tyr705 (Supplementary Fig. 1 A-C), consistent with our earlier observation that HU-210-stimulated neurite outgrowth depends on Stat3-mediated transcription. We confirmed the presence of the receptors for IL-6 (gp130) and CB1 (CB1R) in our neuronal cultures by RT-PCR, as shown in Supplementary Fig. 3-I. We also detected the receptors by immunolabeling and found both gp130 and CB1R were expressed in the somal compartment of microfluidic chambers (Supplementary Fig. 3-II). Axotomy significantly increased the number of dead somata *in vitro,* while IL-6 and the combination of IL-6 and HU-210 alleviated cell death (Supplementary Fig. 3-III). These experiments indicate that HU-210 + IL6 applied at the cell body stimulate biosynthetic processes to enable neurite outgrowth.

### IL-6 and HU-210 Treatment for Axonal Regeneration *in vivo*

To evaluate whether treatment with HU-210 and IL-6 promotes axon regeneration following injury *in vivo,* we used the optic nerve crush (ONC) model.^29^ Immediately following surgery and three days after surgery, we treated the injured eye with 5 mg/ ml HU-210 and 300μM IL-6 injected intravitreally, and after two weeks we evaluated for regeneration, first by sectioning the nerve and immunostaining with GAP-43 antibody (Fig. 2C). We observed a modest effect of combined HU-210 and IL-6 treatment. Most of the crushed axons without drug treatment had no detectable GAP-43 labeling even 0.2mm distal to the crush site, while intravitreal injections of a combination of HU-210 + IL-6 resulted in detectable axons nearly 3mm away from the crush site (Fig. 2C bottom image). We also detected and visualized regenerating processes by labeling the axons with cholera toxin-B subunit (CTB) coupled to Alexa-488,^24^ chemically clearing the nerves using the 3-DISCO technique and obtaining high intensity projection images.^26^

### Effects of individual drugs and three drug combinations

We tested whether, when delivered either individually or in pairs, the drugs affected axonal regeneration. Axonal regeneration was not observed in animals that received either no drug treatment or treatment with low doses of intravitreally injected IL-6 or HU-210 (5 mg/ml, 300 μM, respectively) individually, as assessed by CTB-labeled fibers 0.2 mm distal to the crush site (Fig. 3A). The 3-DISCO technique allowed us to image the crush site from proximal to distal end, and to examine the Z-stack to confirm that the crush was complete with no spared fibers. To evaluate the efficiency of the crush method, we performed ONC with no drug treatment and CTB-labeled the nerves immediately afterwards. When we examined the nerve 3 days later, we observed complete crushes in the brightfield setting and the CTB-label was not detected past the crush site (Fig, 3B, bottom panel). Using this technique, we confirmed that IL-6 and HU-210 in combination promoted axonal regeneration, suggesting that **the combination of the two drugs is likely to be synergistic** (Fig. 3C). In nerves from animals that received intravitreal injections of IL-6 & HU-210, we could observe CTB-labeled fibers over 2 mm from the crush site (Fig. 3C2). Fluorescence was quantified at 250-μm intervals from the crush site, revealing a striking difference between the treated and untreated groups (Fig. 3C3). These experiments done in Sprague-Dawley rats showed that the drug effects are not limited to a certain strain of rats.

**Figure 3.**
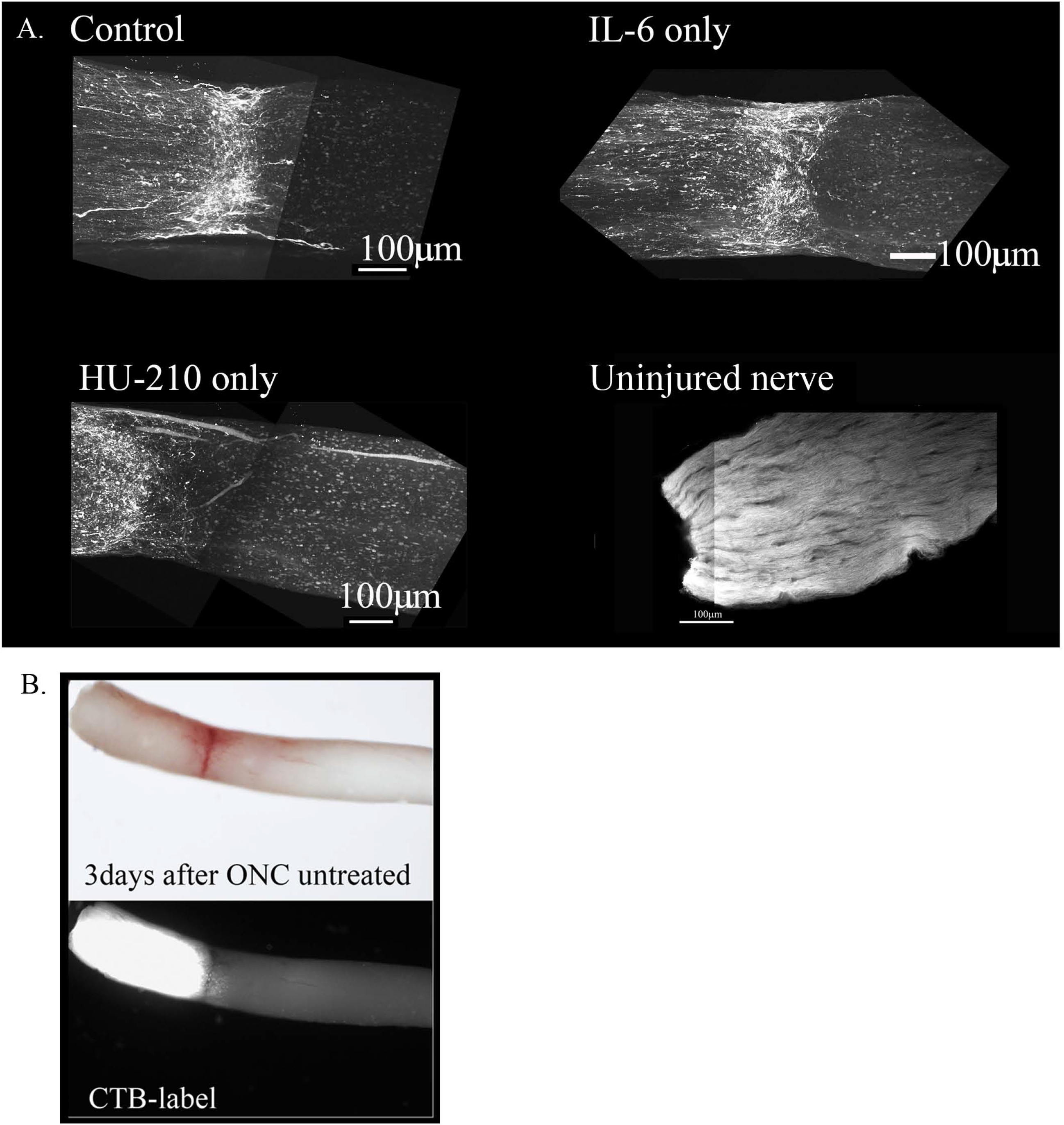
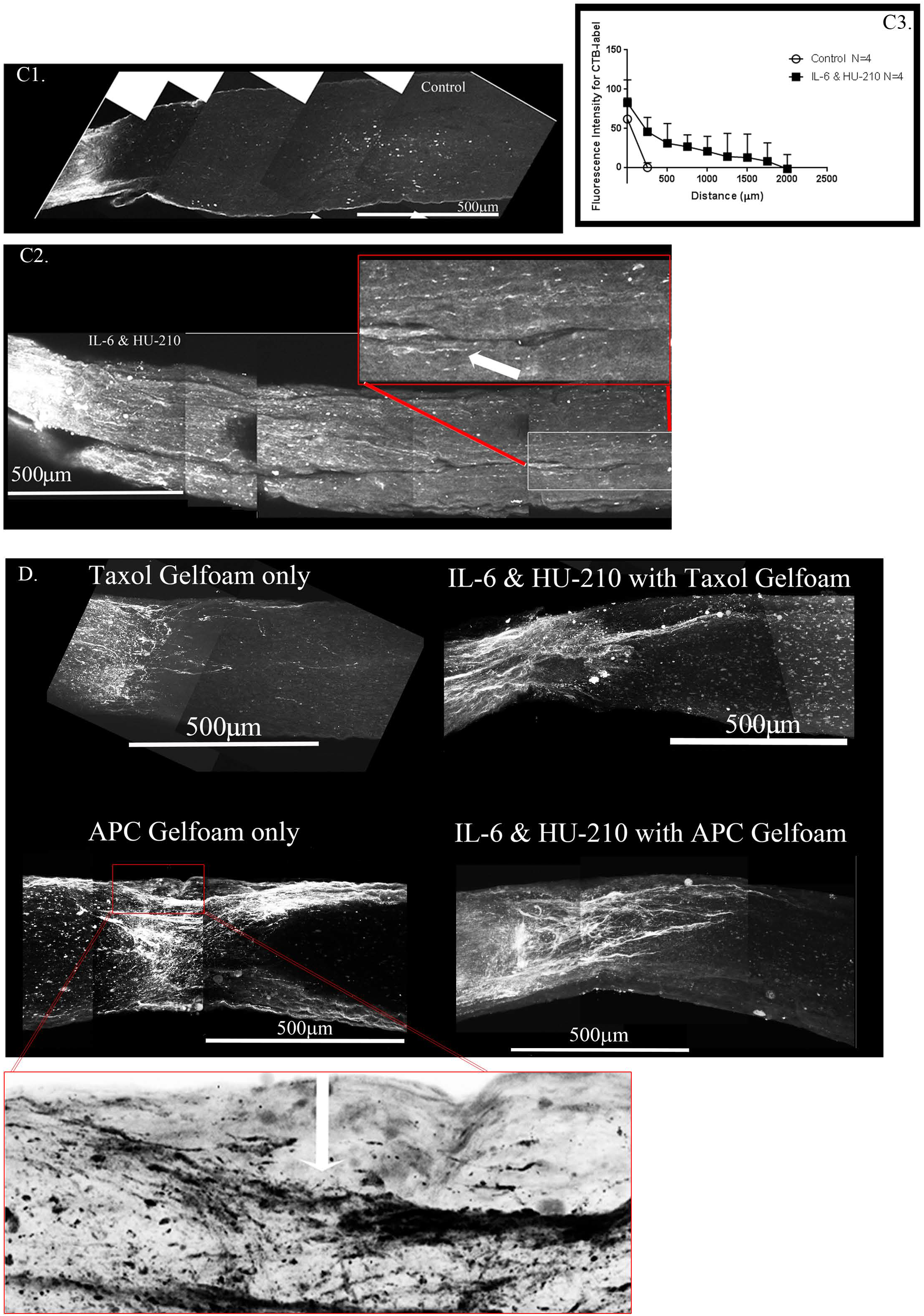
Regeneration after ONC in response to individual or a combination of drugs. (A) Using the rat ONC model and retrogradely labeling regenerating axons with Cholera Toxin B (CTB) by intravitreal injections into the retina 3 days prior to sacrificing and then chemical clearing using the 3-DISCO technique, we were visualizing the whole nerve and not just sections using a Multiphoton microscope. A. Control animals injected with 2.5 μl 0.5% DMSO, have few labeled axons crossing a short distance (less than 0.2mm from the crush site), this was consistent for IL-6 (5 mg/ml) and HU-210 (300 μM) evaluated individually as well, where no significant axonal regeneration was detected past the crush site. Uninjured nerve with CTB labeling and 3-DISCO clearing shows the abundance of axons in the rat optic nerve. (B) Untreated crushed optic nerve was labeled with CTB immediately after injury and 3 days after crush we viewed the nerve under the light microscope (top image), and the CTB-label under fluorescence, showing no spared fibers (bottom image). (C) CTB-labeled regenerating axons within the whole nerve were detected after applying the 3-DISCO clearing technique. Control animals had clear crush sites with few fibers crossing three weeks after injury (C1). However, with IL-6 and HU-210 treatment we could detect CTB-labeled axons within the nerve bundle shown (C2), above the boxed panel is a magnified region demarcated by the red box, showing the extent of regenerating axons over 2 mm away from the crush site, as pointed out by the white arrows. We quantified the extent of axonal regeneration in these chemically cleared nerves labeled with CTB using ImageJ analysis (C3). (D) Taxol applied in gelfoam alone had promoted limited axonal regeneration. Adding Taxol gelfoam with IL-6 and HU-210 injections resulted in more robust growth, but still less than 1mm from the crush site. APC gelfoam alone also promoted modest axonal regeneration from the crush site. Slightly more robust growth is detected when we combine IL-6 and HU-210 with APC gelfoam. The CTB-labeled axons are growing towards the edge where the APC-gelfoam was applied. In the enhanced and inverted image of APC gelfoam demarcated by the red box, we show the crush site that was treated with APC-gelfoam over the injury site. The white arrow in the inverted image of the original emphasizes the crushed fibers that are regenerating, as determined by CTB-labeling.

The morphological observations here and our previous biochemical data indicate that the effects of HU-210 and IL-6, injected intravitreally, targeting the cell bodies reflect upregulated transcriptional and biosynthetic processes and that these agents have the potential to function as therapeutic agents for axonal regeneration *in vivo*.^15,17^ However, in neither Spague-Dawley or Long Evans rats did HU210 and IL6 stimulated axonal regeneration reach the optic chiasm. It has been recognized that biosynthesis alone does not result in significant axonal regeneration after injury. Axonal damage is first induced by neurofilament (microtubule) disassembly, and inflammation can lead to apoptosis and inhibit regeneration.^30,31^ Therefore, we looked for subcellular processes that could be targeted by drugs at the injury site.

Computational models (Supplementary Materials & Supplementary Figures 9-11) showed that the combination of stabilizing growing microtubules and enabling fusion of membrane vesicles at the growth cone under conditions of continuous biosynthesis promoted synergistic growth of axons. Thus, we focused on external inhibitors of axonal growth, including myelin-associated proteins such as Nogo. Biochemical experiments have shown that Nogo receptors and gangliosides mediate glycoprotein inhibition of neurite outgrowth and binding of extracellular agents such as Nogo to gangliosides can promote endocytosis,^32,33^ thus disrupting delivery of new membrane vesicles to the growth cone plasma membrane that is essential for continued growth of the axon. Since the site of injury contains many such myelin-related proteins, to relieve inhibition we decided to test the application of a protease that could act locally. For this we chose APC, which has been reported to promote neuronal repair and axonal regeneration.^23,34^ Taxol is known to stabilize dynamic microtubules, leading to elongation of stable microtubules, and thus promotes axonal regeneration.^19^ We used gel foam to apply 5 μM Taxol or 4.1 mg/kg APC to the site of injury, either alone or in combination with HU-210 (300 μM) and IL-6 (5 mg/ml) applied to the cell body through two intravitreal injection. Taxol alone in gelfoam had a modest effect on promoting some regenerating fibers as reported earlier (Fig.3D).^35^ APC alone promoted significant axonal regeneration about 0.5 mm past the crush site (Fig. 3D), as reported earlier.^34^ The effect of APC was most prominent on one edge of the nerve, presumably where we had placed the drug-impregnated gelfoam (Fig. 3D, bottom panel). In order to ascertain whether these were not spared fibers, we carefully examined the crush site of this nerve treated with APC-highlighted in the red box and an enlarged image where we inverted the image to help reveal the crushed regenerating fibers (Fig. 3D, bottom panel and in the magnified inverted image)- and observed that the crush was in fact complete, with regenerating CTB-labeled fibers growing along the edge of the nerve. Combining IL-6 and HU-210 with Taxol gelfoam (Fig. 3D) produced a more robust response compared to the three agents individually or IL-6 and HU-210 combined, growing over 0.5 mm distal to the crush site. More extensive growth was observed when APC was combined with IL-6 and HU-210 (Fig 3D). However, none of the drugs tested could promote extensive axonal regeneration individually or in three drug combinations, as shown in the summary graph in Fig. 3E.

### Four Drug Combination Treatment for Axonal Regeneration *in vivo*

We tested a four-drug combination with intravitreal injection of HU-210 and IL-6 to deliver these agents to the retinal ganglion cell body, and application of Taxol and APC in gelfoam at the site of injury. We sacrificed the animals after 3 weeks of drug treatment. In a subset of studies, we intravitreally injected CTB one week prior to euthanasia to visualize the regenerated axons. The whole nerve was chemically cleared using the 3-DISCO technique. In separate studies, we labeled the whole nerve with antibodies against GAP-43, a growth cone protein and chemically cleared them using the iDisco technique (Supplementary Figure 4).^36,37^ Both CTB-labeled and GAP-43 immunostained nerves were imaged on a multiphoton microscope. Supplementary Figure 4A shows GAP-43 labeled axons regenerating past the crush site, and Supplementary Figure 4B & C shows axons near and within the optic chiasm, respectively.

A close-up view of the crush site with inverted image colors shows that our crushes are complete with no observable sparing (Fig. 4A, the right panel shows a magnification and color inversion of the area in the red box for clarity). The axons we observe at the crush site and in the chiasm displayed hallmarks of regenerating fibers (Fig. 4A&B). Magnifications B1 and B2 show random processes and bifurcation of axons due to growth without guidance cues. Magnification B3 shows more torturous axon growth. Furthermore, we see CTB-labeled axons in the center of the injured nerve and not just at one edge of the nerve, which would be indicative of spared fibers. Given the remarkable regrowth of the severed axons up to the optic chiasm we decided to look for regeneration into the brain. We sectioned the contralateral brain and observed with immunostaining for CTB that there was label in the superior colliculus, an area innervated by the optic nerve. We also looked for the synaptic marker PSD95 and found it co-localizing with the CTB, indicating the presence of new synapses (Fig 4C).

**Figure 4.**
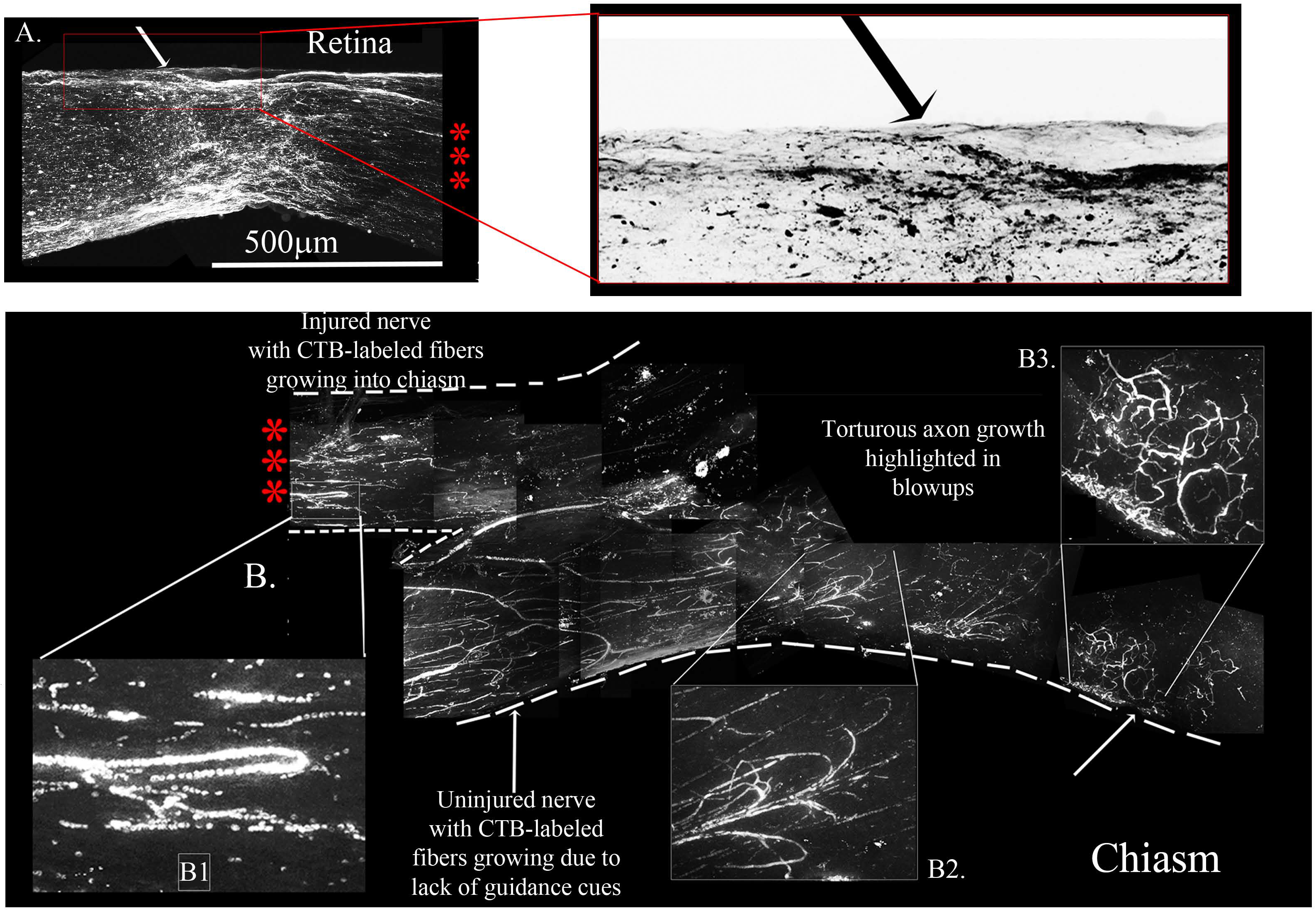
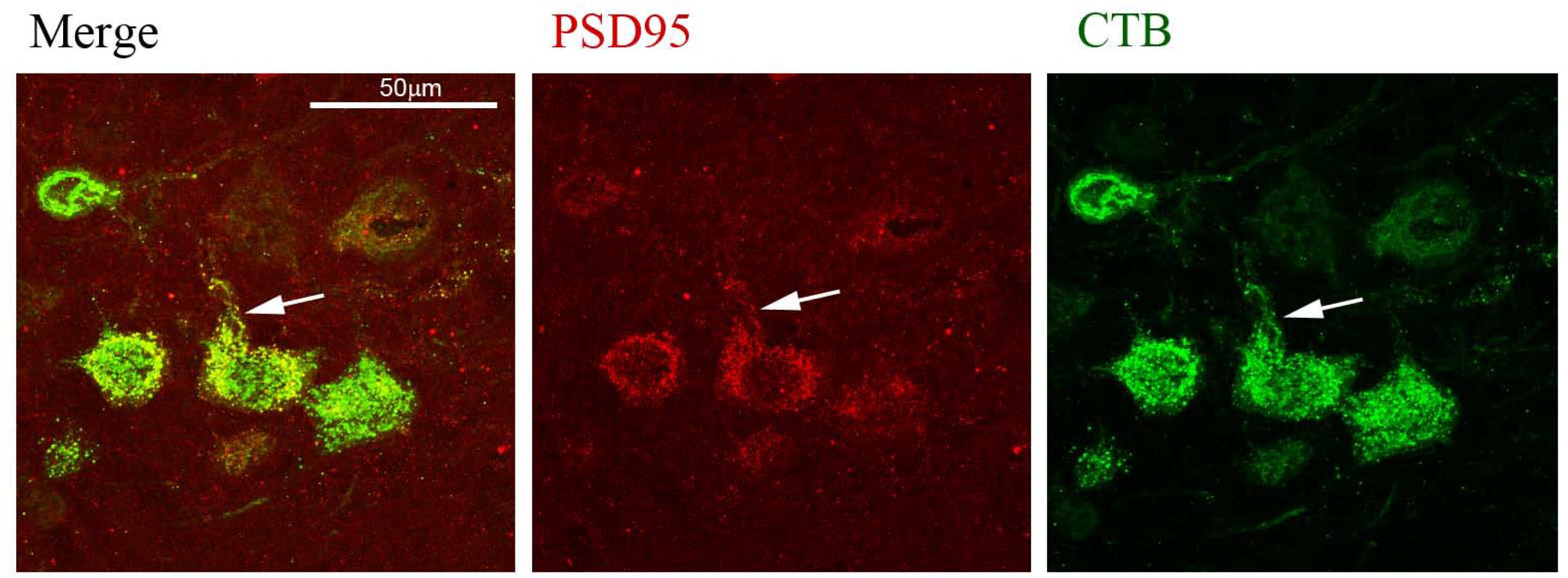
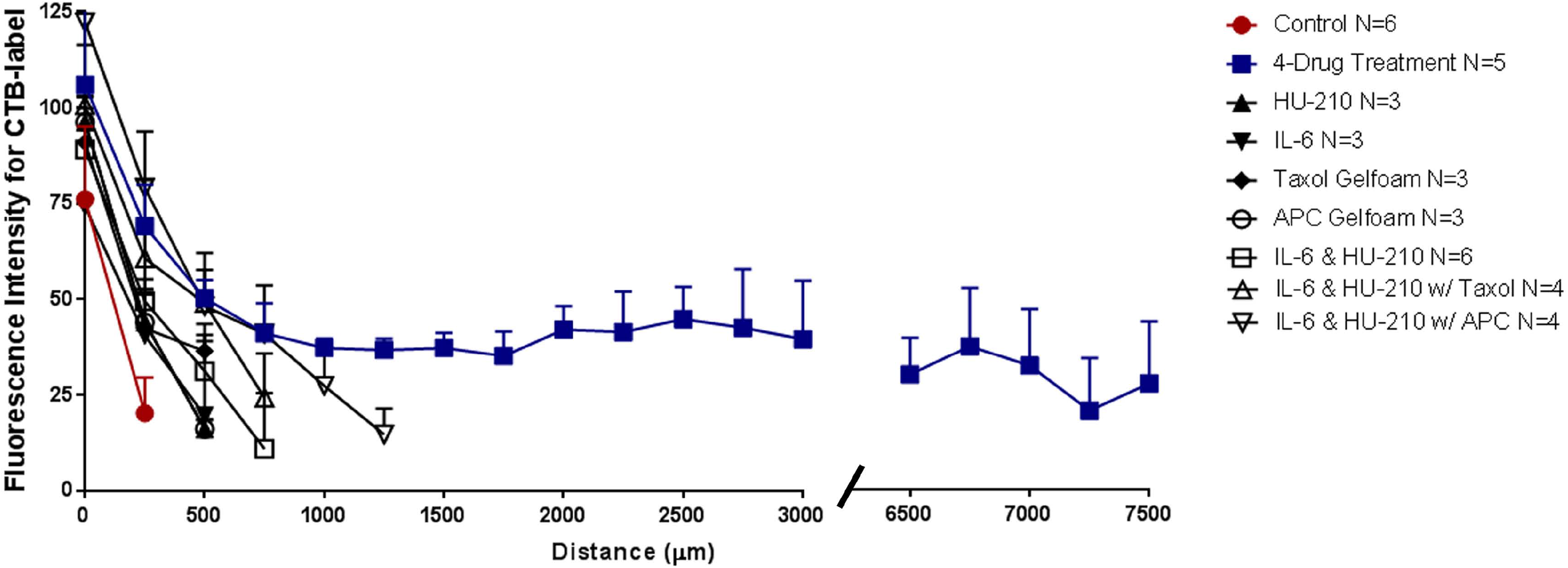
Four-Drug combination in the rat ONC model promotes axonal regeneration to the chiasm. Four-Drug treatment revealing torturous pattern of growth from the Retina (A) chiasm (B) from the same animal, this animal we waited 7days post CTB-labeling. The three red asterisks between the Retina with the crush site (A) to the chiasm (B) represent the same nerve without 6mm of the nerve between, from 0.5mm from the crush site to the chiasm. The red box is enlarged of the crush site in A is inverted to reveal the crushed regenerating axons **C**. Brain sections that are immunostained for CTB revealed CTB labeled neurons and fibers in the superior colliculus. CTB (green) and PSD95 (red) colocalize (yellow) in neurons in the superior colliculus, as highlighted by the white arrows. D. Graph of control, 1-3, and Four-Drug treatments showing long axons only grew in the presence of all four drugs.

With our four-drug combination, we observed that seven out twenty-nine animals tested (5 with CTB-labeling and 2 with GAP-43 immunostaining) had regenerating axons that could be detected at the crush site (Fig. 4A) and continuing to the chiasm, at 7.5 mm from the crush site. (Fig.4B, chiasm, the boxes are magnified regions of the CTB-labeled fibers). Thus, using two separate labeling approaches, we were able to document that regenerated axons reach the chiasm. Four additional animals with the four-drug treatment combination showed axonal regeneration extending to about 3 mm from the crush site (Table 1). A summary of these experiments is shown in Figure 4D. Briefly, eleven out of twenty-nine animals show significant regeneration, with seven of those animals showing axonal growth 7.5 mm past the crush site. ANOVA found a difference among the six groups, *p*<0.0001. Tukey pairwise comparisons between-group differences conserving the overall Type one error at 0.05, i.e., comparing the four-drug treatment to the four other groups measured (control no treatment, APC alone, Taxol alone and IL-6 & HU-210) and concluded that four drug treatment was significant from each other group by *p*<0.0001. Data are represented as mean ± STDEV for optic nerve regeneration.

To determine if the regeneration we saw was real we tested another method of labeling, using AAV8-GFP. The virus was intravitreally injected into rats with ONC and either no treatment or the Four-Drug treatment (Fig. 5). Like the CTB label, we injected the AAV8-GFP virus one week prior to euthanasia. After 3 weeks, the animals were sacrificed, axons were labeled with anti-GFP antibody (abcam, ab13970), and iDISCO was performed to detect GFP label in regenerating axons in the optic nerve of animals with the Four-Drug treatment. We show a robust response to our Four-Drug treatment starting at the crush site in Fig. 5A. We observed tortured growth at the crush site (Fig. 5A1), and by preparing a 3D image of the crush site, we see fibers that are growing towards the center of the nerve (highlighted by asterisk in Fig. 5A2). Together, these observations indicate that the axons are not spared fibers. The growth continues along the optic nerve into Fig. 5B, where we point out GFP labeled axons crossing one another (highlighted by asterisk in Fig. 5B1) and displaying tortured, discontinuous growth in the center of the nerve (Figs. 5B2 & B3). We detected growth into the chiasm, as shown in Fig. 5C. In the magnified area shown in Fig. 5C1, we see torturous axonal growth, including bifurcating axons, which is better displayed in the inverted image in Fig. 5C2. Blue arrows indicate tortured growing axons, and the blue oval indicates the bifurcation, hallmarks of regenerating fibers.^38^ Fig. 5E displays a second nerve labeled with GFP showing long axons also with hallmarks of regenerating fibers. In total, three out of eleven animals labeled with GFP show long, regenerating axons at least 3 mm past the crush site.

**Figure 5.**
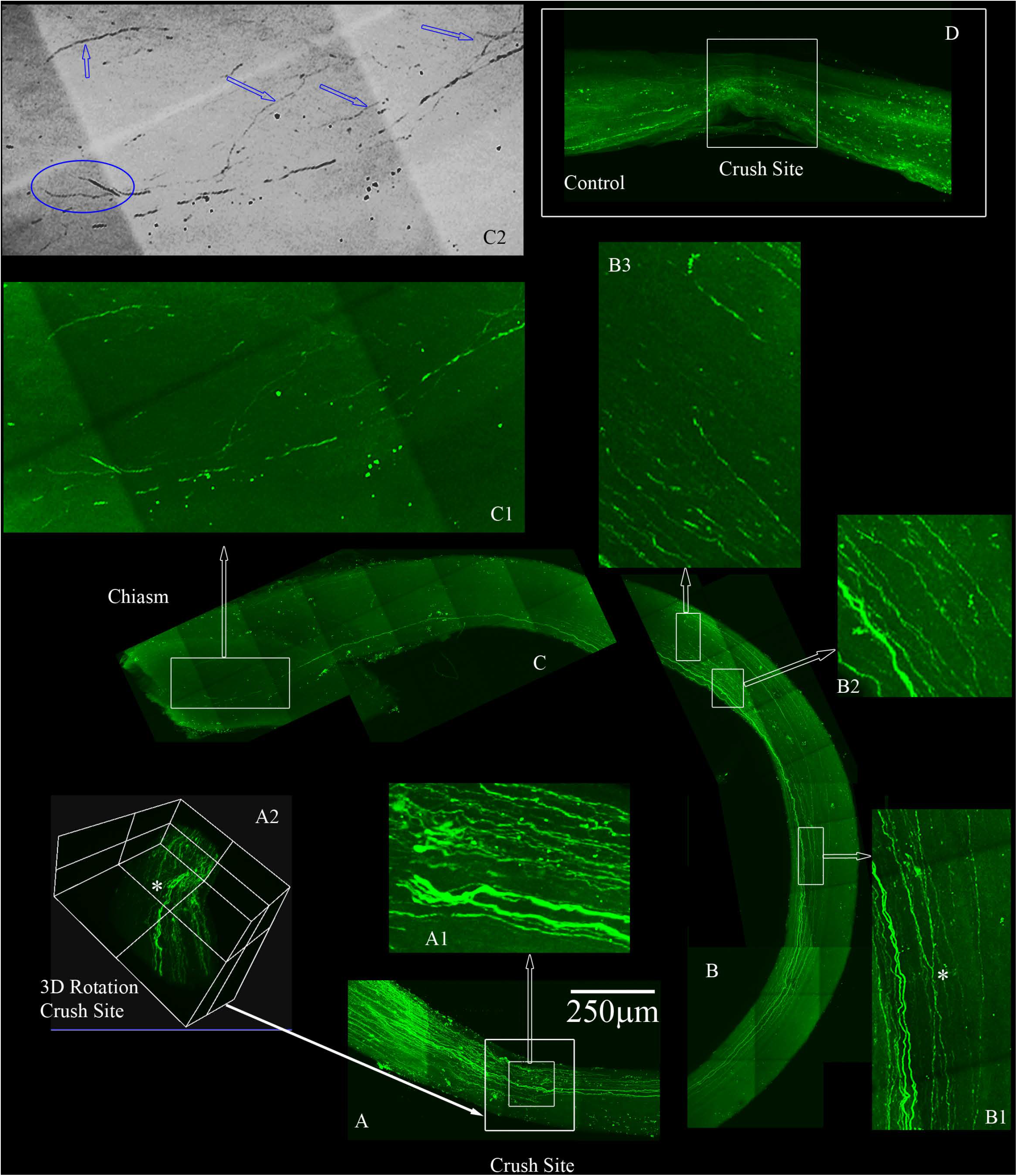
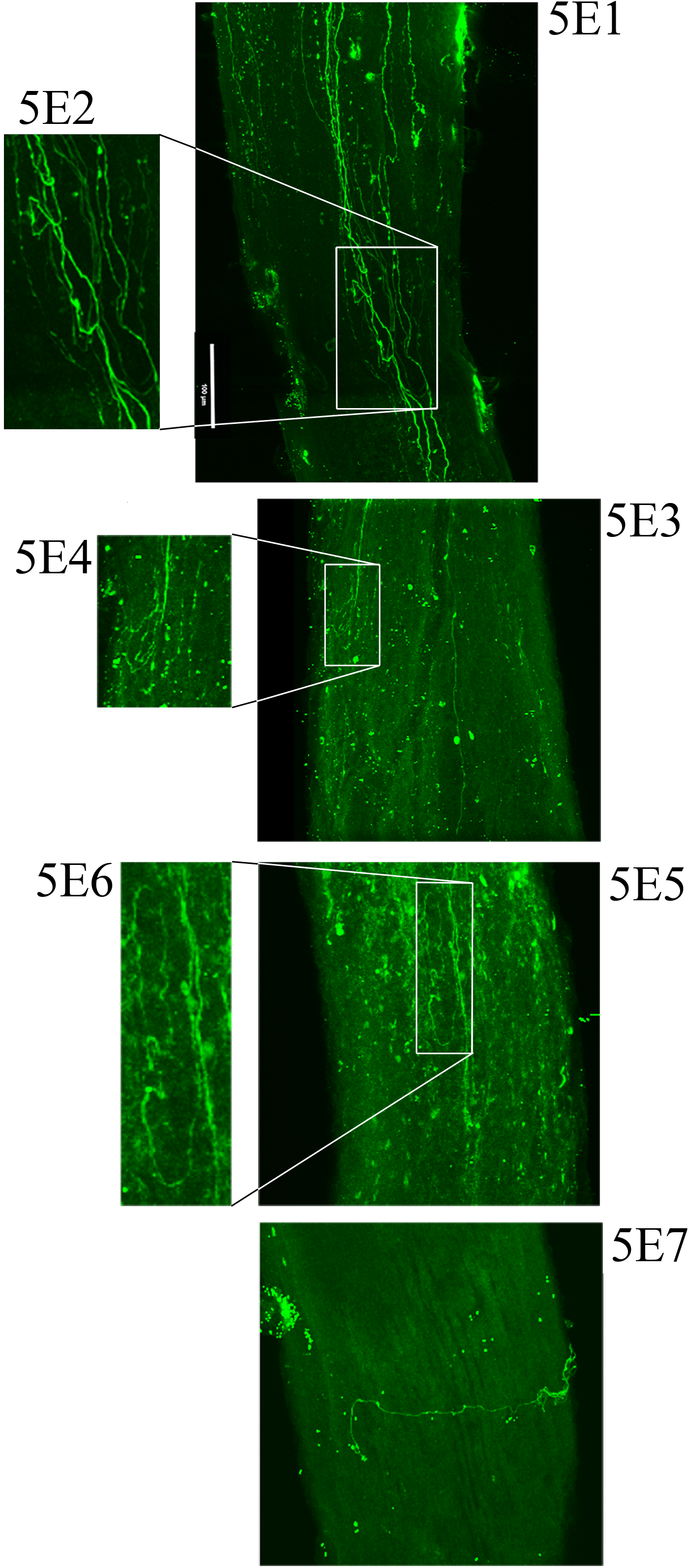
Four-Drug treatment promotes robust axonal regeneration in the rat optic nerve crush model as detected by AAV8-GFP labeling. Adult Sprague-Dawley rats had optic nerve crushes and received gelfoam soaked with APC and Taxol applied over the injury site with IL-6 and HU-210 intra-vitreal injections immediately after the crush. We labeled the regenerating axons with AAV8-GFP injected intra-vitreally to the RGC cell bodies 2 weeks prior to euthanizing. The optic nerves are immunolabeled by iDISCO with anti-GFP and then chemically cleared. **A**. Crush site of the nerve of displaying regenerating fibers expressing GFP. The inner box is magnified in A1, displaying tortured growing axons in the center of the nerve (inner box with traced white arrow). A2, is a 3D rendering of the nerve at the crush site (outer white box in A shown by filled white arrow) that is rotated on its axis to emphasize regenerating axons are not growing on the edges but more in the center of the nerve (see asterisk). **B**. Continuation of the nerve displaying GFP expressing axons that are growing in the center of the nerve and even crossing over in B1 (see asterisk). B2 and B3 are magnified regions with GFP labeled axons over 4mms away from the crush site. **C**. We detect GFP labeled axons into the chiasm of the nerve and the end of the chiasm, we have a magnified region in C1. The image in C1 was converted to grayscale to remove the color and then the image was inverted in C2, to emphasize the tortured growth into the chiasm as demarcated by the blue arrows and the bifurcation (blue circle in C2) of the axons growing into the chiasm. **D**. An example of an adult rat that only received vehicle controls and no GFP labeled axons are displayed past the crush site. All images are taken on an Olympus Multiphoton microscope with a 25X water immersion lens. The images in A1, B1-3 and C1 are taken at 25x with a 1.5x Zoom. Micrometer of 250microns is shown in A.

### The Four-drug combination treatment restores function in animals subjected to optic nerve crush

We tested if the morphologically observed axonal regeneration led to restoration of functional electrophysiological responses. Pattern electroretinograms (pERG) were recorded from both eyes of rats with one crushed optic nerve to determine if the four-drug combination could improve overall retinal electrophysiology, particularly of retinal ganglion cells (RGCs; Fig. 6A).^39–41^ In normal non-injured eyes, pattern electoretinograms (pERGs) elicited by a contrast-reversing checkerboard or sinusoidal grating are characterized by a positive component between 45-60 ms (P1), and a large negative component at 90-100 ms (N2). However, the injured eye of control animals with only vehicle injections had an aberrant pERG, while injured animals that received the four-drug combination treatment showed partial pERG recovery in the injured eye. The recovery approached pre-injury responses for the P50 component. These observations indicate that the four-drug combination stabilizes the cell bodies of the injured retinal ganglion neurons leading to some recovery of electrophysiological function.

**Figure 6.**
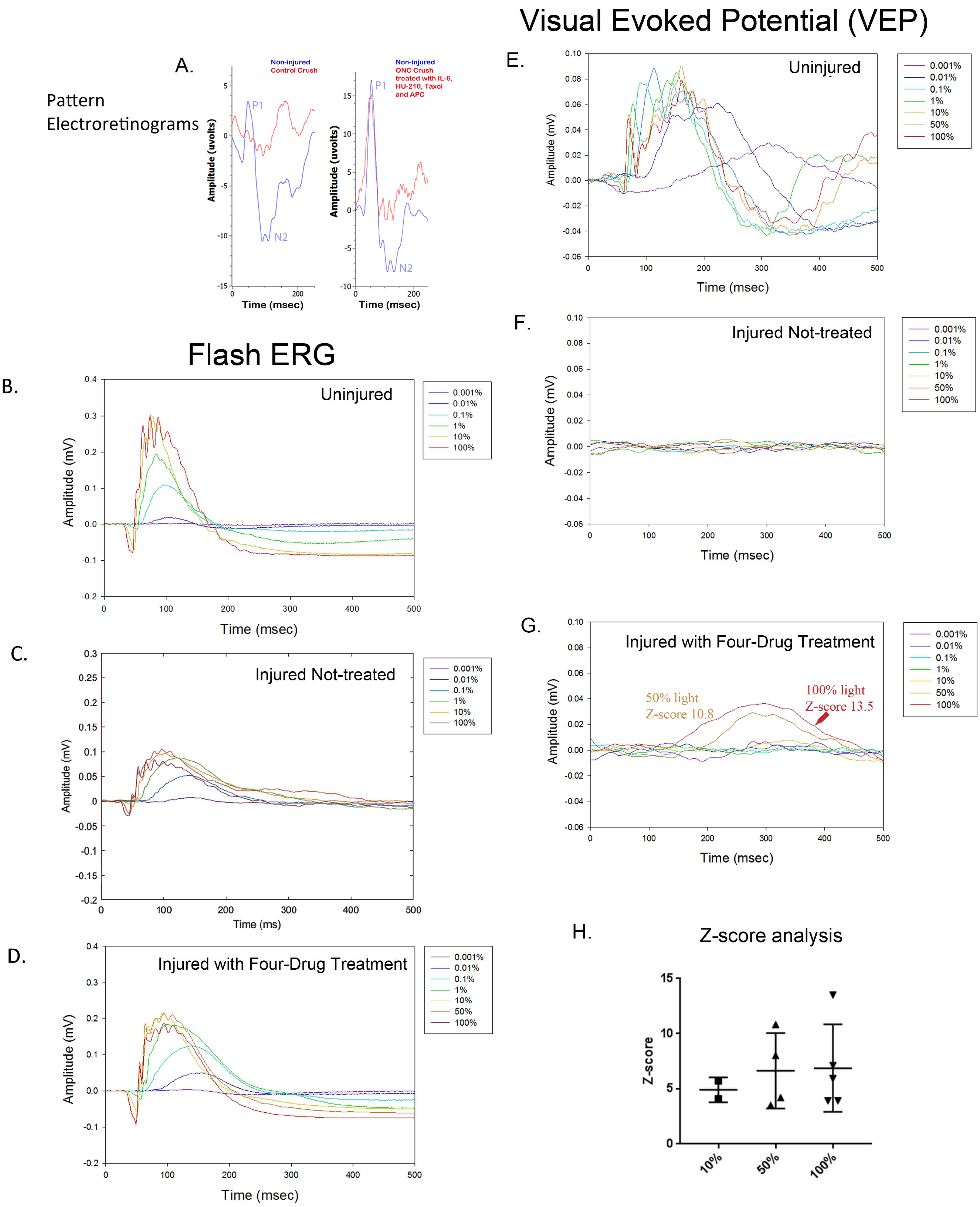
The four-drug combination enhances pattern electroretinograms and VEP responses compared to injured optic nerve treated with vehicle only. (A) We performed pattern electroretinograms (pERG) using the UTAS system (LKC Technologies). Two weeks after optic nerve crushes anesthetized animals had contact lens electrodes placed to record ERGs in response to pattern stimuli. We recorded from both non-injured (blue) and injured (red) eyes with (right panel) or without (left panel) the four-drug combination treatment. Representative recordings of control animals with vehicle only (left panel) showed aberrant pERGs compared to their non-injured eye, while animals with the four-drug combination had pERGs in their injured eye (right panel) approaching baseline levels of their non-injured eye. These are individual raw recordings and not normalized. We performed flash electroretinograms (fERG) which is a strong response to the natural stimuli of light in a graded series of intensities (0.001 to 100% max flash intensity). Representative panels of flash electroretinograms (fERG) traces in response to full light spectrum: B: Uninjured, C: Injured Not-treated, and D: Injured with Four-drug treatment. We simultaneously recorded visual evoked potentials (VEP, E-G) in the brain. Flashes were delivered under dark adapted conditions. In the uninjured eye (E) the VEP was slow for the dimmest flash, peaking around 300ms, and had fast onset with multiple wave components and compound action potentials for the brightest flashes. Injured eyes with no treatment (F) have no detectable VEP activity. The VEP in several treated animals (G) exhibited a slow waveform in response to the highest intensities of light (50 and 100%), suggesting that some axons regenerated all the way to the brain. Z-score values for 50% and 100% are shown in the figure, indicating they are significantly different from the noise. (H) Summary of the Z-scores calculated for the five animals with four-drug treatment that showed VEP signals at some intensity. Of the five animals tested at 10%, 2/5 met the criterion to be above a Z-score of 3; all 4 tested at 50%, showed a response; and all 5 tested at 100% showed a response. The one animal not tested at 50% showed VEPs at 10% and 100%.

pERGs reflect the physiology of retinal neuronal cell bodies, but not of optic nerve axons.^39^ Since we obtained some degree of axonal regeneration into the brain with the four-drug treatment, we determined whether there is concomitant recovery of brain electrophysiological responses to visual stimuli. We stimulated each eye with a series of light flashes at different intensities (0.001 to 100% of 1800cd/m^2^/s) and simultaneously recorded the flash ERG (fERG)-which is indicative of activity in all neuronal cells of the retina (not just the RGCs that innervate the optic nerve)- and the visual evoked potential (VEP) detected bilaterally from the superior colliculus.^42^ Figure 6B-D shows representative fERGs for light flashes to the uninjured eye (Figure 6B), the injured eye of a vehicle-treated animal (Figure 6C), and the injured eye of a drug-treated animal (Figure 6D). fERG signals were altered and often reduced at all intensities for injured eyes with vehicle treatment relative to the uninjured control side, consistent with neurodegeneration that occurs due to the crush. Notably, the photopic negative response, which is attributed to the hyperpolarization of RGCs, is reduced.^43^ The fERGs in animals that received the four-drug treatment, in particular the photopic negative response, appeared nearly normal, indicating that the therapy was neuroprotective for injured retinal neurons.

We then tested if the visual stimulation elicited activity in the brain. For this we recorded VEPs in the visual cortex region of the brain. In Figure 6E, we show the VEP for light flashes presented to the non-injured eye. The VEP is slow and monotonic at lower intensities (<0.1%) and becomes markedly faster and multiphasic at higher intensities. The VEP is completely absent for flashes to the injured eye of vehicle-treated animals (Fig. 6F). In contrast, VEPs were reproducibly detected at high light intensities in several animals that were treated with the four-drug combination (Fig. 6G). At high intensities, the signal was slow and monotonic in these animals, like non-injured eyes at low light intensities. Z-score analysis was performed on each VEP to determine if potentials rose above the levels of noise (further details in Methods section). We found that responses for all the vehicle-treated animals (*n* = 7) were not significantly different from noise, even at the highest flash intensities. However, 5 of 19 drug-treated animals showed VEP responses above the cut-off criterion at different flash intensities (Fig 6H and Table 1B), indicating partial restoration of functional connectivity in those animals. Data of VEP recordings for three individual animals for the non-injured eye, injured eye with vehicle only (DMSO), and injured eye with four-drug treatment are shown in Supplementary Figs. 6-8.

In eleven animals for which electrophysiological measurements were taken, visual function was also assessed using the optokinetic response (OKR). The OKR is a reflexive movement of the head and eyes that results from visual stimulation of the accessory optic system, which is independent of the primary visual pathway.^44,45^ The OKR of each eye can be quantified individually, allowing for the assessment of the crushed optic nerve. Three animals that were treated with the four-drug combination displayed a moderate OKR (Fig 7B). Without the Four-drug treatment no injured animal showed OKR (Fig 7A). A representative video of an animal performing the OKR is shown in the Supplementary Materials. A response in the injured eye upon drug treatment can be observed at 34 seconds.

**Figure 7.**
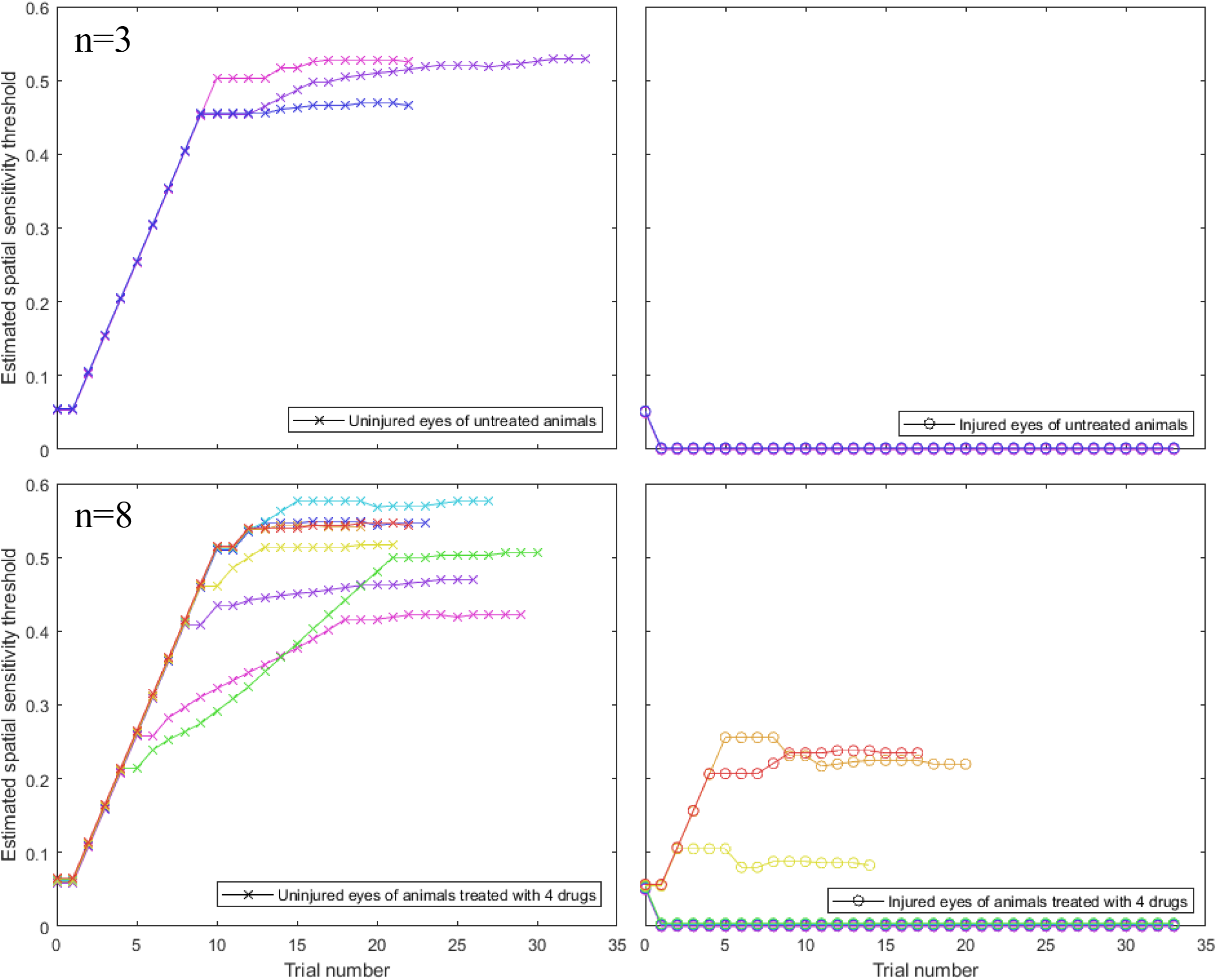
Four-drug combination partially restores visual function compared to injured optic nerves treated with vehicle only. Optokinetic responses (OKRs) of the injured eye (CCW, circles) and uninjured eye (CW, crosses) of animals without and with drug treatment (upper and lower panel, respectively. No responses were observed in the injured eye. Three out of eight of the animals displayed modest responses in their injured eye.

## Discussion

In this study, we designed a drug therapy for a complex pathophysiology by using systems-based logic. We considered the multiple subcellular processes occurring at distinct locations within the cell that could be involved in axonal regeneration and identified four drugs that could regulate these subcellular processes. Three of these four drugs had previously never been tested for axonal regeneration activity *in vivo*. Hence our logic was based on in-vitro experiments, mostly on cultured neurons like cell lines. We focused on drugs that would increase capacity for regeneration at the neuronal cell body and enhance the ability of the axons to grow longer by modulating local subcellular processes in the axonal shaft. We assumed that the process of neurite outgrowth *in vitro* involves many if not all the subcellular processes that neurons use to regenerate axons in vivo. Cell signaling experiments in our laboratory had shown that transcriptional regulation through STAT3 and CREB played a key role in neurite outgrowth.^12,17^ Independent studies had shown a role for the cAMP pathway and SOCS3,^2^ a STAT inhibitor.^9^ Hence, we reasoned that application of receptor ligands that stimulate the STAT3 pathway among other transcriptional regulators could increase the intrinsic capacity of neurons to regenerate.^15^ Since CB-1 and IL-6 receptors are expressed in adult neurons, we selected agonists for these receptors as regenerative drugs. *In vitro* experiments showed that application of these drugs at the cell body was more effective in promoting regeneration, and hence *in vivo* we applied the drug at the cell body as well. As we used the optic nerve crush model, we injected the receptor agonists intravitreally so that they could act on the RGC cell bodies. This resulted in modest but significant drug-stimulated axon regeneration beyond the site of injury. These experiments provided us with an important guideline: that not only should we select the right subcellular process to target, but also that the drug should be applied at the right location. Using this reasoning we selected the two core sets of subcellular processes that function within the growing axon and could targeted by drugs: microtubule growth, and membrane vesicle fusion at the growth cone. Since Taxol stabilizes dynamic microtubules, thus allowing axonal processes to grow, we applied Taxol to the growing axonal regions. By integrating observations in the literature,^32,33^ we hypothesized that the debris at the injury site including myelin-associated proteins might inhibit axonal growth by inhibiting the exocytotic delivery of new membranes to the growth cone, which is needed for axonal growth. Hence, we selected a locally acting protease that could clear the debris field and block these inhibitory agents. Our experiments show that this combination of increasing capacity at the cell body and modulating subcellular processes in the growing axon enables extensive regeneration from the site of injury near the eye to the chiasm. This was a novel effect that we could not have predicted from the *in vitro* studies alone. As axonal injury, via ischemia and reperfusion injury, is implicated in neuronal apoptosis, we anticipated RGC death in our optic nerve crush model. This was confirmed by changes in the ERG following injury, but more importantly we observed rescue of RGCs (and potentially other retinal neurons) following drug intervention, as seen in the repolarization of the pERG and PhNR. We also find that we were able to partially restore electrophysiological communication from the eye to the brain. This appears to have translated into functional recovery of vision-dependent behavior, as animals with partially-restored VEPs also exhibited an OKR. Our results provide an example of the systems therapeutic approach providing a rational basis for the design of combination therapies to treat complex pathophysiologies, in this case nerve damage in the CNS.

Although it is encouraging that we have an identifiable path for a systems logic-based design of rational therapeutics, much further work is needed to translate these findings into viable treatments even in animal models. We have not considered dosing regimens or adverse events associated with this drug therapy. We also need additional strategies to further establish the relationship between the recovery of electrophysiological function and vision-dependent behaviors in the whole animal. These types of experiments will form the basis for future studies. Nevertheless, since observations in the optic nerve crush model are most often applied to spinal cord injury, it will be useful to assess if this four-drug combination can partially reverse spinal cord injury and restore some movement.

## Abbreviations

CB1R: cannabinoid 1 receptor

## Acknowledgements

Multiphoton microscopy was performed in the Microscopy CORE at the Icahn School of Medicine at Mount Sinai and was supported with funding from NIH Shared Instrumentation Grant 1S10RR026639-01. We thank Dr Joseph Goldfarb for a critical reading of the paper and Dr. Alan Weinberg for assistance with the statistical analysis.

