## Supplementary Figures for "Central Nervous System axonal regeneration by spatially targeted drug combinations"

Supplementary Figure 1

A.

B.

C.

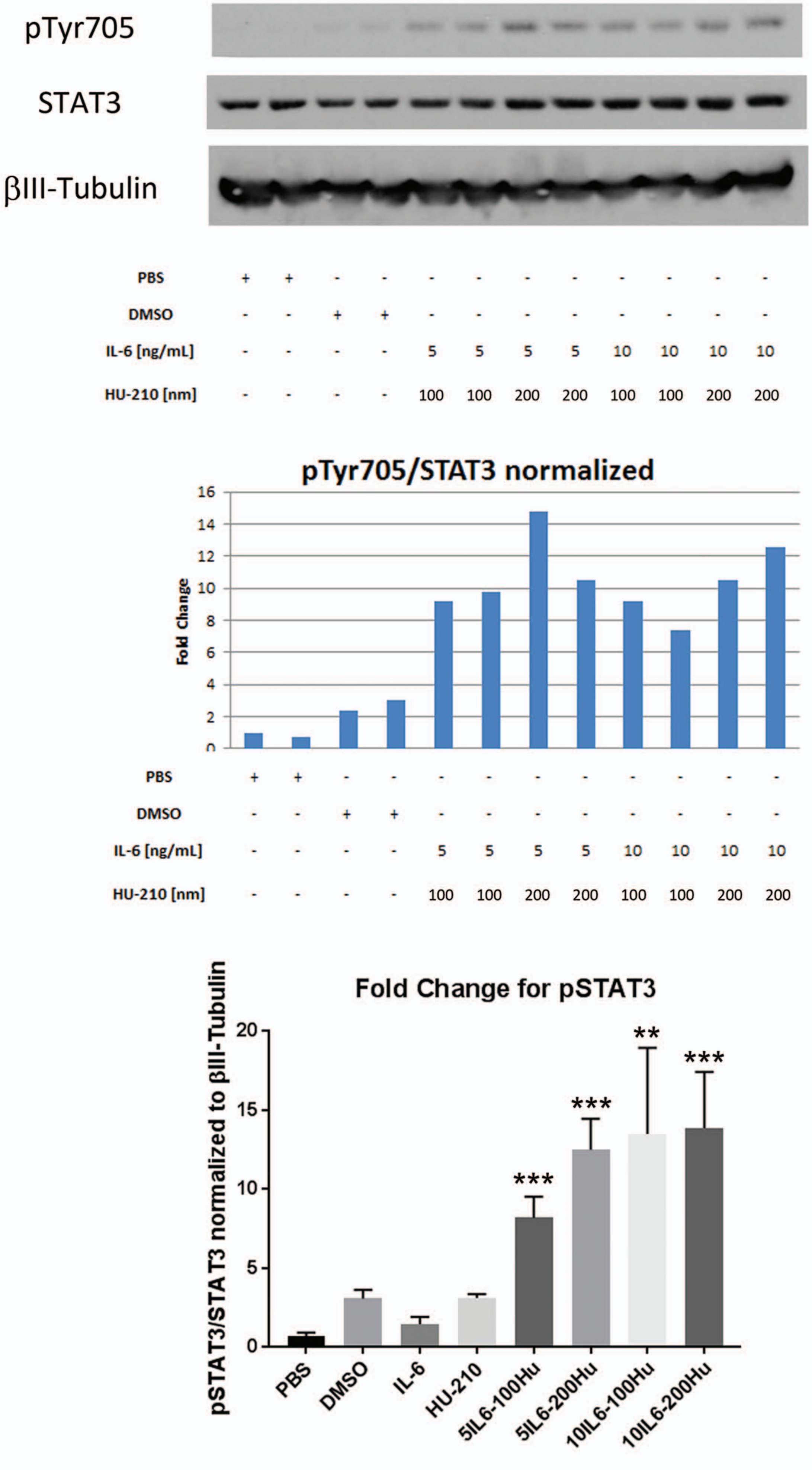

**Figure S1: Combination of IL-6 and HU-210 activates STAT-3.** (A) Cortical neurons in non-supplemented neurbasal media were treated with IL-6 and HU-210 or controls as described for 2 hrs. The cells were lysed and subjected to immunoblot analysis ofr phospho-STAT3 (pTyr705), total STAT3 and βIII-tubulin. (B) We normalized the samples for phospho-STAT3 to total STAT3, and fold changes are displayed. (C) In three independent experiments, we averaged phospho-STAT3 to total STAT3 normalized to βIII-tubulin. For controls we used PBS and 0.5% DMSO. For the treated samples we used 5 ng/ml IL-6 and 100 nM HU-210 (5IL6-100Hu), 5 ng/ml IL-6 and 200 nM HU-210 (5IL6-200Hu), 10 ng/ml IL-6 and 100 nM HU-210 (10IL6-100Hu) and 10 ng/ml and 200 nM HU-210 (10IL6-200Hu). All statistics are unpaired t-test, $^{**}p<0.01$, $^{***}p<0.001$.

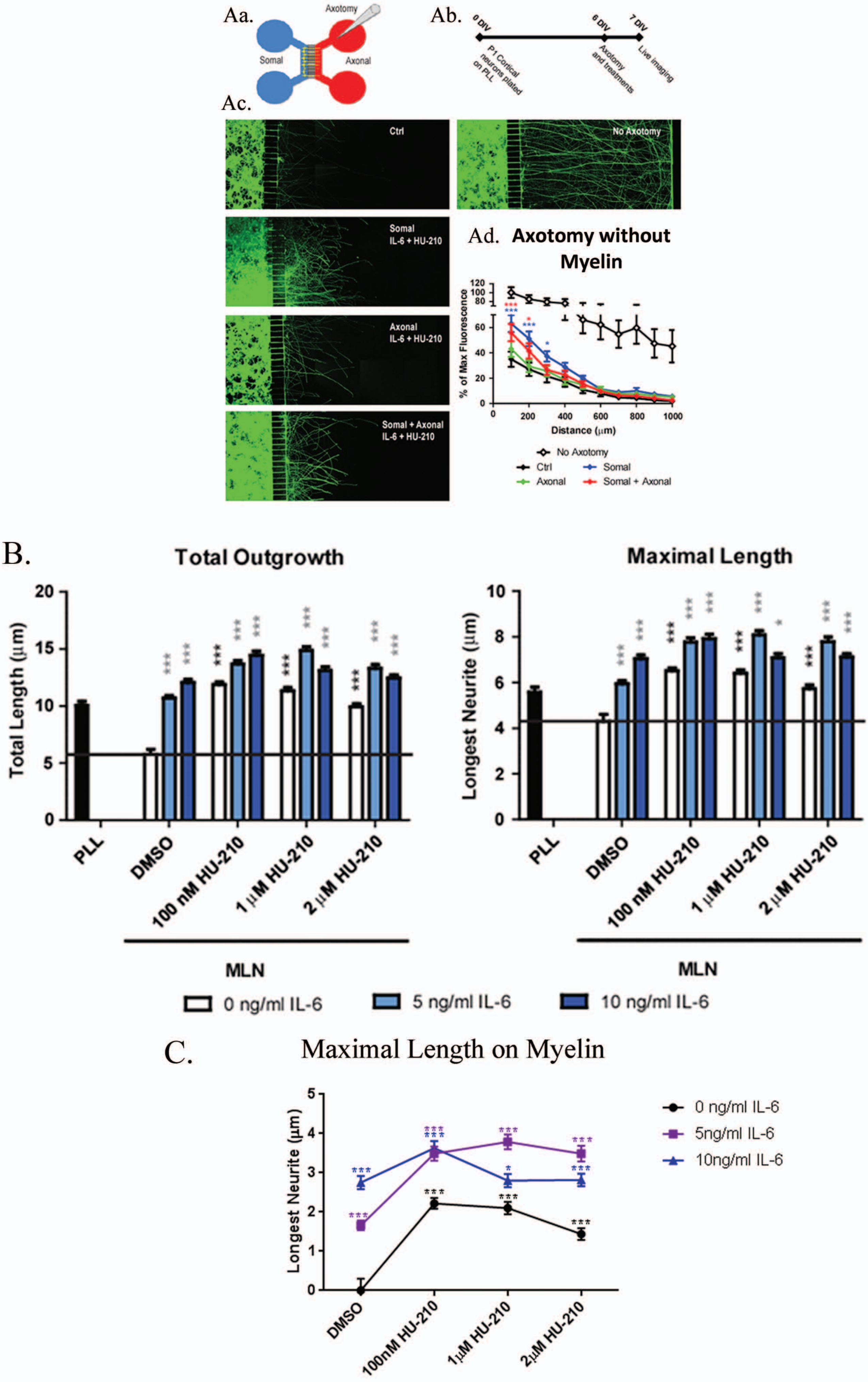

Aa.
Axotomy
Somal
Axonal
Ab.
0 DIV
6 DIV
7 DIV
P1 Cortical neurons plated on PLL
Axotomy and treatments
Live imaging
Ac.
Ctrl
No Axotomy
Somal IL-6 + HU-210
Axonal IL-6 + HU-210
Somal + Axonal IL-6 + HU-210
Ad. Axotomy without Myelin
% of Max Fluorescence
Distance (μm)
No Axotomy
Ctrl
Somal
Axonal
Somal + Axonal
B.
Total Outgrowth
Total Length (μm)
Maximal Length
Longest Neurite (μm)
PLL
DMSO
100 nM HU-210
1 μM HU-210
2 μM HU-210
MLN
0 ng/ml IL-6
5 ng/ml IL-6
10 ng/ml IL-6
C.
Maximal Length on Myelin
Longest Neurite (μm)
DMSO
100nM HU-210
1μM HU-210
2μM HU-210
0 ng/ml IL-6
5ng/ml IL-6
10ng/ml IL-6

**Figure S2A. Somal application of IL-6 and HU-210 results in enhanced neurite outgrowth after axotomy.** Cortical neurons were plated in the somal compartment (blue) of microfluidic chambers, schematically represented in (Aa.) and allowed to grow axons across the microgrooves into the axonal compartment (red). The gray cone depicts a pipet used to perform axotomy by aspirating medium from the axonal compartment. (Ab.) For axotomy experiments, neurons were cultured in chambers on PLL for 6-7 days, and axotomy was performed by aspirating all medium from the axonal compartments 2-4 times. 5 ng/ml IL-6 and 100 nM HU-210 were added together to the somal, axonal or both compartments after axotomy. Treatments were added to the indicated compartments in neurobasal (NB) immediately after axotomy, and neurons were imaged live after 24 hrs using Calcein AM. (Ac.) Representative confocal images after axotomy and treatment. (Ad.) Neurite outgrowth was quantified by measuring the total fluorescence at multiple cross-sections of the image. The zero point on the x-axis represents the right edge of microgrooves. All data points were normalized to the first point of "No Axotomy" at 100 μm from the microgrooves as 100 percent. Statistical differences were calculated from two independent experiments using two-way ANOVA. Asterisks show significance compared to Ctrl treatment at a given distance; *, $p < 0.05$; ***, $p < 0.001$.

**Figures S2B & C. Stimulation of cortical neurons with IL-6 and HU-210 together promotes neurite outgrowth on myelin in a dose-dependent manner.** P1 cortical neurons were plated on myelin–coated slides and treated with IL-6 and/or HU-210 over a range of concentrations. The neurons were fixed after 24 hrs and labeled for β-III-tubulin. Quantifications of neurite outgrowth were performed from 1500-2000 cells per condition. The bar graphs show mean ± S.E.M. of the longest neurite of each cell on myelin. Statistical differences were calculated using a one-way ANOVA followed by Bonferroni's multiple comparison test (B). We also made a line graph of the data for the Maximal Length, black asterisks show significance between the treatment and DMSO alone; blue or purple asterisks show significance between each combinational treatment and the sole HU-210 treatment at the corresponding concentration; *, $p<0.05$; ***, $p<0.001$ (C).

### Supplementary Figure 3

I.

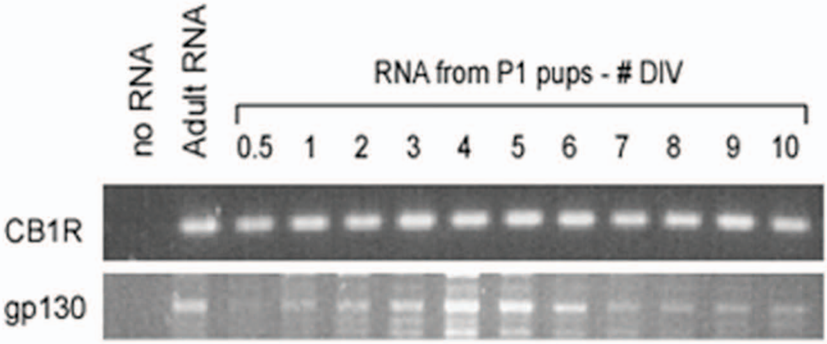


II.

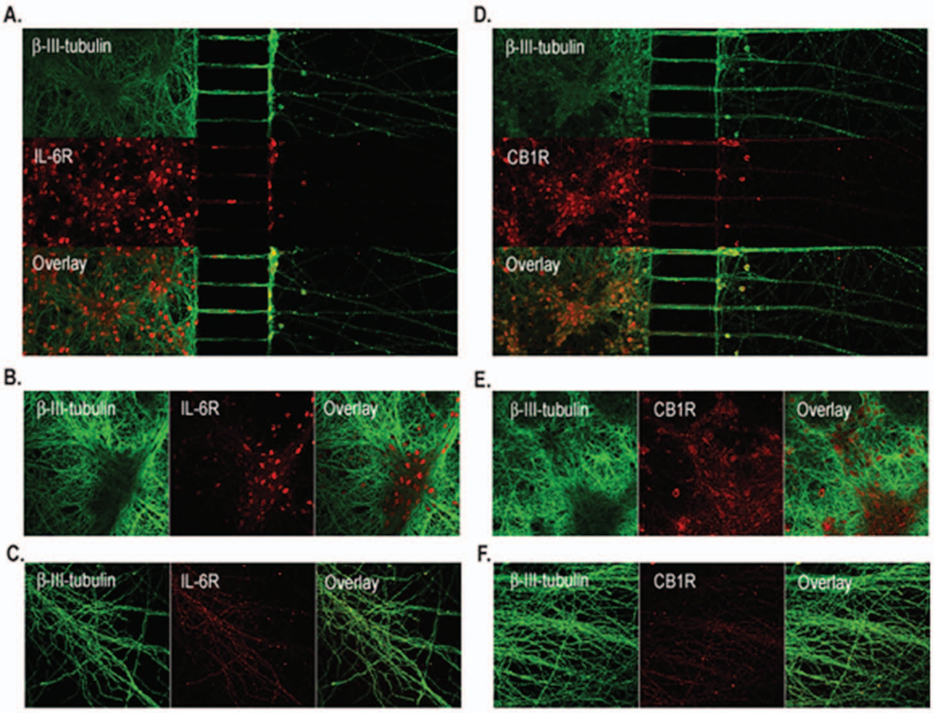


III.

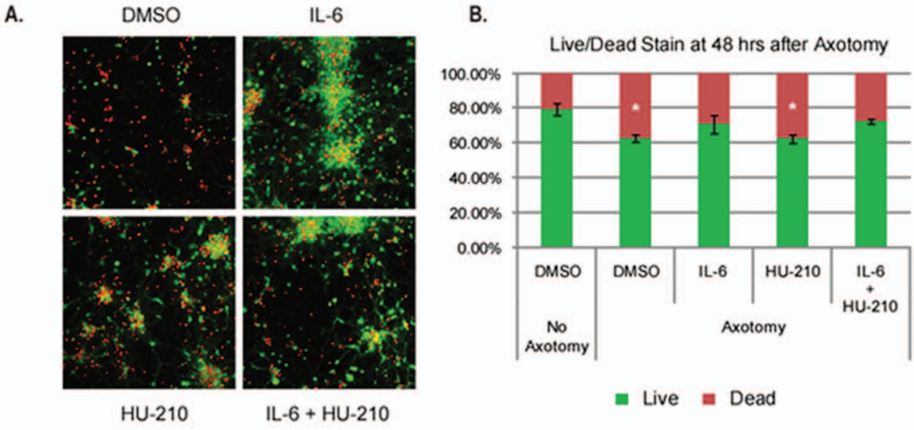

**Figure S3-I: CB1R and IL-6R expression in rat cortical neurons.** P1 cortical neurons were plated on poly-L-lysine-coated dishes for the indicated number of days *in vitro*, and mRNA was isolated and subjected to RT-PCR for CB1R or gp130.

**Figure S3-II: IL-6R and CB1R are expressed strongly in the soma and weakly in the axons of cortical neurons.** Cortical neurons were plated in microfluidic chambers and allowed to grow axons across the microgrooves for 6-7 days. Neurons were fixed and immunolabeled for β-III-tubulin (green) and (A-C) IL-6R (red) or (D-F) CB1R (red), to determine the location of receptors. (A, D) Tiled composite images of neurons in microfluidic chambers. (B, E) Representative fields of neurons in the somal compartments. (C, F) Representative fields of neurons in the axonal compartments.

**Figure S3-III: Somal treatment with IL-6 promotes neuronal survival after axotomy.** Neurons were grown in chambers for 7 days, starved in NBS for 1-2 h, subjected to axotomy and treated for 48 h. Calcein AM (green) and ethidium homodimer-1 (red) were applied to the somal compartment and live imaging was performed. Images were taken from regions immediately to the left of the microgrooves to ensure imaging neurons that were previously axotomized. (A) Representative confocal images after axotomy and treatment. (B) Quantification of total green to red fluorescence ratio was performed by measuring number of pixels of each color. Statistical differences from two independent experiments were calculated using a one-way ANOVA followed by Bonferroni's multiple comparison test. Asterisks show significance in comparison to "No Axotomy"; *, $p < 0.05$.

### Supplementary Figure 4 - GAP-43 i-DISCO

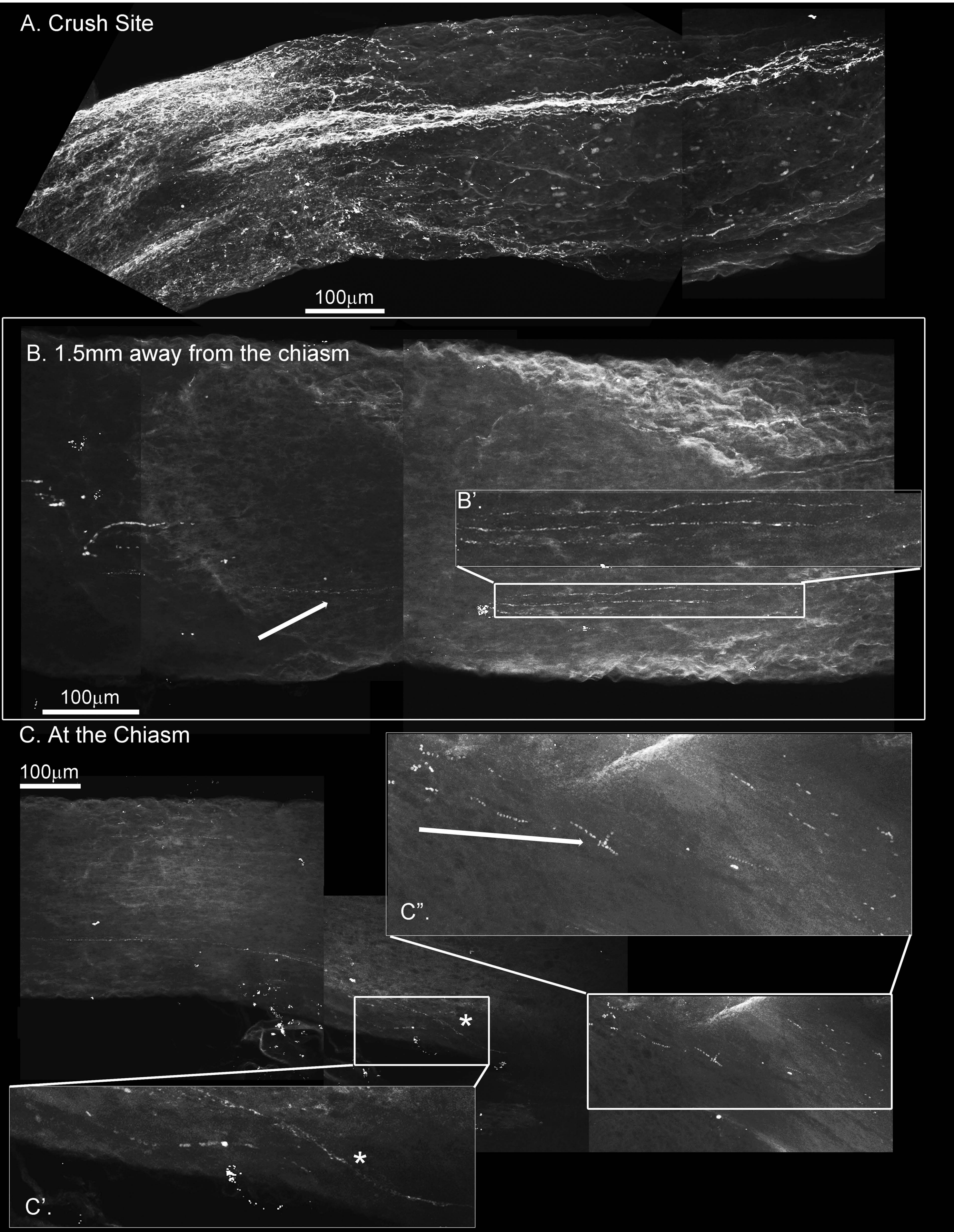

**Figure S4: GAP-43 immunostaining using the i-DISCO method to stain and chemically clear the entire optic nerve that was crushed and 4-drug treated reveals GAP-43 staining in the chiasm.** ONC with 4-drug treatment and i-DISCO GAP-43 staining reveals the crush site with regenerating fibers (A.). Approximately 6mms away from the crush, near the chiasm we also detect GAP-43 positive axons (B.) the blowup displays several of these GAP-43+ axons. We even detect fibers in the chiasm (C.) as revealed by the blowups in C' and C".

### Supplementary Figure 5

4-drug treatment individual slices displaying torturous pattern of axonal regeneration

A. Z-Slice #240 out of 537 slices

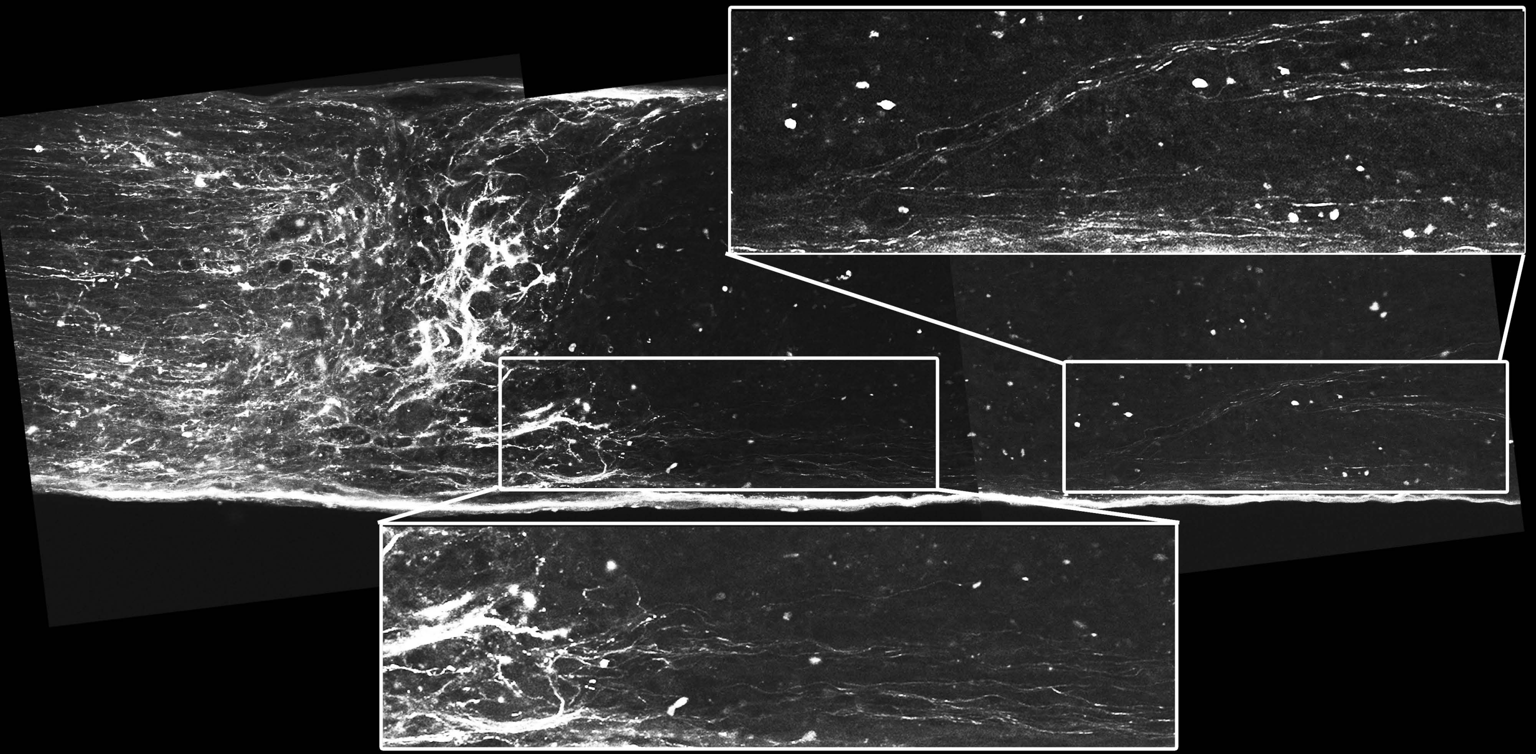

B. Z-Slice #274 out of 537 slices

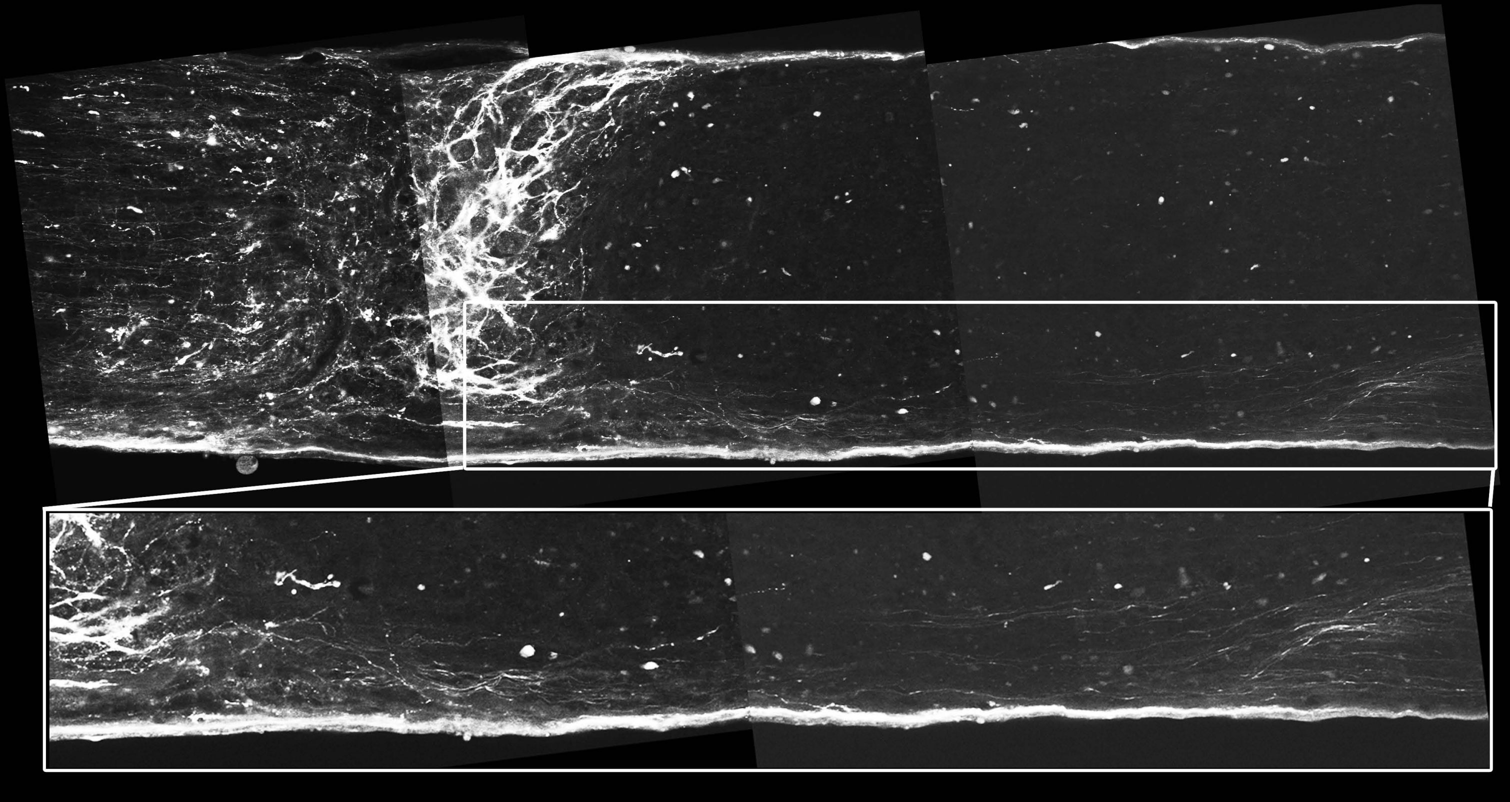

**Figure S5: Individual Z-slices of ONC with 4-drug treatment reveal a torturous pattern of axonal regeneration.** The image in Figures 2-II & III are a 2-D projection after 3-DISCO clearing of nearly 540 Z-slices, to reveal the pattern of CTB-labeled axons within the whole nerve bundle. Looking at individual Z-slices of this same samples, towards the center of the nerve we look at section #240 out of 537 (A.) and #274 out of 537 (B.). These individual slices or planes of the nerve reveal the CTB-labeled crush site and torturous growth at the crush site and about 1mm beyond that. These two images are just two slices out of the 537, that are used to generate the images in Figures 2-II & III.

### Supplementary Figure 6

#### UNINJURED VEP

I.

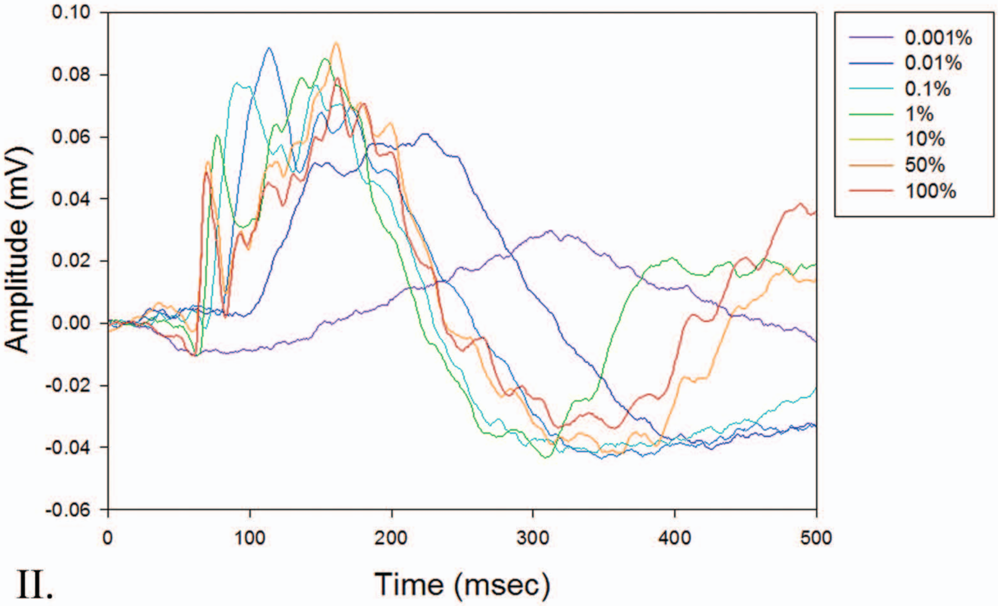


II.

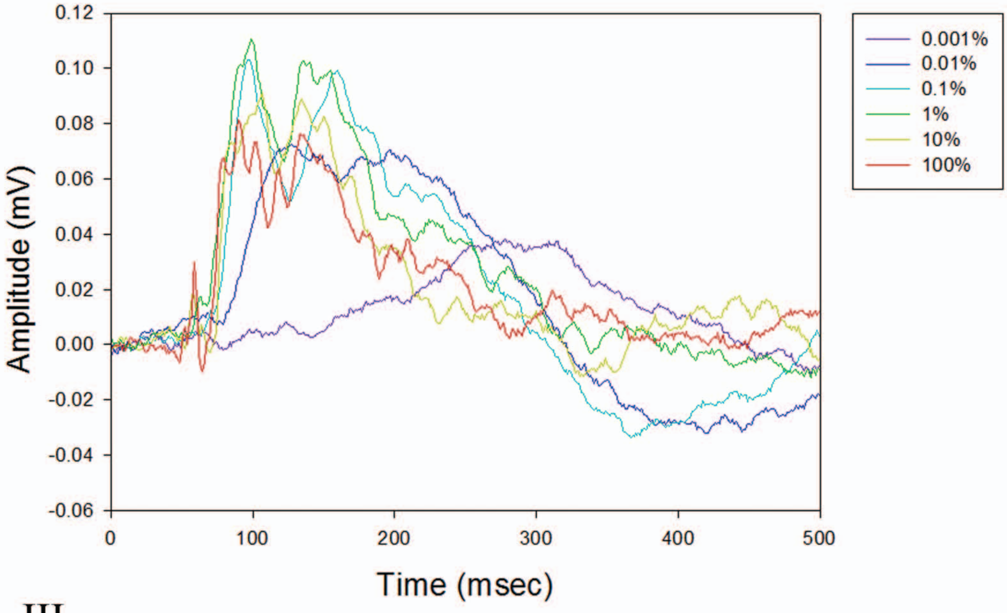


III.

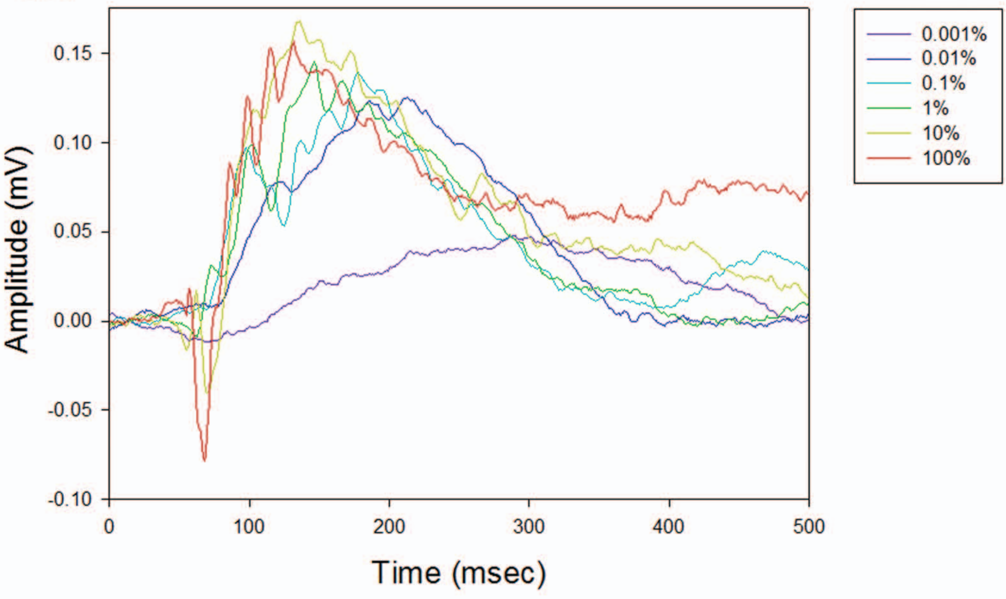

### Supplementary Figure 7

#### INJURED NOT-TREATED

I.

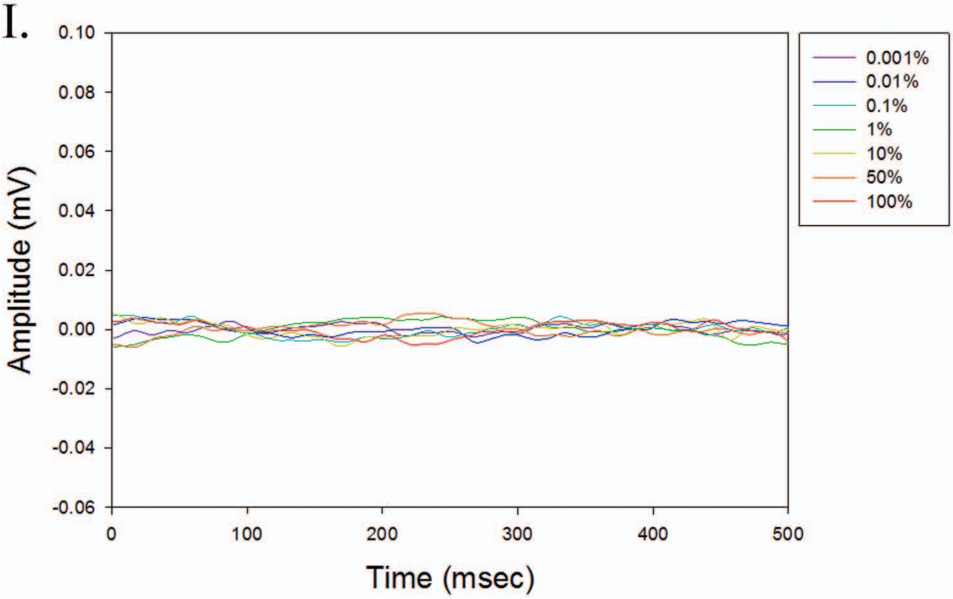


II.

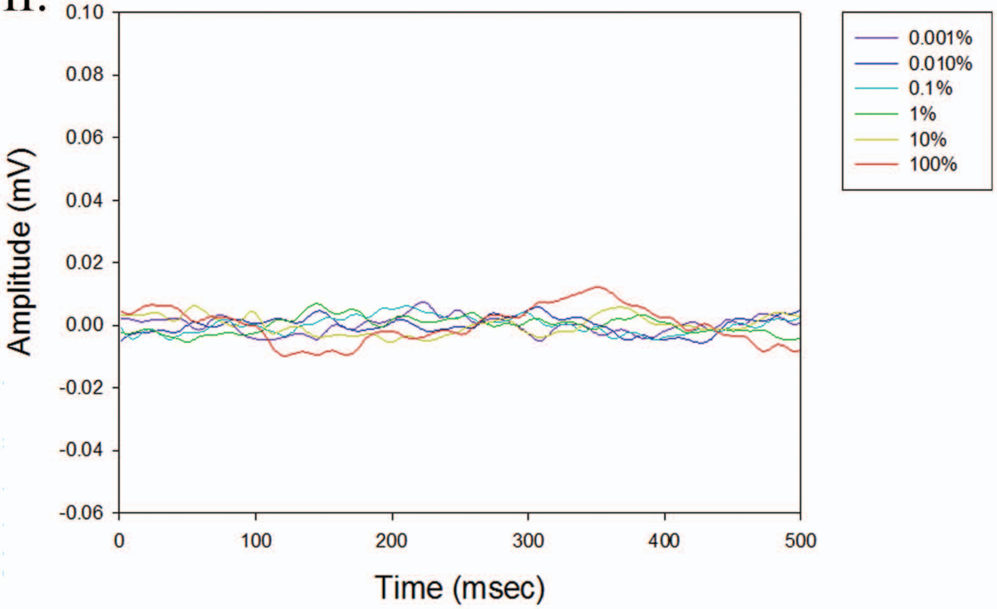


III.

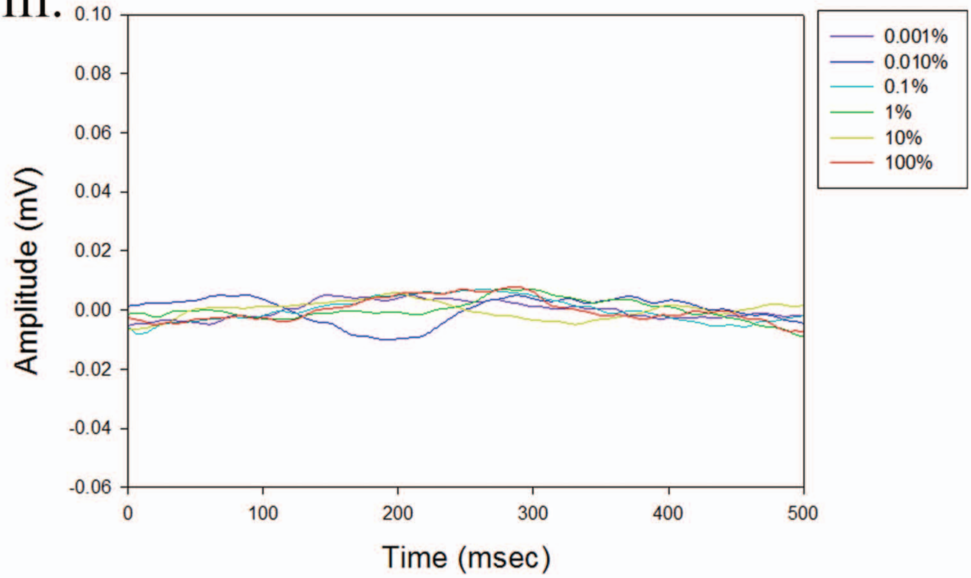

### Supplementary Figure 8

#### INJURED WITH FOUR-DRUG TREATMENT

I.

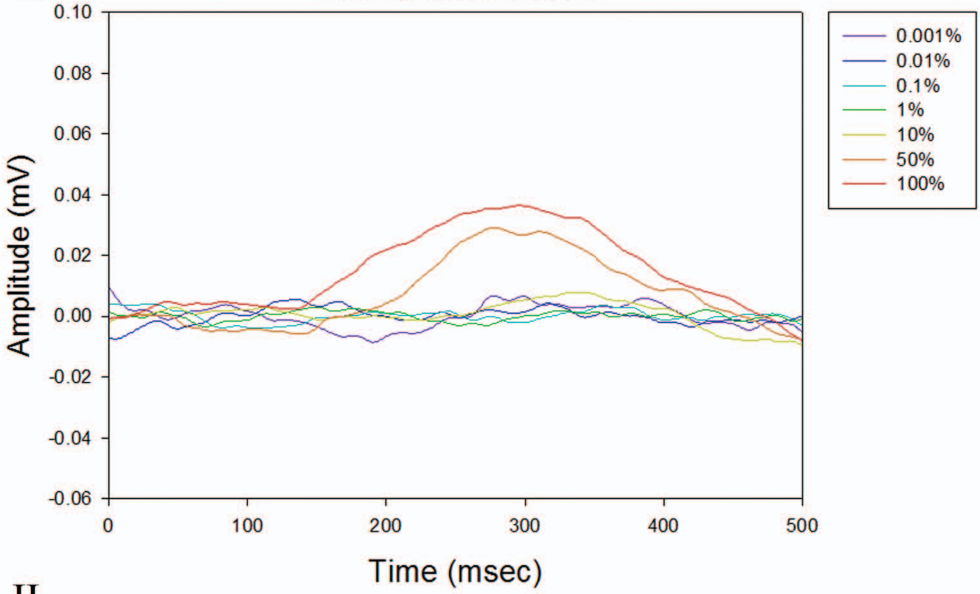


II.

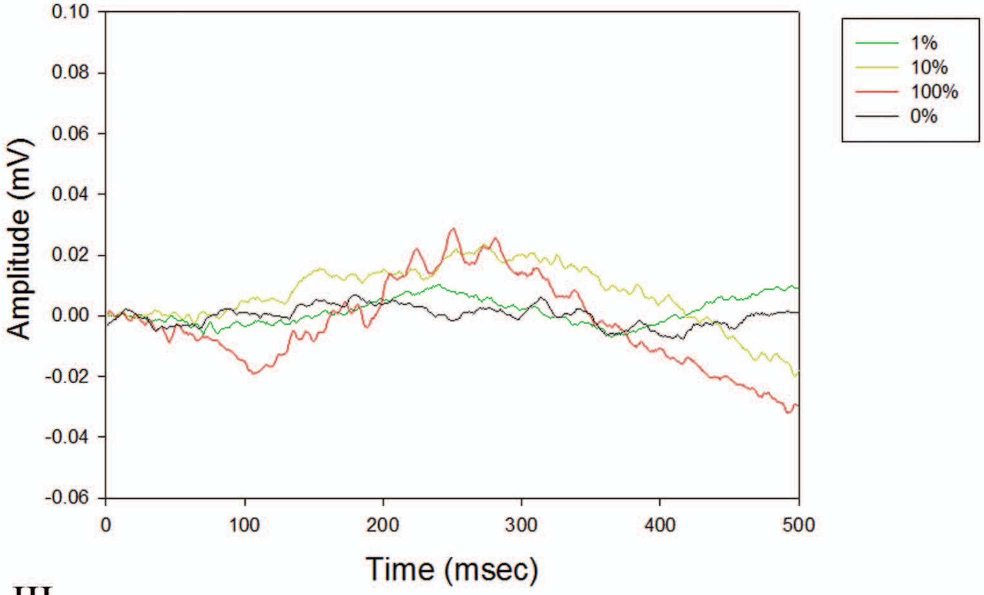


III.

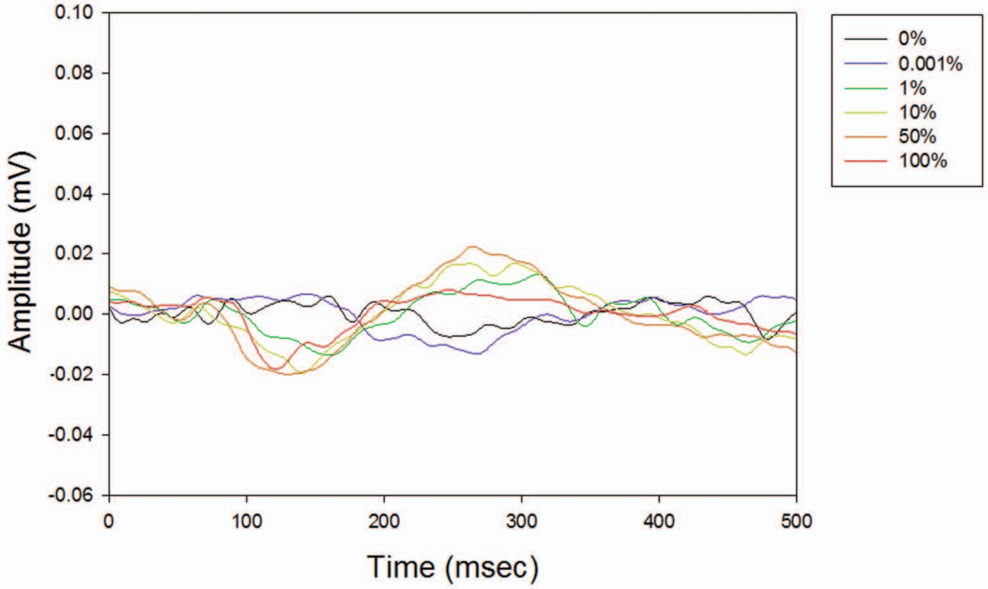

**Figure S6: Three representative VEP responses in uninjured eyes.** Three examples of uninjured nerves VEP responses to full-light intensity series.

**Figure S7: Three representative VEP responses in injured eyes not treated.** Three examples of injured nerves with no treatment VEP responses to full-light intensity series.

**Figure S8: 4-drug treatment partially restores VEP responses.** Three examples of injured nerves with four-drug treatment VEP responses to full-light intensity series.

Supplementary Figures 9-11 Computational model of Drug Action

The extensive regeneration obtained by the four-drug combination raises the question of how might the drug combination observed effects. For this we used the top down version of the neurite-outgrowth multicompartment ODE model we have developed (18). This model analyzes the dynamics of a whole cell process in terms of interactions between three sub cellular processes *Production of Membrane Components, Vesicle Transport and Exocytosis* and *Microtubule Growth.* This model is built on the microtubule growth model developed by Margolin et al. (31) with the vesicle transport model by Heinrich and Rapaport (32). We used this model of neurite outgrowth as a surrogate for regeneration of the injured axon.

**Basic model**

This model is an extension of the previous model by Yadaw et al (18). A key difference between this model and the previous one is the number of compartments. Here we use six compartments as this needed to account for the action of drugs. The six compartment include two terminal compartments Trans Golgi Network (TGN), growth cone plasma membrane (GC-PM) and four intermediate compartments (cell body cytoplasm (CBC), neurite shaft cytoplasm proximal (NSC-P), neurite shaft cytoplasm distal (NSC-D), growth cone plasma membrane (GCC). The dynamics of the model starts with production of membrane components at TGN and followed by coat protein B mediated vesicles budding from TGN into cell body cytoplasm, coat protein B containing vesicles are anterograde moving vesicles which are actively transported from cell body cytoplasm to growth cone cytoplasm by the kinesin motor protein. Retrograde (endocytosed) vesicles movement process starts with coat protein A containing vesicles budding at growth cone plasma membrane and it is actively transported to TGN by dynein motor protein. When anterograde and retrograde vesicles reach to target compartments GC-PM and TGN respectively then vesicles tether to the target compartments by interactions between vesicle-SNARE and target-SNAREs and this is PLOS followed by fusion with target compartment. Kinesin mediated anterograde moving vesicles have different binding rates in intermediate compartments because the microtubule associated protein tau is distributed as a gradient increasing from CBC to GCC where Kinesin competes with tau for microtubule binding sites (37-43). This results in presence of a reservoir of membrane vesicles in the distal neurite shaft near the growth cone. Using these specifications, we determined the size of reservoir under various drug treatment conditions. The size of the membrane reservoir near the growth cone would be indicative of the capability of axonal growth in response to treatment with different drugs and the four -drug combination. As the reservoir is depleted more membrane is added to the growth cone allowing the neurite to grow longer. Some but not all of the kinetic parameters are known. Consequently, we used an analytical approach to calculate the remaining parameter when an overall experimentally measured constrained such as velocity of axonal outgrowth is measured. The validity of this approach was established by experimental validation of computational predictions for the formation of dystrophic bulbs and velocity of neurite outgrowth Arjun's Paper,). The overall model used for the calculation of kinetic parameters from the analytical solution is shown below

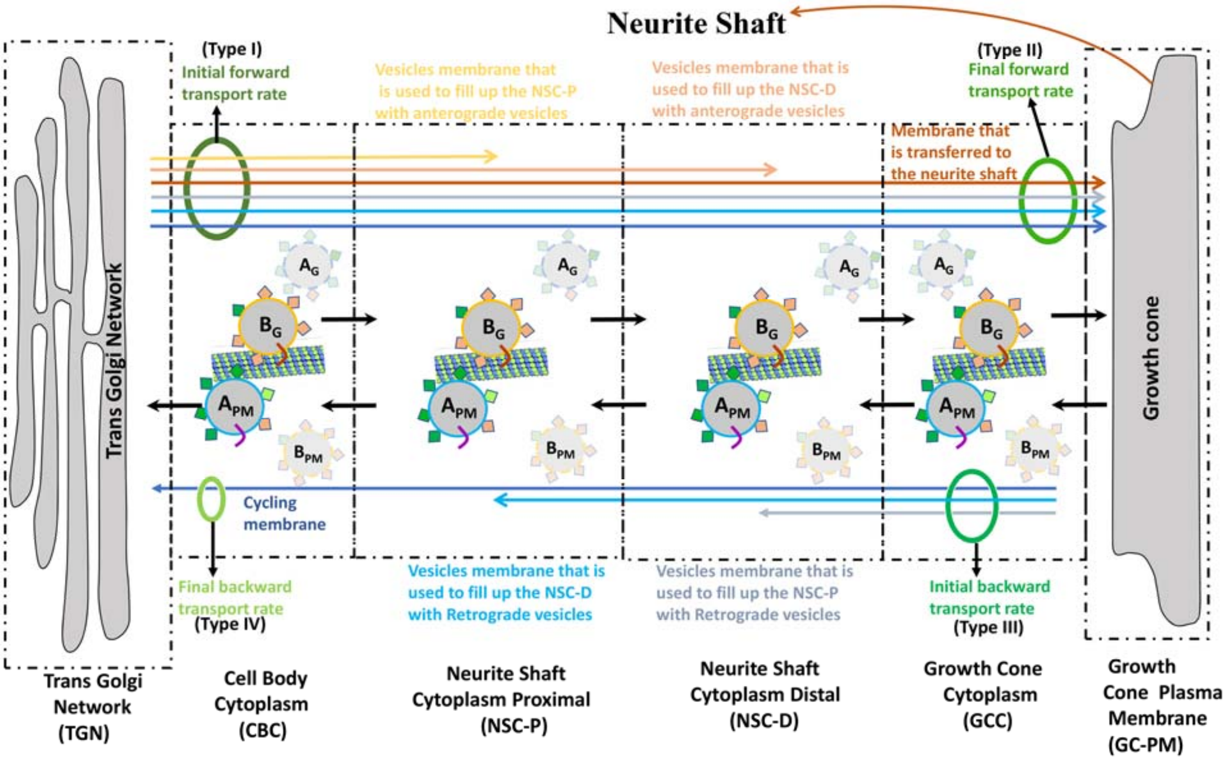


**Supplementary Figure** 9: Schematic representation of the model used for generation of an analytical solution for the prediction of kinetic parameters at steady state. The development of the analytical solution depends on the categorization of membrane fluxes based on their destination compartment. Such classification distinguishes four different membrane types (I-IV) which are colored in four different colors (Green color ovals) from TGN to growth cone. Initial and final anterograde (initial forward transport rate & final forward transport rate) membrane transport rates refer to budding and fusion rates at the TGN and GC respectively and retrograde (initial backward transport rate & final backward transport rate) membrane transport rates represent endocytosis rate at growth cone and fusion rate at TGN. Budded vesicles membrane at TGN are divided into six different types of membranes based on their destination (i) Yellow color arrow represent membrane loss of anterograde vesicles in NSC-P compartment, (ii) Light brown color arrow represent membrane loss of anterograde vesicles in NSC-D compartment (iii) The brown arrow from the growth cone plasma membrane (GC-PM) to the shaft membrane represents the rate of membrane transfer that allows the growth cone to maintain its constant size and the neurite shaft to grow in length. (iv) Light gray arrow from the growth cone membrane to NSC-D, represent membrane loss of retrograde moving vesicles (v) Light blue color arrow from GC-PM to NSC-P represent the membrane loss of retrograde moving vesicles in NSC-P compartment (vi) The dark blue color arrow from GC-PM to TGN represent the recycling membrane from GC-PM to TGN. Here anterograde moving vesicles are ($A_G$ & $B_G$) where $A_G$ does not move forward because coat A budded vesicles SNAREs have higher affinity with SNAREs at TGN and hence back fuse to the TGN. Similarly, endocytosed vesicles are ($A_{PM}, B_{PM}$) where $B_{PM}$ vesicles back fuse with GC-PM.

**Model of Drug Action on Axonal Regeneration**

Since HU-210 and IL-6 stimulate transcription, we have assumed that this stimulation leads to biosynthesis of molecules needed for axon regeneration at sufficient levels at all times. Based on this assumption the levels of all components are buffered in the model when the cell body is stimulated by HU-210 + IL-6.

The experimentally observed velocities of axonal growth for different drugs and drug combinations were used as a constraint for our model to calculate required membrane fusion at growth cone plasma membrane. The model constraints specified the rate of back-transported membrane from the growth cone plasma membrane to the TGN and the fraction of moving and stationary vesicles in the CBC, NSC-P, NSC-D and GCC compartments (Supp Figure 9) We assumed that endocytosed vesicles should transport 0.5 $\mu m^2$ membrane/min back to the TGN based on experiments in the literature (18) and calculated other parameters of the model analytically for a given neurite outgrowth velocity. Velocities of growth were obtained from the morphology experiments. We determined these rates based upon in vivo axonal regeneration, where we graphed the Intensity of Fluorescent signal of CTB over distance (µm) from the crush site and measured average growth with the indicated combinations or single drug over a 3 weeks period. They then divided the total average length of growth by 21days and determined the average velocities. In Supplementary Figures 10A&B we show at steady state, the distribution of all membrane proteins (SNAREs, motor proteins and recruitment factors – Coat proteins A (retrograde) & B (anterograde)), membrane (vesicles) and fluxes in the terminal and intermediate compartments used in the model. (Supplementary Figure 10).

Application of taxol and APC in gel foam at the site of injury has two effects. Taxol increases stabilization of the dynamic microtubule thus enabling microtubule elongation. APC clears extracellular inhibitory environment at injury site which can increase intracellular vesicles mobility from vesicles reservoir NSC-D to growth cone plasma membrane (GC-PM). Experimentally HU-210 and IL-6 applied at cell body allow neurite outgrowth with a velocity v = 0.5 µm/h. When Taxol along with HU-210 + IL-6 is applied, neurites grow at 1 µm/h. Although Taxol stabilizes dynamic microtubules and we should achieve faster growth rates, overall growth of microtubule bundle inhibited by external debris and hence it cannot grow faster despite having fast growing capacity. It is known that when growing microtubules hit, the membrane they collapse (44). In case of APC application at injury site, neurite outgrowth shows a velocity v = 2 µm/h because APC clears inhibitory environment, hence vesicles can move from vesicle reservoir NSC-D to GCC and fuse with growth cone plasma membrane (0.6052 $\frac{\mu m^2}{min}, calculated\ analytically$). However, since there is insufficient microtubule elongation the neurite does not grow very fast.

Analytical solutions (equations 1-158 shown below ) for the model identified how the four drug combination could allow axons to grow with a velocity v = 16 µm/h. Depletion of the membrane vesicle pool in the NSC-D compartment allows for the expansion of the growth cone plasma membrane which in turn enables the microtubule’s to elongate without inhibition. . A threedimensional plot relating axon length to apparent GTPase activity of the microtubules and concentrations of vesicles in the NSC-D compartment is shown in. XA in the main paper The calculated values for the vesicles levels in the NSC-D compartment under the various drug conditions are shown in and corresponding rates of microtubule elongation and axon length are shown in XC and XD in the main paper. The profile of the plots indicate that the seemingly synergistic effects of the four drug combination can be explained parsimoniously by the action of APC, which by removing the extracellular inhibitory environment allows for vesicle movement from the NCS-D compartment to the GC-PM compartment. This increase in the plasma membrane at the e growth cone coordinated with the enhanced growth rate of the microtubule due to Taxol dependent stabilization of the dynamic microtubules allows for the observed overall growth rate of the axon. Overall our model indicates that relief of inhibition of growth cone plasma membrane expansion is sufficient to account for the observed experimental growth rates.

**Analytical Solution**

The development of the analytical solution is extension and rearrangement of previously developed model (18). Here, we assumed initial forward transport rate at TGN is constant because HU-210 and IL-6 applied at cell body by intravitreal injection increases the capacity of membrane production at TGN so transport is not limited by availability of membrane vesicles. To achieve a specific axonal outgrowth velocity, we calculated fraction of bound kinesin mediate membrane vesicle transport from the reservoir (NSC-D) to GCC. Membrane that is incorporated into vesicles that bud from the TGN can be separated into 4 different membrane vesicle types (I, II, III, IV). Membrane vesicles of the type-I will be added to the growing neurite shaft cytoplasm distal (NSC-D) as part of the neurite shaft cytoplasm reservoir for anterograde moving vesicles where NSC-D work as a reservoir next to growth cone which can be mobilized on sudden demand for membrane at growth cone. Type-II membrane vesicles will be added to the GC and from there to the growing neurite shaft. Type-III and type-IV membrane vesicles will also be added to the GC, but the fused membrane will then be incorporated into endocytic retrograde vesicles. Similar to first membrane vesicle set, the third vesicle set will be added to the growing neurite shaft cytoplasm, though this set is much smaller due to an assumed nine-fold lower amount of stationary retrograde vesicles in the neurite shaft cytoplasm. The fourth membrane vesicle set will be back transported to the TGN, i.e. this set constitutes the cycling membrane between the TGN and the GC. As a simplification, we assume in our solution that membranes transported in an anterograde manner is only transported as vesicles that bud from the TGN with coat protein B and retrogradely transported membranes are vesicles that bud from the GC with coat protein A.

The microtubule dynamics model depends on dynamic microtubule profile, nucleation rate, degradation rate, stabilization rate and hydrolysis rates. We used Margolin model (31) for simulation of dynamic microtubules for fixed effective tubulin concentration and different hydrolysis rates (0.60 $sec^{-1}$ – 0.80 $sec^{-1}$). To calculate the microtubule bundle length in axonal shaft, we calculated the length of dynamic and stable microtubules and divided by the number of microtubules in cross section. Dynamic microtubules have a stochastic profile, which changes as they undergo catastrophic disassembly by hydrolysis of GTP bound tubulins at tip. Application of taxol and APC in gel foam at the site of injury create four different criteria of GTP hydrolysis rate of dynamic microtubules. We call (i) 'apparent base hydrolysis rate' in absence of (Taxol & APC) where catastrophic disassembly of dynamic microtubules happens due to GTP hydrolysis as well as external inhibitory signal at crush site; (ii) 'apparent hydrolysis rate under taxol' when taxol present at injury site which reduce GTP hydrolysis rate but not external inhibitory signal, (iii) 'base hydrolysis rate' when APC alone is present at injury site, (iv) 'hydrolysis rate under taxol' when (Taxol & APC) both drugs are present at injury site, in this case taxol reduces hydrolysis rate. We found ascending order relationship in (i) > (ii) > (iii) > (iv) for these four conditions.

For an experimentally observed axonal outgrowth velocity, the amount of cycling membrane vesicles and percentage of stationary anterograde vesicles in (NSC-P & NSC-D) and retrograde vesicles in (NSC-P & NSC-D) allows for the calculation of the fluxes of the four different membrane vesicle types. Since HU210 and IL-6, applied at cell body increases biosynthesis of molecules and membrane vesicles at Trans Golgi Network such that membrane vesicle production is not limiting. We calculated the optimum value of "Initial forward transport rate" which satisfies the growth velocities of all drug combination by satisfying know biological constraints such as ( recycling membrane rate, retrograde vesicles motor binding rate with microtubule bundle and mobilization of anterograde vesicles from NSC-D compartment to growth cone plasma membrane with full capacity. We need to produce membrane budding at TGN compartment is 1.4001 $\mu m^2/min$.

$$initial\ forward\ transport\ rate = 1.4001\ \frac{\mu m^2}{min} \quad (1)$$

**Type II membrane vesicle transport rate (Final forward transport rate):**

$$Growth\ related\ k\ membrane\ production = \frac{Axonal\ outgrowth\ velocity}{2\pi\ \times radius\ of\ neurite}\ . \quad (2)$$

**Type IV membrane vesicle transport rate (Final Backward transport rate):**

$$final\ back\ transport\ rate = cycling\ rate\ , \quad (3)$$

$$a2\ travel\ time\ new\ NSCP = \frac{Axonal\ outgrowth\ velocity}{v_d} \quad (4)$$

$$a2\ travel\ time\ new\ NSCD = \frac{Axonal\ outgrowth\ velocity}{v_d} \quad (5)$$

$$rate\ lost\ na2\ NSCP = \frac{final\ backward\ transport\ rate \times a2\ travel\ time\ new\ NSCP}{anticipated\ fraction\ of\ bound\ a2\ vesicles\ in\ NSCP} \quad (6)$$

$$rate\ lost\ na2\ NSCD = \frac{final\ backward\ transport\ rate \times a2\ travel\ time\ new\ NSCD}{anticipated\ fraction\ of\ bound\ a2\ vesicles\ in\ NSCD} \quad (7)$$

**Type III membrane vesicle transport rate (Initial back transport rate):**

$$
\begin{aligned}
&initial\ back\ transport\ rate \\
&\qquad = final\ backward\ transport\ rate \\
&\qquad + \ rate\ of\ accumulation\ of\ stationary\ na_2\ NSCP \\
&\qquad + \ rate\ of\ accumulation\ of\ stationary\ na_2\ NSCD
\end{aligned} \tag{8}
$$

$$
\begin{aligned}
&Final\ forward\ transport\ rate = initial\ back\ transport\ rate\ + \\
&\qquad\qquad Growth\ related\ k\ membrane\ production
\end{aligned} \tag{9}
$$

$$
b_1\ travel\ time\ new\ NSCP = \frac{Axonal\ outgrowth\ velocity}{v_k} \tag{10}
$$

$$
b_1\ travel\ time\ new\ NSCD = \frac{Axonal\ outgrowth\ velocity}{v_k} \tag{11}
$$

$$
\begin{aligned}
&rate\ of\ accomulation\ of\ stationary\ vesicles\ (surface\ area)\ nb_1 NSCP \\
&\qquad = Initial\ forward\ transport\ rate \times b_1\ travel\ time\ new\ NSCP \\
&\qquad \times fraction\ of\ bound\ b_1\ vesicles\ in\ NSCP
\end{aligned} \tag{12}
$$

$$
\begin{aligned}
&rate\ of\ accomulation\ of\ stationary\ vesicles\ (surface\ area)\ nb_1 NSC\mathrm{D} \\
&= Initial\ forward\ transport\ rate - Final\ forward\ transport\ rate \\
&- rate\ of\ accomulation\ of\ stationary\ vesicles\ (surface\ area)\ nb_1 NSCP
\end{aligned} \tag{13}
$$

$$
\begin{aligned}
&fraction\ of\ bound\ b_1 vesicles\ in\ NSC\mathrm{D} \\
&= \frac{Final\ forward\ transport\ rate \times b_1 travel\ time\ new\ NSCD}{rate\ of\ accomulation\ of\ stationary\ vesicles\ (surface\ area)\ nb_1 NSC\mathrm{D}}
\end{aligned} \tag{14}
$$

**Type I membrane vesicle transport rate (Initial forward transport rate):**

$$
\begin{aligned}
&rate\ of\ accomulation\ of\ stationary\ vesicles (surface\ area) in\ NSC \\
&\qquad = rate\ of\ accumulation\ of\ stationary\ vesicles\ (surface\ area)\ in\ NSCP \\
&\qquad + rate\ of\ accumulation\ of\ stationary\ vesicles\ (surface\ area) in\ NSCD \\
&\qquad = (rate\ of\ accumulation\ of\ stationary\ na_2\ in\ NSCP \\
&\qquad + \ rate\ of\ accumulation\ of\ stationary\ nb_1\ in\ NSCP) \\
&\qquad + (rate\ of\ accumulation\ of\ stationary\ na_2\ in\ NSCD \\
&\qquad + \ rate\ of\ accumulation\ of\ stationary\ nb_1\ in\ NSCD)
\end{aligned} \tag{15}
$$

$$w_G^B = \frac{initial\ forward\ transport\ rate}{cc1_G \times s_G}, \quad (16)$$

$$w_{PM}^A = \frac{initial\ backward\ transport\ rate}{cc2_{PM} \times s_{PM}}, \quad (17)$$

$$w_{PM}^B = w_G^B \times factor\ for\ site\ specific\ budding, \quad (18)$$

$$w_G^A = w_{PM}^A \times factor\ for\ site\ specific\ budding. \quad (19)$$

The fluxes also allowed us to calculate the initial membrane surface areas in the cytoplasmic compartments.

$$b_1\ travel\ time\ NSCP\ at\ start = \frac{NSCP\ length\ at\ start}{v_k}, \quad (20)$$

$$b_1\ travel\ time\ NSCD\ at\ start = \frac{NSCD\ length\ at\ start}{v_k}, \quad (21)$$

$$n_{MTCP}^{B_G} = \frac{final\ forward\ transport\ rate\ \times\ b_1\ travel\ time\ NSCP\ at\ start}{anticipated\ fraction\ of\ bound\ b_1\ vesicles\ in\ NSCP}, \quad (22)$$

$$n_{MTCD}^{B_G} = \frac{final\ forward\ transport\ rate\ \times b_1\ travel\ time\ NSCD\ at\ start}{anticipated\ fraction\ of\ bound\ b1\ vesicles\ in\ NSCD}, \quad (23)$$

$$n_{GCC}^{B_G} = \frac{final\ forward\ transport\ rate}{anticipated\ fraction\ of\ fusing\ b_1\ \ vesicles\ in\ GCC}, \quad (24)$$

$$a_2\ travel\ time\ NSCP\ at\ start = \frac{NSCP\ length\ at\ start}{v_d}, \quad (25)$$

$$a_2\ travel\ time\ NSCD\ at\ start = \frac{NSCD\ length\ at\ start}{v_d}, \quad (26)$$

$$n_{NSCP}^{A_{PM}} = \frac{final\ backword\ transport\ rate\ \times\ a_2\ travel\ time\ NSCP\ at\ start}{anticipated\ fraction\ of\ bound\ a_2\ vesicles\ in\ NSCP}, \quad (27)$$

$$n_{NSCD}^{A_{PM}} = \frac{final\ backword\ transport\ rate\ \times\ a_2\ travel\ time\ NSCD\ at\ start}{anticipated\ fraction\ of\ bound\ a_2\ vesicles\ in\ NSCD}, \quad (28)$$

$$n_{CBC}^{A_{PM}} = \frac{final\ backword\ transport\ rate}{anticipated\ fraction\ of\ fusion\ a_2\ vesicles\ in\ CBC}. \quad (29)$$

Similarly, to the fraction of anterograde vesicles that are bound to the microtubule and therefore moving in the neurite shaft cytoplasm, we define the fractions of moving anterograde vesicles in the GC cytoplasm (i.e. vesicles that fuse with the GC membrane) and moving retrograde vesicles in the CB cytoplasm (i.e. vesicles that fuse with the TGN). The anticipated membrane fusion rate at the growth cone (or final forward transport rate) is equal to the sum of the fluxes of the second, third and fourth membrane types, the anticipated membrane fusion rate at the TGN (or final backward transport rate) is equal to the flux of the fourth membrane type. This allows the calculation of the number of v-SNAREs that are associated with each set of vesicles in both cytoplasmic compartments. We want to estimate the amount of v-SNAREs V in the GC cytoplasm or U in the CB cytoplasm that enables dynamic balance For this the fractions of fusing vesicles should equal the anticipated fractions delivered to the compartment and that the total membrane fluxes equal the final forward (type II) and final backward (Type IV) membrane fluxes for a predefined amount of t-SNAREs Y at the TGN and X at the GC for specified tethering rates tethering rates.

$$kappaXU_{pm} = kappaXU_g \times factor\ for\ site\ specific\ SNARE\ complex\ formation\ , \quad (30)$$

$$kappaXU_g = kappaXU_{pm} \times factor\ for\ site\ specific\ SNARE\ complex\ formation\ , \quad (31)$$

$$YV\ SNARE\ complexes\ GCC\ PM = \frac{final\ forward\ transport\ rate}{anterograde\ vesicle\ surface\ area} \times SNARE\ complex\ per\ vesicle\ fusion\ , \quad (32)$$

$$VV_{GCC}^{B_G} = \frac{YV\ SNARE\ complexes\ GCC\ PM}{kappaYV_{pm}} \times YY_{pm}\ , \quad (33)$$

$$V_{GCC}^{B_G} = \frac{VV_{GCC}^{B_G}}{n_{GCC}^{B_G}}\ . \quad (34)$$

Vesicles that fuse with the TGN or the GC transmit their SNAREs to the target organelle. Consequently, the vesicles that enter the CB cytoplasm or GC cytoplasm from the NSC should contain the same number of SNAREs to keep the number of SNARE molecules in the cytoplasmic compartments constant. Additionally, v-SNAREs need to be back transported to the GC or the TGN to be available for the next set of budding vesicles. The fluxes for each v-SNARE can be associated to 4 different types in a similar manner as the membrane fluxes. Initial and final forward and backward fluxes can be calculated accordingly.

Similarly, to the v-SNAREs V, we calculate the corresponding parameter for v-SNARE U.

$$XU\ SNARE\ complexes\ CBC\ G = \frac{final\ backtransport\ rate}{anterograde\ vesicle\ surface\ area} \times SNARE\ complex\ per\ vesicle\ fusion\ , \quad (35)$$

$$UU_{CBC}^{A_{PM}} = \frac{XU\ SNARE\ complexes\ CBC\ G}{kappaXU_G} \times XX_G\ , \quad (36)$$

$$U_{CBC}^{A_{PM}} = \frac{UU_{CBC}^{A_{PM}}}{n_{CBC}^{A_{PM}}} , \quad (37)$$

$$U\_backward = U_{CBC}^{A_{PM}} , \quad (38)$$

$$UU\ final\ backward = U\_backward \times final\ backtransport\ rate, \quad (39)$$

$$UU\ initial\ backward = U\_backward \times initial\ backtransport\ rate, \quad (40)$$

$$UU\ final\ forward = UU\ Initial\ backward , \quad (41)$$

$$U_{backward} = \frac{UU\ final\ forward}{final\ forward\ transport\ rate} , \quad (42)$$

$$UU\ initial\ forward = initial\ forward\ transport\ rate \times U\_forward. \quad (43)$$

SNAREs are recruited into the budding vesicles in a competitive manner, i.e. all four SNAREs compete with each other for the SNARE binding spots of the budding vesicles. The amounts of t-SNAREs Y and X at the GC and the TGN are predefined. Considering this we calculated the amount of v-SNAREs V and U at the TGN and GC that allow the calculated initial forward and backward v-SNARE fluxes.

$$Y_{PM} = \frac{YY_{PM}}{s_{PM}}, \quad (44)$$

$$X_{PM} = kxa \times \frac{Y_{PM}}{kya}, \quad (45)$$

$$cont_{PM} = 1 + \frac{Y_{PM}}{kya} + \frac{X_{PM}}{kxa}, \quad (46)$$

$$bs = initial\ backtransport\ rate \times snare\ binding\ site\ per\ vesicle\ area , \quad (47)$$

$$U_{PM} = \frac{UU\ initial\ backward \times cont_{PM} \times kua}{\left(bs - (UU\ initial\ backward + VV\ initial\ backward\ )\right)} , \quad (48)$$

$$V_{PM} = \frac{VV\ initial\ backward \times cont_{PM} \times kva}{\left(bs - (UU\ initial\ backward + VV\ initial\ backward\ )\right)} , \quad (49)$$

$$UU_{PM} = U_{PM} \times s_{PM} , \quad (50)$$

$$XX_{PM} = X_{PM} \times s_{PM}. \quad (51)$$

Though t-SNAREs have a higher dissociation constant for association with the vesicles than v-SNAREs they still are transported between the TGN and GC. We calculate their transport rates in a similar manner as for the vesicle SNAREs.

$$snare\ saturation\ denom\ 2a = 1 + \frac{X_{PM}}{kxa} + \frac{U_{PM}}{kua} + \frac{Y_{PM}}{kya} + \frac{V_{PM}}{kva}\ , \quad (52)$$

$$sya2 = \frac{snare\ binding\ site\ per\ vesicle\ area \times \frac{Y_{PM}}{kya}}{snare\ saturation\ denom\ 2a}\ , \quad (53)$$

$$YY\ initial\ backward = sya2 \times initial\ backtransport\ rate\ , \quad (54)$$

$$YY\ final\ backward = sya2 \times final\ backtransport\ rate\ , \quad (55)$$

$$Y\ initial\ backward = \frac{YY\ initial\ backward}{initial\ backtransport\ rate}\ , \quad (56)$$

$$YY\ final\ forward = YY\ initial\ backward, , \quad (57)$$

$$Y\ forward = \frac{YY\ final\ forward}{final\ forward\ transport\ rate}\ , \quad (58)$$

$$Y\ initial\ forward = Y\ forward \times initial\ forward\ transport\ rate\ .\ , \quad (59)$$

The 'consumption' of membrane proteins that are associated with the vesicles that are added to the NSC reservoir demands the continuous production of membrane proteins at the TGN. The protein production rates can be calculated by considering the concentration of each protein at anterograde and retrograde moving vesicles and the amount of membrane that will be added to the NSC reservoir as shown for the SNAREs V and Y.

$$\begin{aligned} k\ VV\ production \\ &= (rate\ of\ accumulation\ of\ stationary\ nb_1\ NSCP \times V\ forward \\ &+ rate\ of\ accumulation\ of\ stationary\ na_2\ NSCP \times V\ backward) \\ &+ (rate\ of\ accumulation\ of\ stationary\ nb_1\ NSCD \times V\ forward \\ &+ rate\ of\ accumulation\ of\ stationary\ na_2\ NSCD \times V\ backward) \end{aligned} \quad (60)$$

$$\begin{aligned} k\ YY\ production \\ &= (rate\ of\ accumulation\ of\ stationary\ nb_1\ NSCP \times Y\ forward \\ &+ rate\ of\ accumulation\ of\ stationary\ na_2\ NSCP \times Y\ backward) \\ &+ (rate\ of\ accumulation\ of\ stationary\ nb_1\ NSCD \times Y\ forward \\ &+ rate\ of\ accumulation\ of\ stationary\ na_2\ NSCD \times Y\ backward) \end{aligned} \quad (61)$$

The concentration of the t-SNARE Y at the TGN is calculated via the following equations.

$$X_G = \frac{XX_G}{s_G}, \quad (62)$$

$$YY_G = \max\left(0, Y\ forward \times \frac{X_G}{kxb} \times kyb \times s_G - k\ YY\ production\right). \quad (63)$$

TGN concentrations for the v-SNARE V and U are calculated in a similar manner as their concentrations at the GC.

t-SNARE X movements are calculated similarly as t-SNARE Y movements.

$$snare\ saturation\ denom\ 1b = 1 + \frac{X_G}{kxb} + \frac{U_G}{kub} + \frac{Y_G}{kyb} + \frac{V_G}{kvb}, \quad (64)$$

$$sxb1 = \frac{snare\ binding\ spot\ per\ vesicle\ area \times \frac{X_G}{kxb}}{snare\ saturation\ denom\ 1b}, \quad (65)$$

$$X\ forward = sxb1, \quad (66)$$

$$XX\ initial\ forward = X\ forward \times initial\ forward\ transport\ rate, \quad (67)$$

$$XX\ final\ forward = X\ forward \times final\ forward\ transport\ rate, \quad (68)$$

$$XX\ initial\ backward = XX\ final\ forward, \quad (69)$$

$$X\ backward = \frac{XX\ initial\ forward}{initial\ backward\ transport\ rate}, \quad (70)$$

$$XX\ final\ backward = X\ backward \times final\ backward\ transport\ rate. \quad (71)$$

Production rates for SNAREs U and X are calculated.

SNARE X, U and V protein amounts at the TGN are calculated:

$$\begin{aligned} k\ UU\ production &= (rate\ of\ accumulation\ of\ stationary\ nb_1\ NSCP \times U\ forward \\ &+ rate\ of\ accumulation\ of\ stationary\ na_2\ NSCP \times U\ backward) \\ &+ (rate\ of\ accumulation\ of\ stationary\ nb_1\ NSCD \times U\ forward \\ &+ rate\ of\ accumulation\ of\ stationary\ na_2\ NSCD \times U\ backward) \end{aligned} \quad (72)$$

$$\begin{aligned} k\ XX\ production &= (rate\ of\ accumulation\ of\ stationary\ nb_1\ NSCP \times X\ forward \\ &+ rate\ of\ accumulation\ of\ stationary\ na_2\ NSCP \times X\ backward) \\ &+ (rate\ of\ accumulation\ of\ stationary\ nb_1\ NSCD \times X\ forward \\ &+ rate\ of\ accumulation\ of\ stationary\ na_2\ NSCD \times X\ backward) \end{aligned} \quad (73)$$

Protein for all vesicle membrane proteins in the different compartments are calculated:

$$UU_G = \max(0, (UU_G - k\ UU\ production)), \quad (74)$$

$$XX_G = \max(0, (XX_G - k\ XX\ production)), \quad (75)$$

$$VV_G = \max(0, (VV_G - k\ VV\ production)). \quad (76)$$

We calculated the amount of motor protein receptors that are associated with anterograde and retrograde vesicles in the neurite shaft cytoplasm that is necessary to allow the anticipated fraction of microtubule bound (i.e. moving) vesicles. This enabled the calculation of the motor protein receptors in the other cytoplasmic compartments and the amount of motor protein receptors at the TGN and GC as well as their production rates in a similar manner as we did it for the v-SNAREs.

$$\left.\begin{array}{r} VV_{CBC}^{B_G} = n_{CBC}^{B_G} \times V\ forward \\ VV_{NSC}^{B_G} = n_{NSC}^{B_G} \times V\ forward \\ VV_{GCC}^{B_G} = n_{GCC}^{B_G} \times V\ forward \\ VV_{GCC}^{A_{PM}} = n_{GCC}^{A_{PM}} \times V\ backward \\ VV_{NSCP}^{A_{PM}} = n_{NSCP}^{A_{PM}} \times V\ backward \\ VV_{NSCD}^{A_{PM}} = n_{NSCD}^{A_{PM}} \times V\ backward \\ VV_{CBC}^{A_{PM}} = n_{CBC}^{A_{PM}} \times V\ backward \end{array}\right\}, \quad (77)$$

$$\left.\begin{array}{r} UU_{CBC}^{B_G} = n_{CBC}^{B_G} \times U\ forward \\ UU_{NSC}^{B_G} = n_{NSC}^{B_G} \times U\ forward \\ UU_{GCC}^{B_G} = n_{GCC}^{B_G} \times U\ forward \\ UU_{GCC}^{A_{PM}} = n_{GCC}^{A_{PM}} \times U\ backward \\ UU_{NSCP}^{A_{PM}} = n_{NSCP}^{A_{PM}} \times U\ backward \\ UU_{NSCD}^{A_{PM}} = n_{NSCD}^{A_{PM}} \times U\ backward \\ UU_{CBC}^{A_{PM}} = n_{CBC}^{A_{PM}} \times U\ backward \end{array}\right\}, \quad (78)$$

$$\left.\begin{array}{r} XX_{CBC}^{B_G} = n_{CBC}^{B_G} \times X\ forward \\ XX_{NSC}^{B_G} = n_{NSC}^{B_G} \times X\ forward \\ XX_{GCC}^{B_G} = n_{GCC}^{B_G} \times X\ forward \\ XX_{GCC}^{A_{PM}} = n_{GCC}^{A_{PM}} \times X\ backward \\ XX_{NSCP}^{A_{PM}} = n_{NSCP}^{A_{PM}} \times X\ backward \\ XX_{NSCD}^{A_{PM}} = n_{NSCD}^{A_{PM}} \times X\ backward \\ XX_{CBC}^{A_{PM}} = n_{CBC}^{A_{PM}} \times X\ backward \end{array}\right\}, \quad (79)$$

$$\left.\begin{aligned} YY_{CBC}^{B_G} &= n_{CBC}^{B_G} \times Y\,forward \\ YY_{NSC}^{B_G} &= n_{NSC}^{B_G} \times Y\,forward \\ YY_{GCC}^{B_G} &= n_{GCC}^{B_G} \times Y\,forward \\ YY_{GCC}^{A_{PM}} &= n_{GCC}^{A_{PM}} \times Y\,backward \\ YY_{NSCP}^{A_{PM}} &= n_{NSCP}^{A_{PM}} \times Y\,backward \\ YY_{NSCD}^{A_{PM}} &= n_{NSCD}^{A_{PM}} \times Y\,backward \\ YY_{CBC}^{A_{PM}} &= n_{CBC}^{A_{PM}} \times Y\,backward \end{aligned}\right\}. \tag{80}$$

Finally, we calculated the fluxes for the other stationary proteins, i.e. recruitment factors 1 and 2 as well as the initial protein amounts in the different cytoplasmic compartments.

$$KK_{NSCP}^{B_G} = \frac{log(1 - anticipated\ fraction\ of\ bound\ vesicles\ in\ NSCP)}{log(1 - fraction\ bound\ kk\ a1b1\ NSCP)}, \tag{81}$$

$$KK_{NSCD}^{B_G} = \frac{log(1 - anticipated\ fraction\ of\ bound\ vesicles\ in\ NSCD)}{log(1 - fraction\ bound\ kk\ a1b1\ NSCD)}, \tag{82}$$

$$K_{NSCP}^{B_G} = \frac{KK_{NSCP}^{B_G}}{anterograde\ vesicles\ surface\ area}, \tag{83}$$

$$K_{NSCD}^{B_G} = \frac{KK_{NSCD}^{B_G}}{anterograde\ vesicles\ surface\ area}, \tag{84}$$

$$K\ forward = K_{NSCD}^{B_G}, \tag{85}$$

$$KK\ initial\ forward = K\ forward \times initial\ forward\ transport\ rate, \tag{86}$$

$$KK\ final\ forward = K\ forward \times final\ forward\ transport\ rate, \tag{87}$$

$$KK\ initial\ backward = KK\ final\ forward, \tag{88}$$

$$K\ backward = \frac{KK\ initial\ backward}{initial\ backward\ traport\ rate}, \tag{89}$$

$$MBSPVA = motor\ protein\ binding\ sites\ per\ vesicles\ area,$$

$$K_G = \frac{kkb \times KK\ initial\ forward}{initial\ forward\ traport\ rate \times MBSPVA - KK\ initial\ forward}, \tag{90}$$

$$K_{PM} = \frac{kka \times KK\ initial\ backward}{initial\ backward\ traport\ rate \times MBSPVA - KK\ initial\ backward}, \tag{91}$$

$$KK_G = K_G \times s_G, \tag{92}$$

$$KK_{PM} = K_{PM} \times s_{PM}, \tag{93}$$

$$KK_{CBC}^{B_G} = k\ forward \times n_{CBC}^{B_G}, \quad (94)$$

$$KK_{NSC}^{B_G} = k\ forward \times n_{NSC}^{B_G}, \quad (95)$$

$$KK_{GCC}^{B_G} = k\ forward \times n_{GCC}^{B_G}, \quad (96)$$

$$KK_{CBC}^{B_{PM}} = k\ backward \times n_{CBC}^{A_{PM}}, \quad (97)$$

$$KK_{NSC}^{B_{PM}} = k\ backward \times n_{NSC}^{A_{PM}}, \quad (98)$$

$$KK_{GCC}^{B_{PM}} = k\ backward \times n_{GCC}^{A_{PM}}, \quad (99)$$

$$\begin{aligned} k\ KK\ production &= (rate\ of\ accumulation\ of\ stationary\ nb_1\ NSCP \times K\ forward \\ &+ rate\ of\ accumulation\ of\ stationary\ na_2\ NSCP \times K\ backward) \\ &+ (rate\ of\ accumulation\ of\ stationary\ nb_1\ NSCD \times K\ forward \\ &+ rate\ of\ accumulation\ of\ stationary\ na_2\ NSCD \times K\ backward) \end{aligned} \quad (100)$$

$$DD_{NSCP}^{B_G} = \frac{log(1 - anticipated\ fraction\ of\ bound\ vesicles\ in\ NSCP)}{log(1 - fraction\ bound\ kk\ a1b1\ NSCP)}, \quad (101)$$

$$DD_{NSCD}^{B_G} = \frac{log(1 - anticipated\ fraction\ of\ bound\ vesicles\ in\ NSCD)}{log(1 - fraction\ bound\ kk\ a1b1\ NSCD)}, \quad (102)$$

$$D_{NSCP}^{B_G} = \frac{DD_{NSCP}^{B_G}}{anterograde\ vesicles\ surface\ area}, \quad (103)$$

$$D_{NSCD}^{B_G} = \frac{DD_{NSCD}^{B_G}}{anterograde\ vesicles\ surface\ area}, \quad (104)$$

$$D\ backward = DD_{NSC}^{A_{PM}}, \quad (105)$$

$$DD\ initial\ backward = D\ backward \times initial\ backward\ transport\ rate, \quad (106)$$

$$DD\ final\ forward = DD\ initial\ backward, \quad (107)$$

$$DD\ initial\ forward = D\ forward \times initial\ forward\ transport\ rate, \quad (108)$$

$$D_{G=} = \frac{kdb \times DD\ initial\ forward}{initial\ forward\ traport\ rate \times MBSPVA - DD\ initial\ forward}, \quad (109)$$

$$K_{PM} = \frac{kda \times \ DD\ initial\ backward}{initial\ backward\ traport\ rate \times MBSPVA - DD\ initial\ backward}\,, \tag{110}$$

$$DD_G = D_G\ \times s_G\,, \tag{111}$$

$$DD_{PM} = D_{PM}\ \times s_{PM}\,, \tag{112}$$

$$DD_{CBC}^{B_G}\ = D\ forward \times n_{CBC}^{B_G}\,, \tag{113}$$

$$DD_{NSC}^{B_G}\ = D\ forward \times n_{NSC}^{B_G}\,, \tag{114}$$

$$DD_{GCC}^{B_G}\ = D\ forward \times n_{GCC}^{B_G}, \tag{115}$$

$$DD_{CBC}^{A_{PM}} = D\ backward \times n_{CBC}^{A_{PM}}, \tag{116}$$

$$DD_{NSC}^{A_{PM}} = D\ backward \times n_{NSC}^{A_{PM}}, \tag{117}$$

$$DD_{GCC}^{A_{PM}} = D\ backward \times n_{GCC}^{A_{PM}}, \tag{118}$$

$$\begin{aligned} k\ DD\ production \\ &= (rate\ of\ accumulation\ of\ stationary\ nb_1\ NSCP \times D\ forward \\ &+\ rate\ of\ accumulation\ of\ stationary\ na_2\ NSCP \times D\ backward) \\ &+ (rate\ of\ accumulation\ of\ stationary\ nb_1\ NSCD \times D\ forward \\ &+ rate\ of\ accumulation\ of\ stationary\ na_2\ NSCD\ \times D\ backward) \end{aligned} \tag{119}$$

$$C1_G = \frac{CC1_G}{s_G}\,, \tag{120}$$

$CBSPVA =\ cargo\ binding\ sites\ per\ unit\ vesicle\ area.$

$$sc1b1 = \frac{CBSPVA\ \times \frac{C1_G}{kc1b}}{\left(1 + \frac{C1_G}{kc1b}\right)}\,, \tag{121}$$

$$C1\ forward =\ sc1b1, \tag{122}$$

$$CC1\ final\ forward =\ C1\ forward\ \times final\ forward\ transport\ rate, \tag{123}$$

$$CC1\ initial\ backward =\ CC1\ final\ forward, \tag{124}$$

$$C1\ backward =\ \frac{CC1\ initial\ backward}{initial\ backtransport\ rate}\,, \tag{125}$$

$$C1_{PM} =\ \frac{kc1a \times\ CC1\ initial\ backward}{initial\ backward\ traport\ rate \times MBSPVA - CC1\ initial\ backward}\,, \tag{126}$$

$$CC1_{PM} = C1_{PM} \times s_{PM}, \tag{127}$$

$$CC1_{CBC}^{B_G} = C1\,forward \times n_{CBC}^{B_G}, \tag{128}$$

$$CC1_{NSC}^{B_G} = C1\,forward \times n_{NSC}^{B_G}, \tag{129}$$

$$CC1_{GCC}^{B_G} = C1\,forward \times n_{GCC}^{B_G}, \tag{130}$$

$$CC1_{CBC}^{A_{PM}} = C1\,backward \times n_{CBC}^{A_{PM}}, \tag{131}$$

$$CC1_{NSC}^{A_{PM}} = C1\,backward \times n_{NSC}^{A_{PM}}, \tag{132}$$

$$CC1_{GCC}^{A_{PM}} = C1\,backward \times n_{GCC}^{A_{PM}}, \tag{133}$$

$$\begin{aligned} k\,CC1\,production &= (rate\ of\ accumulation\ of\ stationary\ nb_1\ NSCP \times C1\,forward \\ &+ rate\ of\ accumulation\ of\ stationary\ na_2\ NSCP \times C1\,backward) \\ &+ (rate\ of\ accumulation\ of\ stationary\ nb_1\ NSCD \times C1\,forward \\ &+ rate\ of\ accumulation\ of\ stationary\ na_2\ NSCD \times C1\,backward) \end{aligned} \tag{134}$$

$$C2_{PM} = \frac{CC2_{PM}}{s_{PM}}, \tag{135}$$

$$sc2a2 = \frac{CBSPVA \times \frac{C2_{PM}}{kc2a}}{\left(1 + \frac{C2_{PM}}{kc2a}\right)}, \tag{136}$$

$$C2\,backward = sc2a2, \tag{137}$$

$$CC2\,initial\,backward = C2\,backward \times initial\,back\,transport\,rate, \tag{138}$$

$$CC2\,final\,forward = CC2\,initial\,backward, \tag{139}$$

$$C2\,forward = \frac{CC2\,final\,forward}{final\,forward\,transport\,rate}, \tag{140}$$

$$C2\,initial\,forkward = C2\,forward \times initial\,forward\,transport\,rate, \tag{141}$$

$$C2_G = \frac{kc2b \times CC2\,initial\,forward}{initial\,forward\,traport\,rate \times MBSPVA - CC2\,initial\,forward}, \tag{142}$$

$$CC2_G = C2_G \times s_G, \tag{143}$$

$$CC2_{CBC}^{B_G} = C2\,forward \times n_{CBC}^{B_G}, \tag{144}$$

$$CC2_{NSC}^{B_G} = C2\,forward \times n_{NSC}^{B_G}, \quad (145)$$

$$CC2_{GCC}^{B_G} = C2\,forward \times n_{GCC}^{B_G}, \quad (146)$$

$$CC2_{CBC}^{A_{PM}} = C2\,backward \times n_{CBC}^{A_{PM}}, \quad (147)$$

$$CC2_{NSC}^{A_{PM}} = C2\,backward \times n_{NSC}^{A_{PM}}, \quad (148)$$

$$CC2_{GCC}^{A_{PM}} = C2\,backward \times n_{GCC}^{A_{PM}}, \quad (149)$$

$$\begin{aligned} k\,CC2\,production \\ &= (rate\ of\ accumulation\ of\ stationary\ nb_1\ NSCP \times C2\,forward \\ &+\ rate\ of\ accumulation\ of\ stationary\ na_2\ NSCP \times C2\,backward) \\ &+ (rate\ of\ accumulation\ of\ stationary\ nb_1\ NSCD \times C2\,forward \\ &+ rate\ of\ accumulation\ of\ stationary\ na_2\ NSCD\ \times C2\,backward) \quad (150) \end{aligned}$$

We have assumed a fixed effective tubulin concentration 9 µM and used Margolin model (Margolin et al., 2012) for simulations of dynamics microtubules for a range of hydrolysis rates (0.60 $sec^{-1}$ – 0.80 $sec^{-1}$). To find the relationship between hydrolysis rate and average dynamic MT length, degradation rate of dynamic MT, axonal growth velocity. We assumed fixed Nucleation rate (65 $\min^{-1}$) and Stabilization rate (0.015 65 $\min^{-1}$) and used power fitting method to calculated fitting parameters values.

$$\left.\begin{aligned} a &= 0.7497 \\ b &= -0.07383 \\ c &= 0.091 \\ d &= -7.256 \\ e &= 13.35 \\ f &= 6.287 \end{aligned}\right\} \quad (151)$$

Consider an axonal growth velocity ‘v’ and hydrolysis rate ‘x’.

$$hydrolysis\ rate\ (x) = av^b \quad (152)$$

$$Average\ dyn\ MT\ length\ = cx^d \quad (153)$$

$$degradation\,rate\ = ex^f \quad (154)$$

$$\frac{d\ dyn_{MT}}{dt} = nucleation\ rate - (stabilization\ rate + degradation\ rate) \times dyn_{MT} \quad (155)$$

$$\frac{d\ stable_{MT\ length}}{dt} = \frac{stabilization\ rate \times dyn_{MT} \times Average\ dyn\ MT\ length}{Mts\ per\ crosssection} \quad (156)$$

$$dynamic_{MT\ length} = \frac{dyn_{MT} \times Average\ dyn\ MT\ length}{Mts\ per\ crosssection} \quad (157)$$

$$microtubule\ bundle\ length = dynamic_{MT\ length} + stable_{MT\ length} \quad (158)$$

**Supp Table 1: Parameters and Initial Values** This parameter set allows growth at 2 µm/h

| Parameter | Value | Units | Reference |
|---|---|---|---|
| Number of microtubules per cross-section | 20<br>(10 – 100) | # | (Fadic et al., 1985) |
| Diameter of neurite | 1.0<br>(1.0 – 3.0) | µm | (Harris and Stevens, 1989) |
| Length of growth cones | 10 | µm | (Beller et al., 2013) |
| Kinesin motor protein anterograde velocity | 1 | µm/s | (Vale et al., 1996)<br>(Carter and Cross, 2005) |
| Dynein motor protein retrograde velocity | 0.65 | µm/s | (King and Schroer, 2000)<br>(Nishiura et al., 2004) |
| Vesicle diameter | 39.7 ± 6.6 | nm | (Zhang et al., 1998) |
| Growth cone surface area | 70 - 200 | $\mu m^2$ | (Kunda et al., 2001) |
| Dissociation constant of SNARE SX with coat protein B. ($k_{sx}^{A}$ ) | 1.00 | Molecules | Heinrich and Rapaport |
| Dissociation constant of SNARE SU with coat protein B. ($k_{su}^{A}$ ) | 1.00 | Molecules | Heinrich and Rapaport |
| Dissociation constant of SNARE SX with coat protein B. ($k_{sx}^{B}$ ) | 10000 | Molecules | Heinrich and Rapaport |
| Dissociation constant of SNARE SU with coat protein B. ($k_{su}^{B}$ ) | 100 | Molecules | Heinrich and Rapaport |
| Dissociation constant of SNARE SY with coat protein B. ($k_{sy}^{A}$ ) | 10000 | Molecules | Heinrich and Rapaport |
| Dissociation constant of SNARE SV with coat protein B. ($k_{sv}^{A}$ ) | 100 | Molecules | Heinrich and Rapaport |
| Dissociation constant of SNARE SY with coat protein B. ($k_{sy}^{B}$ ) | 1.00 | Molecules | Heinrich and Rapaport |
| Dissociation constant of SNARE SV with coat protein B. ($k_{sv}^{B}$ ) | 1.00 | Molecules | Heinrich and Rapaport |

| | | | |
|---|---|---|---|
| Dissociation constants of recruitment factor 1 from coat protein A ($k_{r_1}^{A}$) | 1.00 | Molecules | Based on Henrich and Rappaport model |
| Dissociation constants of recruitment factor 1 from coat protein A ($k_{r_1}^{B}$) | 10000 | Molecules | Based on Henrich and Rappaport model |
| Dissociation constants of recruitment factor 2 from coat protein A ($k_{r_2}^{A}$) | 10000 | Molecules | Based on Henrich and Rappaport model |
| Dissociation constants of recruitment factor 2 from coat protein A ($k_{r_2}^{B}$) | 1.00 | Molecules | Based on Henrich and Rappaport model |
| Cargo binding spots per vesicle surface area | 2 | # | |
| Dissociation constants of motor kinesin $kin_G$ from coat protein A ($k_{kin}^{A}$) | 10 | Molecules | Based on Henrich and Rappaport model |
| Dissociation constants of motor kinesin $kin_G$ from coat protein B ($k_{kin}^{B}$) | 0.1 | Molecules | Based on Henrich and Rappaport model |
| Dissociation constants of motor Dynein $Dyn_G$ from coat protein B ($k_{dyn}^{A}$) | 0.1 | Molecules | Based on Henrich and Rappaport model |
| Dissociation constants of motor Dynein $Dyn_G$ from coat protein B ($k_{dyn}^{B}$) | 10 | Molecules | Based on Henrich and Rappaport model |
| Length of CBC compartment | 1 | µm | Assumed based on initial neurite length |
| Length of GCC compartment | 2 | µm | Assumed based on initial neurite length |
| Length of NSC compartment | 20 | µm | Assumed based on initial neurite length |
| Scale parameter multiplier ($a_1$) | 3.146e-12 | | Predicted from (Margolin et al., 2012) |
| Scale parameter exponent ($a_2$) | 1.0311 | | Predicted from (Margolin et al., 2012) |
| Scale parameter multiplier ($b_1$) | 8.7342e-19 | | Predicted from (Margolin et al., 2012) |
| Scale parameter exponent ($b_2$) | 4.2122 | | Predicted from (Margolin et al., 2012) |
| Shape multiplier (m) | 0.02784 | | Predicted from (Margolin et al., 2012) |
| Shape constant (b) | 0.15547 | | Predicted from (Margolin et al., 2012) |
| Degradation multiplier (d1) | -12.78 | | Predicted from (Margolin et al., 2012) |
| Degradation exponent (d2) | 2.93e10 | | Predicted from (Margolin et al., 2012) |
| Anterograde vesicle surface area | 0.05 | $\mu m^2$ | (Zhang et al., 1998) |
| Retrograde vesicle surface area | 0.05 | $\mu m^2$ | (Zhang et al., 1998) |
| Snare binding spots per vesicle surface area | 1400 | # | Predicted |

| | | | |
|---|---|---|---|
| Motor binding spots per vesicle surface area | 140 | # | Predicted |
| SNARE Y production rate at TNG | 3.4372 | $min^{-1}$ | Predicted |
| SNARE V production rate at TNG | 0.74401 | $min^{-1}$ | Predicted |
| SNARE X production rate at TNG | 9.4813 | $min^{-1}$ | Predicted |
| SNARE U production rate at TNG | 774.5352 | $min^{-1}$ | Predicted |
| Motor K production rate at TNG | 14.9856 | $min^{-1}$ | Predicted |
| Motor D production rate at TNG | 12.3926 | $min^{-1}$ | Predicted |
| Recruitment factor 1 production rate at TNG | 2.9971e-05 | $min^{-1}$ | Predicted |
| Recruitment factor 2 production rate at TNG | 2.4785e-05 | $min^{-1}$ | Predicted |
| Vesicles budding rate at TGN with coat A ( $w_G^A$) | 1.0009e-05 | $\mu m^2$ | Predicted |
| Vesicles budding rate at Plasma Membrane with coat A ( $w_{PM}^A$) | 0.00010009 | $\mu m^2$ | Predicted |
| Vesicles budding rate at TGN with coat B ( $w_G^B$) | 0.00027089 | $\mu m^2$ | Predicted |
| Vesicles budding rate at Plasma Membrane with coat A ( $w_{PM}^B$) | 2.7078e-05 | $\mu m^2$ | Predicted |
| Tethering rate constant of vesicles (X, U) to target compartment Golgi ($\kappa_{XU_G}$) | 2e-06 | 1/ (molecules min) | Predicted |
| Tethering rate constant of vesicles (X, U) to target compartment Growth Cone Plasma Membrane ($\kappa_{XU_{PM}}$) | 2e-07 | 1/ (molecules min) | Predicted |
| Tethering rate constant of vesicles (Y, V) to target compartment Golgi ($\kappa_{YV_G}$) | 5e-06 | 1/ (molecules min) | Predicted |
| Tethering rate constant of vesicles (Y, V) to target compartment Growth Cone Plasma Membrane ($\kappa_{YV_{PM}}$) | 5e-05 | 1/ (molecules min) | Predicted |
| Required snare complex per vesicle fusion ($R_{SCPVF}$ ) | 5.00 | # | Predicted |
| Fraction of bound kinesin motor with microtubules in CBC compartment | 95 | percent | Assumed based on (Dixit at al., 2008, LaPointe et al., 2009) |
| Fraction of bound kinesin motor with microtubules in NSC-P compartment | 95 | Percent | Assumed based on (Dixit at al., 2008, LaPointe et al., 2009) |

| | | | |
|---|---|---|---|
| Fraction of bound kinesin motor with microtubules in NSC-D compartment | 0.022 | Percent | Assumed based on (Dixit at al., 2008, LaPointe et al., 2009) |
| Fraction of bound kinesin motor with microtubules in GCC compartment | 1.0 | Percent | Assumed based on (Dixit at al., 2008, LaPointe et al., 2009) |
| Fraction of bound dynein motor with microtubules in CBC compartment | 1.0 | Percent | Assumed based on (Dixit at al., 2008, LaPointe et al., 2009) |
| Fraction of bound dynein motor with microtubules in NSC-P compartment | 90 | Percent | Assumed based on (Dixit at al., 2008, LaPointe et al., 2009) |
| Fraction of bound dynein motor with microtubules in NSC-D compartment | 90 | Percent | Assumed based on (Dixit at al., 2008, LaPointe et al., 2009) |
| Fraction of bound dynein motor with microtubules in CBC compartment | 95 | percent | Assumed based on (Dixit at al., 2008, LaPointe et al., 2009) |
| Nucleation rate of microtubules | 65 | 1/min | Predicted from (Margolin et al., 2012) |
| Membrane production rate at TGN | 0.8539 | 1/min | Predicted |
| Stabilization rate of dynamic microtubules | 0.015 | 1/min | Predicted |
| Degradation rate of dynamic MTs | 1.5819 | 1/min | Predicted from (Margolin et al., 2012) |
| **Initial Values** | | | |
| Trans Golgi Network (TGN) | 50 | $\mu m^2$ | Predicted |
| Coat B budded vesicles membrane surface area from TGN in CBC | 0.03 | $\mu m^2$ | Predicted |
| Coat B budded vesicles membrane surface area from TGN in NSC-P | 0.10617 | $\mu m^2$ | Predicted |
| Coat B budded vesicles membrane surface area from NSC-P in NSC-D | 449.0088 | $\mu m^2$ | Predicted |
| Coat B budded vesicles membrane surface area from TGN in GCC | 60.5195 | $\mu m^2$ | Predicted |
| Growth cone plasma membrane | 50 | $\mu m^2$ | Predicted |
| Coat A budded vesicles membrane surface area from TGN in CBC | 2.2911e-06 | $\mu m^2$ | Predicted |
| Coat A budded vesicles membrane surface area from TGN in NSC | 0.0008038 | $\mu m^2$ | Predicted |

| | | | |
|---|---|---|---|
| | | $\mu m^2$ | |
| Coat A budded vesicles membrane surface area from TGN in GCC | 0.0067794 | | Predicted |
| Coat B budded vesicles membrane surface area from plasma membrane in CBC | 0.01252 | $\mu m^2$ | Predicted |
| Coat B budded vesicles membrane surface area from plasma membrane in NSC-P | 6.5014e-05 | $\mu m^2$ | Predicted |
| Coat B budded vesicles membrane surface area from plasma membrane in NSC-D | 6.5014e-05 | $\mu m^2$ | Predicted |
| Coat B budded vesicles membrane surface area from plasma membrane in GCC | 4.0126e-05 | $\mu m^2$ | Predicted |
| Coat A budded vesicles membrane surface area from plasma membrane in CBC | 1.000 | $\mu m^2$ | Predicted |
| Coat A budded vesicles membrane surface area from plasma membrane in NSC-P | 0.14245 | $\mu m^2$ | Predicted |
| Coat A budded vesicles membrane surface area from plasma membrane in NSC-D | 0.14245 | $\mu m^2$ | Predicted |
| Coat A budded vesicles membrane surface area from plasma membrane in GCC | 0.2 | $\mu m^2$ | Predicted |
| Neurite shaft surface area | 62.8319 | $\mu m^2$ | Predicted |
| SNAREs Y in TGN compartment | 5.7375 | molecules | Predicted |
| Total vesicle snare Y with Coat B budded from TNG in CBC | 0.13762 | molecules | Predicted |
| Total vesicle snare Y with Coat B budded from TNG in NSC-P | 0.48706 | Molecules | Predicted |
| Total vesicle snare Y with Coat B budded from TNG in NSC-D | 2059.7539 | Molecules | Predicted |
| Total vesicle snare Y with Coat B budded from TNG in GCC | 277.6231 | Molecules | Predicted |
| SNAREs Y in GC compartment | 20141.5989 | Molecules | Predicted |

| | | | |
|---|---|---|---|
| Total vesicle snare Y with Coat A budded from TNG in CBC | 1.9228e-11 | Molecules | Predicted |
| Total vesicle snare Y with Coat A budded from TNG in NSC-P | 6.746e-09 | Molecules | Predicted |
| Total vesicle snare Y with Coat A budded from TNG in NSC-D | 6.746e-09 | Molecules | Predicted |
| Total vesicle snare Y with Coat A budded from TNG in GCC | 0.00011986 | Molecules | Predicted |
| Total vesicle snare Y with Coat B budded from GC in CBC | 10.449 | Molecules | Predicted |
| Total vesicle snare Y with Coat B budded from GC in NSC-P | 0.054339 | Molecules | Predicted |
| Total vesicle snare Y with Coat B budded from GC in NSC-D | 0.054339 | Molecules | Predicted |
| Total vesicle snare Y with Coat B budded from GC in GCC | 0.033537 | Molecules | Predicted |
| Total vesicle snare Y with Coat A budded from GC in CBC | 5.5472 | Molecules | Predicted |
| Total vesicle snare Y with Coat A budded from GC in NSC-P | 0.7902 | Molecules | Predicted |
| Total vesicle snare Y with Coat A budded from GC in NSC-D | 0.7902 | Molecules | Predicted |
| Total vesicle snare Y with Coat A budded from GC in GCC | 1.1094 | Molecules | Predicted |
| SNAREs V in TGN compartment | 0.14 | Molecules | Predicted |
| Total vesicle snare V with Coat B budded from TNG in CBC | 0.029789 | Molecules | Predicted |
| Total vesicle snare V with Coat B budded from TNG in NSC-P | 0.10543 | Molecules | Predicted |
| Total vesicle snare V with Coat B budded from TNG in NSC-D | 445.8522 | Molecules | Predicted |

| | | | |
|---|---|---|---|
| Total vesicle snare V with Coat B budded from TNG in GCC | 60.094 | Molecules | Predicted |
| SNAREs V in GC compartment | 43.5983 | Molecules | Predicted |
| Total vesicle snare V with Coat A budded from TNG in CBC | 1.2923e-07 | Molecules | Predicted |
| Total vesicle snare V with Coat A budded from TNG in NSC-P | 4.5337e-05 | molecules | Predicted |
| Total vesicle snare V with Coat A budded from TNG in NSC-D | 4.5337e-05 | molecules | Predicted |
| Total vesicle snare V with Coat A budded from TNG in GCC | 0.00049928 | Molecules | Predicted |
| Total vesicle snare V with Coat B budded from GC in CBC | 7.0517 | Molecules | Predicted |
| Total vesicle snare V with Coat B budded from GC in NSC-P | 0.036543 | Molecules | Predicted |
| Total vesicle snare V with Coat B budded from GC in NSC-D | 0.036543 | Molecules | Predicted |
| Total vesicle snare V with Coat B budded from GC in GCC | 0.022554 | Molecules | Predicted |
| Total vesicle snare V with Coat A budded from GC in CBC | 1.2007 | Molecules | Predicted |
| Total vesicle snare V with Coat A budded from GC in NSC-P | 0.17105 | Molecules | Predicted |
| Total vesicle snare V with Coat A budded from GC in NSC-D | 0.17105 | Molecules | Predicted |
| Total vesicle snare V with Coat A budded from GC in GCC | 0.24015 | Molecules | Predicted |
| SNAREs X in TGN compartment | 19990.5187 | Molecules | Predicted |
| Total vesicle snare X with Coat B budded from TNG in CBC | 0.37962 | Molecules | Predicted |

| | | | |
|---|---|---|---|
| Total vesicle snare X with Coat B budded from TNG in NSC-P | 1.3435 | Molecules | Predicted |
| Total vesicle snare X with Coat B budded from TNG in NSC-D | 5681.6884 | Molecules | Predicted |
| Total vesicle snare X with Coat B budded from TNG in GCC | 765.8041 | Molecules | Predicted |
| SNARE X in GC compartment | 2.0142 | Molecules | Predicted |
| Total vesicle snare X with Coat A budded from TNG in CBC | 0.0019236 | Molecules | Predicted |
| Total vesicle snare X with Coat A budded from TNG in NSC-P | 0.67487 | Molecules | Predicted |
| Total vesicle snare X with Coat A budded from TNG in NSC-D | 0.67487 | Molecules | Predicted |
| Total vesicle snare X with Coat A budded from TNG in GCC | 5.6971 | Molecules | Predicted |
| Total vesicle snare X with Coat B budded from GC in CBC | 0.00048599 | molecules | Predicted |
| Total vesicle snare X with Coat B budded from GC in NSC-P | 5.4461e-10 | Molecules | Predicted |
| Total vesicle snare X with Coat B budded from GC in NSC-D | 5.4461e-10 | Molecules | Predicted |
| Total vesicle snare X with Coat B budded from GC in GCC | 3.3613e-10 | Molecules | Predicted |
| Total vesicle snare X with Coat A budded from GC in CBC | 15.3015 | Molecules | Predicted |
| Total vesicle snare X with Coat A budded from GC in NSC-P | 2.1797 | Molecules | Predicted |
| Total vesicle snare X with Coat A budded from GC in NSC-D | 2.1797 | Molecules | Predicted |

| | | | |
|---|---|---|---|
| Total vesicle snare X with Coat A budded from GC in GCC | 3.0603 | Molecules | Predicted |
| SNAREs U in TGN compartment | 15563.6828 | Molecules | Predicted |
| Total vesicle snare U with Coat B budded from TNG in CBC | 31.0112 | Molecules | Predicted |
| Total vesicle snare U with Coat B budded from TNG in NSC-P | 109.7533 | Molecules | Predicted |
| Total vesicle snare U with Coat B budded from TNG in NSC-D | 464143.3168 | Molecules | Predicted |
| Total vesicle snare U with Coat B budded from TNG in GCC | 62559.3683 | Molecules | Predicted |
| SNARE U in GC compartment | 453.8692 | Molecules | Predicted |
| Total vesicle snare U with Coat A budded from TNG in CBC | 0.001279 | Molecules | Predicted |
| Total vesicle snare U with Coat A budded from TNG in NSC-P | 0.44874 | Molecules | Predicted |
| Total vesicle snare U with Coat A budded from TNG in NSC-D | 0.44874 | Molecules | Predicted |
| Total vesicle snare U with Coat A budded from TNG in GCC | 3.7795 | Molecules | Predicted |
| Total vesicle snare U with Coat B budded from GC in CBC | 0.0011828 | Molecules | Predicted |
| Total vesicle snare U with Coat B budded from GC in NSC-P | 3.6171e-06 | Molecules | Predicted |
| Total vesicle snare U with Coat B budded from GC in NSC-D | 3.6171e-06 | Molecules | Predicted |
| Total vesicle snare U with Coat B budded from GC in GCC | 2.2324e-06 | Molecules | Predicted |
| Total vesicle snare U with Coat A budded from GC in CBC | 1250 | Molecules | Predicted |

| | | | |
|---|---|---|---|
| Total vesicle snare U with Coat A budded from GC in NSC-P | 178.0627 | Molecules | Predicted |
| Total vesicle snare U with Coat A budded from GC in NSC-D | 178.0627 | Molecules | Predicted |
| Total vesicle snare U with Coat A budded from GC in GCC | 250 | Molecules | Predicted |
| Kinesin receptors K in TGN compartment | 0.83333 | Molecules | Predicted |
| Total kinesin receptors K with Coat B budded from TNG in CBC | 0.6 | Molecules | Predicted |
| Total kinesin receptors K with Coat B budded from TNG in NSC-P | 2.1235 | Molecules | Predicted |
| Total kinesin receptors K with Coat B budded from TNG in NSC-D | 8980.1762 | Molecules | Predicted |
| Total kinesin receptors K with Coat B budded from TNG in GCC | 1210.3894 | Molecules | Predicted |
| kinesin receptors K in GC compartment | 104.4112 | Molecules | Predicted |
| Total kinesin receptors K with Coat A budded from TNG in CBC | 7.0251e-07 | Molecules | Predicted |
| Total kinesin receptors K with Coat A budded from TNG in NSC-P | 0.00024645 | Molecules | Predicted |
| Total kinesin receptors K with Coat A budded from TNG in NSC-D | 0.00024645 | Molecules | Predicted |
| Total kinesin receptors K with Coat A budded from TNG in GCC | 0.0020644 | Molecules | Predicted |
| Total kinesin receptors K with Coat B budded from GC in CBC | 1.6756 | Molecules | Predicted |
| Total kinesin receptors K with Coat B budded from GC in NSC-P | 0.0087054 | Molecules | Predicted |
| Total kinesin receptors K with Coat B budded from GC in NSC-D | 0.0087054 | Molecules | Predicted |

| | | | |
|---|---|---|---|
| Total kinesin receptors K with Coat B budded from GC in GCC | 0.0053728 | Molecules | Predicted |
| Total kinesin receptors K with Coat A budded from GC in CBC | 24.1848 | Molecules | Predicted |
| Total kinesin receptors K with Coat A budded from GC in NSC-P | 3.4451 | Molecules | Predicted |
| Total kinesin receptors K with Coat A budded from GC in NSC-D | 3.4451 | Molecules | Predicted |
| Total kinesin receptors K with Coat A budded from GC in GCC | 4.837 | Molecules | Predicted |
| Dynein receptor D in TGN compartment | 66.9821 | Molecules | Predicted |
| Total dynein receptor D with Coat B budded from TNG in CBC | 0.49618 | Molecules | Predicted |
| Total dynein receptor D with Coat B budded from TNG in NSC-P | 1.7561 | Molecules | Predicted |
| Total dynein receptor D with Coat B budded from TNG in NSC-D | 7426.2931 | Molecules | Predicted |
| Total dynein receptor D with Coat B budded from TNG in GCC | 1000.9499 | Molecules | Predicted |
| Dynein receptor D in GC compartment | 0.83333 | Molecules | Predicted |
| Total dynein receptor D with Coat A budded from TNG in CBC | 0.00030677 | Molecules | Predicted |
| Total dynein receptor D with Coat A budded from TNG in NSC-P | 0.10763 | Molecules | Predicted |
| Total dynein receptor D with Coat A budded from TNG in NSC-D | 0.10763 | Molecules | Predicted |
| Total dynein receptor D with Coat A budded from TNG in GCC | 0.90727 | Molecules | Predicted |
| Total dynein receptor D with Coat B budded from GC in CBC | 0.0038582 | Molecules | Predicted |

| | | | |
|---|---|---|---|
| Total dynein receptor D with Coat B budded from GC in NSC-P | 1.9925e-05 | Molecules | Predicted |
| Total dynein receptor D with Coat B budded from GC in NSC-D | 1.9925e-05 | Molecules | Predicted |
| Total dynein receptor D with Coat B budded from GC in GCC | 1.2297e-05 | Molecules | Predicted |
| Total dynein receptor D with Coat A budded from GC in CBC | 20 | Molecules | Predicted |
| Total dynein receptor D with Coat A budded from GC in NSC-P | 2.849 | Molecules | Predicted |
| Total dynein receptor D with Coat A budded from GC in NSC-D | 2.849 | Molecules | Predicted |
| Total dynein receptor D with Coat A budded from GC in GCC | 4 | Molecules | Predicted |
| Recruitment factor 1 in TGN compartment | 100 | Molecules | Predicted |
| Total recruitment factor 1 with Coat B budded from TNG in CBC | 1.2e-06 | Molecules | Predicted |
| Total recruitment factor 1 with Coat B budded from TNG in NSC-P | 4.2469e-06 | Molecules | Predicted |
| Total recruitment factor 1 with Coat B budded from TNG in NSC-D | 0.01796 | Molecules | Predicted |
| Total recruitment factor 1 with Coat B budded from TNG in GCC | 0.00242 | Molecules | Predicted |
| recruitment factor 1 in GC compartment | 0.0012092 | Molecules | Predicted |
| Total recruitment factor 1 with Coat A budded from TNG in CBC | 3.0544e-05 | Molecules | Predicted |
| Total recruitment factor 1 with Coat A budded from TNG in NSC-P | 0.010716 | Molecules | Predicted |
| Total recruitment factor 1 with Coat A budded from TNG in NSC-D | 0.010716 | Molecules | Predicted |

| | | | |
|---|---|---|---|
| Total recruitment factor 1 with Coat A budded from TNG in GCC | 0.090358 | Molecules | Predicted |
| recruitment factor 1 in GC-PM compartment | 0.0012092 | Molecules | Predicted |
| Total recruitment factor 1 with Coat A budded from GC in CBC | 4.8369e-05 | Molecules | Predicted |
| Total recruitment factor 1 with Coat A budded from GC in NSC-P | 6.8901e-06 | Molecules | Predicted |
| Total recruitment factor 1 with Coat A budded from GC in NSC-D | 6.8901e-06 | Molecules | Predicted |
| Total recruitment factor 1 with Coat A budded from GC in GCC | 9.6737e-06 | Molecules | Predicted |
| Total recruitment factor 1 with Coat B budded from GC in CBC | 7.4581e-11 | Molecules | Predicted |
| Total recruitment factor 1 with Coat B budded from GC in NSC-P | 3.6518e-13 | Molecules | Predicted |
| Total recruitment factor 1 with Coat B budded from GC in NSC-D | 3.6518e-13 | Molecules | Predicted |
| Total recruitment factor 1 with Coat B budded from GC in GCC | 2.2539e-13 | Molecules | Predicted |
| Recruitment factor 2 in TGN compartment | 0.00082696 | Molecules | Predicted |
| Total recruitment factor 2 with Coat B budded from TNG in CBC | 9.9234e-07 | Molecules | Predicted |
| Total recruitment factor 2 with Coat B budded from TNG in NSC-P | 3.512e-06 | Molecules | Predicted |
| Total recruitment factor 2 with Coat B budded from TNG in NSC-D | 0.014852 | Molecules | Predicted |
| Total recruitment factor 2 with Coat B budded from TNG in GCC | 0.0020019 | Molecules | Predicted |
| Recruitment factor 2 in GC compartment | 100 | Molecules | Predicted |

| | | | |
|---|---|---|---|
| Total recruitment factor 2 with Coat A budded from TNG in CBC | 1.1325e-14 | Molecules | Predicted |
| Total recruitment factor 2 with Coat A budded from TNG in NSC-P | 3.9661e-12 | Molecules | Predicted |
| Total recruitment factor 2 with Coat A budded from TNG in NSC-D | 3.9661e-12 | Molecules | Predicted |
| Total recruitment factor 2 with Coat A budded from TNG in GCC | 3.829e-11 | Molecules | Predicted |
| Total recruitment factor 2 with Coat B budded from GC in CBC | 0.16687 | Molecules | Predicted |
| Total recruitment factor 2 with Coat B budded from GC in NSC-P | 0.00086636 | Molecules | Predicted |
| Total recruitment factor 2 with Coat B budded from GC in NSC-D | 0.00086636 | Molecules | Predicted |
| Total recruitment factor 2 with Coat B budded from GC in GCC | 0.0005347 | Molecules | Predicted |
| Total recruitment factor 2 with Coat A budded from GC in CBC | 3.9999e-05 | Molecules | Predicted |
| Total recruitment factor 2 with Coat A budded from GC in NSC-P | 5.6979e-06 | Molecules | Predicted |
| Total recruitment factor 2 with Coat A budded from GC in NSC-D | 5.6979e-06 | Molecules | Predicted |
| Total recruitment factor 2 with Coat A budded from GC in GCC | 7.9998e-06 | Molecules | Predicted |
| Effective tubulin | 9 | µM | Gardner et al., 2011 |
| No of dynamic MTs in neurite shaft | 20 | | Yu and Baas, 1994) |
| Initial total length of stable MTs length | 15 | µm | Assumed initial MTB length initial neurite length |

For other velocities parameter values can be calculated using the equations described above. Matlab script for these calculations is available upon request.

#### Supplementary Figure - 10

**A1**

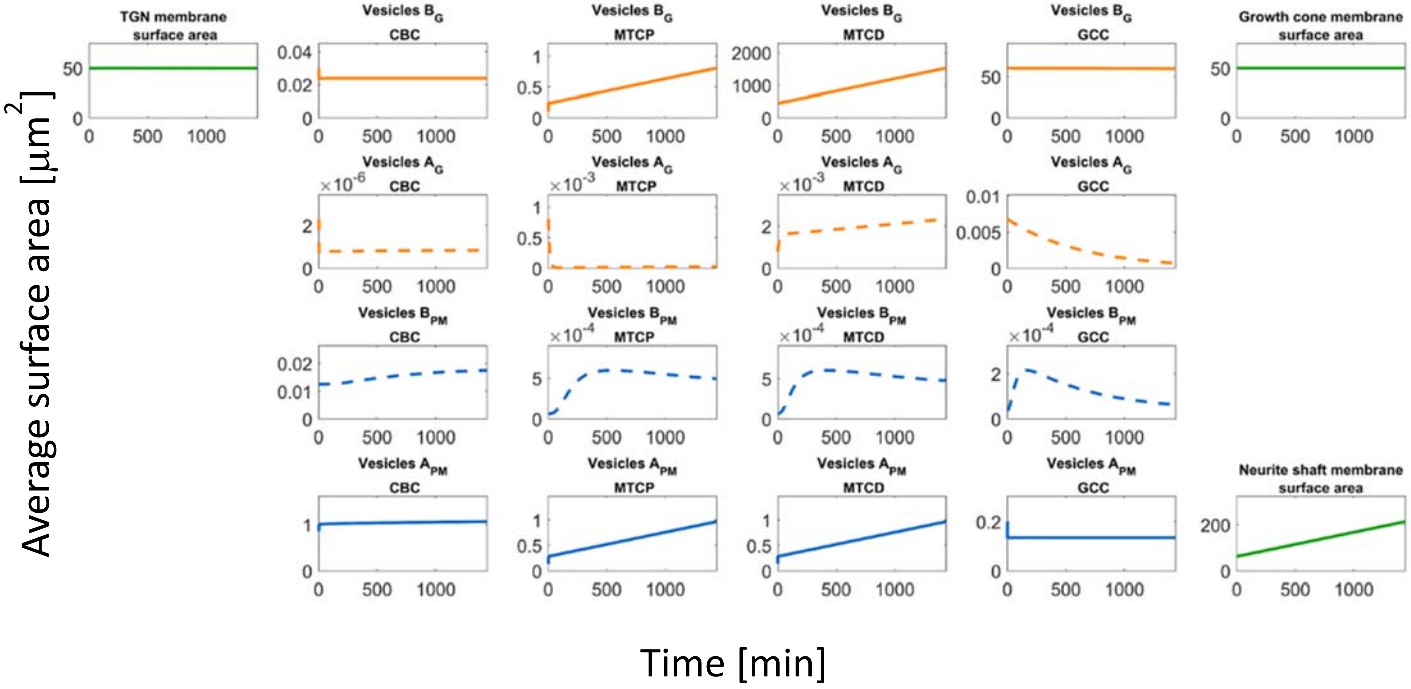


**A2**

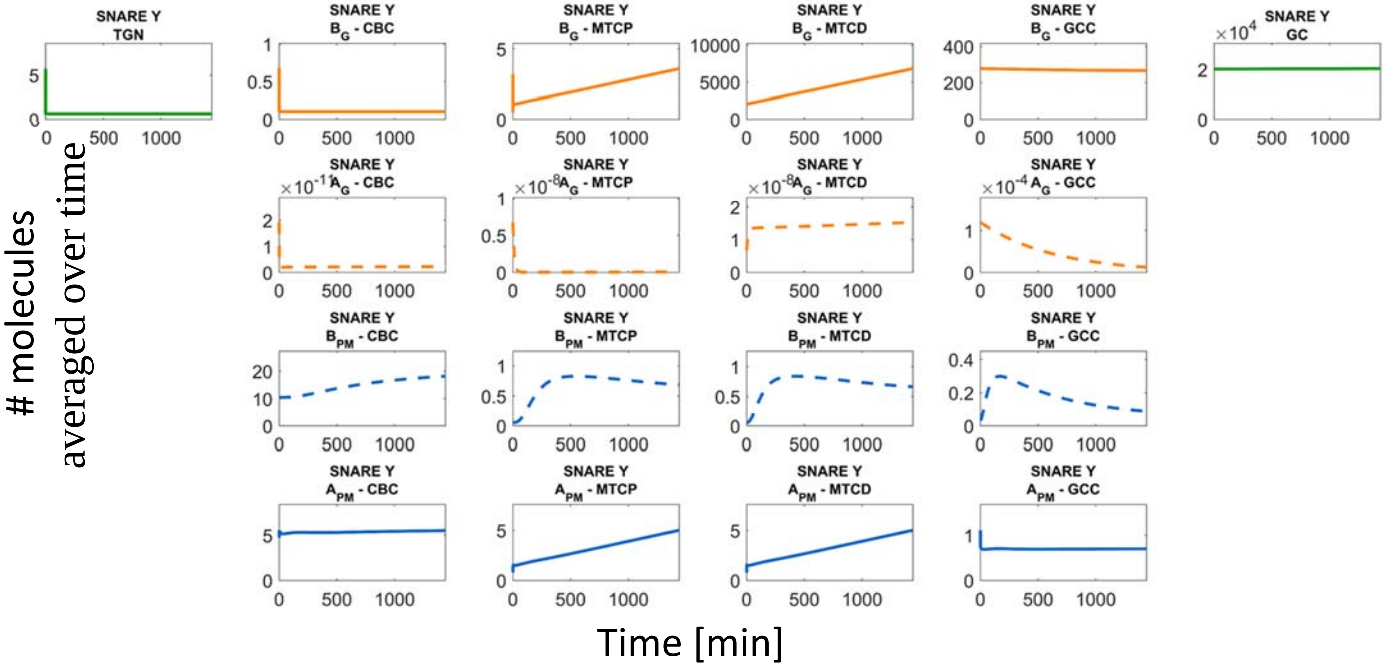

A3

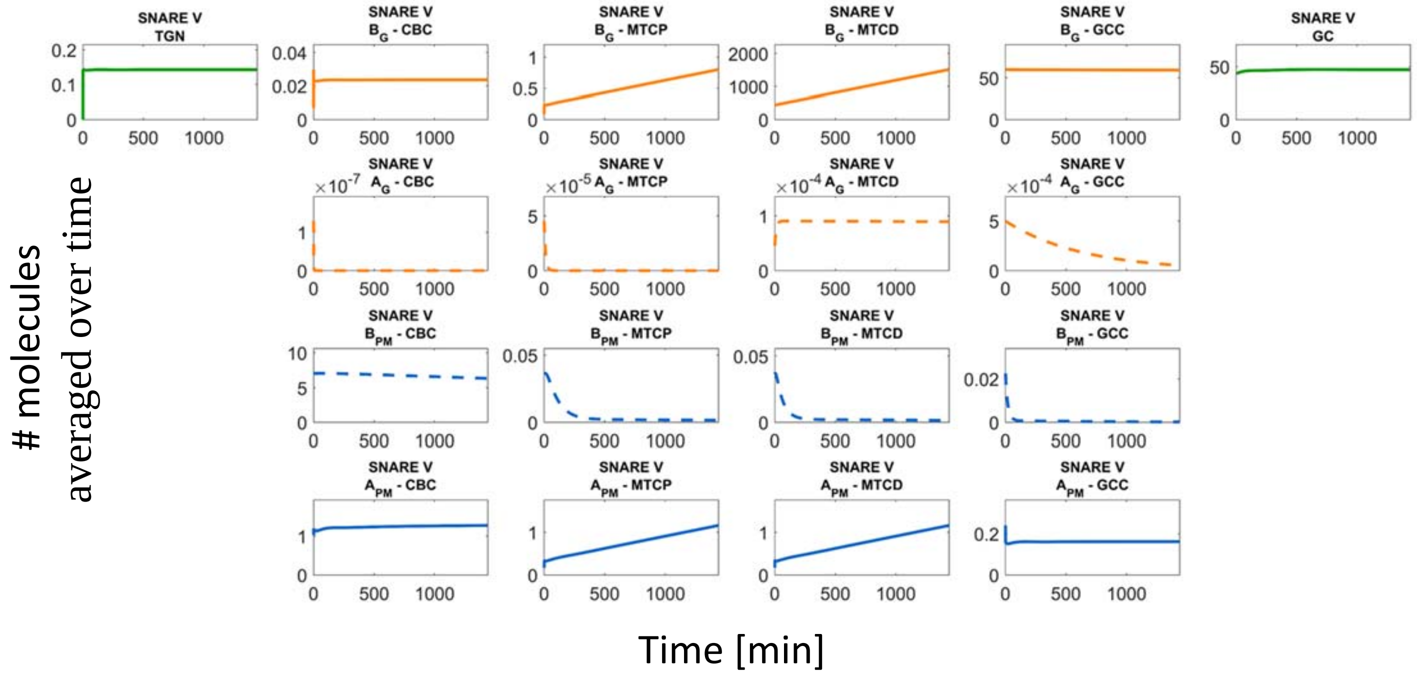
SNARE V TGN
SNARE V $B_G$ - CBC
SNARE V $B_G$ - MTCP
SNARE V $B_G$ - MTCD
SNARE V $B_G$ - GCC
SNARE V GC
SNARE V $A_G$ - CBC
SNARE V $A_G$ - MTCP
SNARE V $A_G$ - MTCD
SNARE V $A_G$ - GCC
SNARE V $B_{PM}$ - CBC
SNARE V $B_{PM}$ - MTCP
SNARE V $B_{PM}$ - MTCD
SNARE V $B_{PM}$ - GCC
SNARE V $A_{PM}$ - CBC
SNARE V $A_{PM}$ - MTCP
SNARE V $A_{PM}$ - MTCD
SNARE V $A_{PM}$ - GCC
### molecules averaged over time
Time [min]

A4

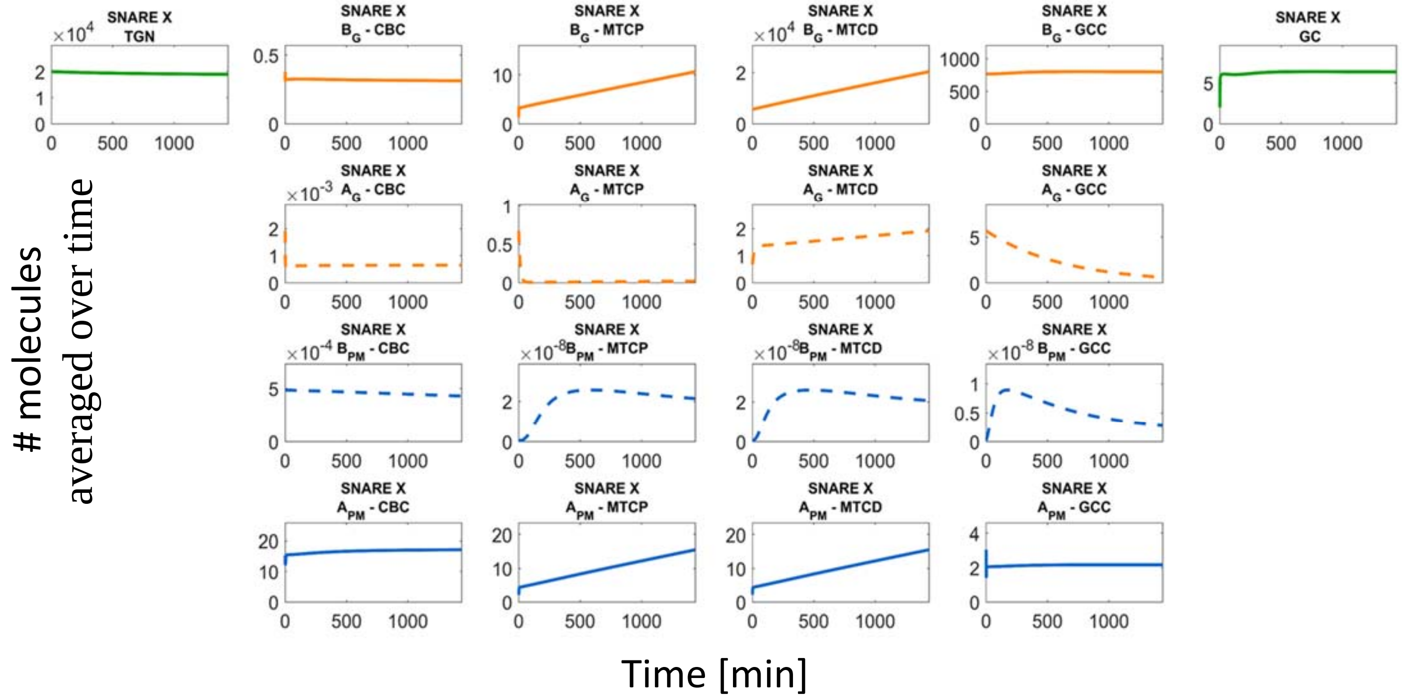
SNARE X TGN
SNARE X $B_G$ - CBC
SNARE X $B_G$ - MTCP
SNARE X $B_G$ - MTCD
SNARE X $B_G$ - GCC
SNARE X GC
SNARE X $A_G$ - CBC
SNARE X $A_G$ - MTCP
SNARE X $A_G$ - MTCD
SNARE X $A_G$ - GCC
SNARE X $B_{PM}$ - CBC
SNARE X $B_{PM}$ - MTCP
SNARE X $B_{PM}$ - MTCD
SNARE X $B_{PM}$ - GCC
SNARE X $A_{PM}$ - CBC
SNARE X $A_{PM}$ - MTCP
SNARE X $A_{PM}$ - MTCD
SNARE X $A_{PM}$ - GCC
### molecules averaged over time
Time [min]

**A5**

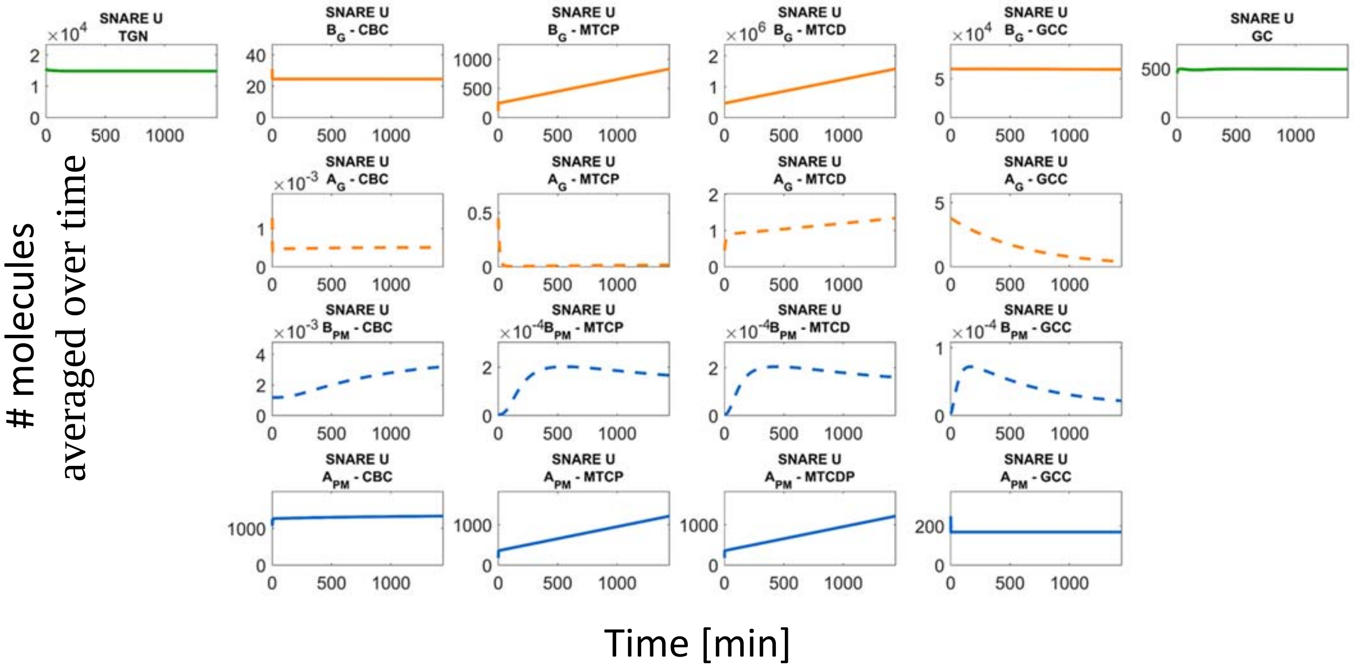
SNARE U TGN
SNARE U BG - CBC
SNARE U BG - MTCP
SNARE U BG - MTCD
SNARE U BG - GCC
SNARE U GC
SNARE U AG - CBC
SNARE U AG - MTCP
SNARE U AG - MTCD
SNARE U AG - GCC
SNARE U BPM - CBC
SNARE U BPM - MTCP
SNARE U BPM - MTCD
SNARE U BPM - GCC
SNARE U APM - CBC
SNARE U APM - MTCP
SNARE U APM - MTCDP
SNARE U APM - GCC
### molecules averaged over time
Time [min]

**A6**

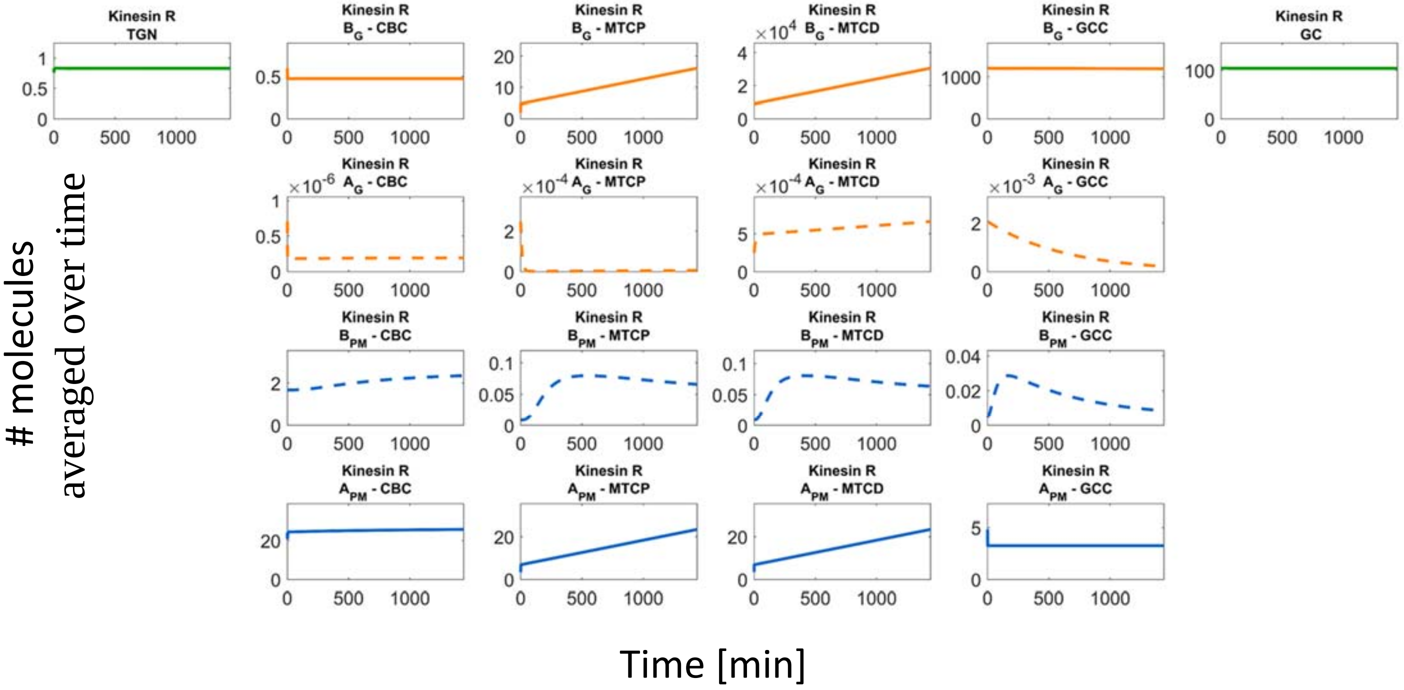
Kinesin R TGN
Kinesin R BG - CBC
Kinesin R BG - MTCP
Kinesin R BG - MTCD
Kinesin R BG - GCC
Kinesin R GC
Kinesin R AG - CBC
Kinesin R AG - MTCP
Kinesin R AG - MTCD
Kinesin R AG - GCC
Kinesin R BPM - CBC
Kinesin R BPM - MTCP
Kinesin R BPM - MTCD
Kinesin R BPM - GCC
Kinesin R APM - CBC
Kinesin R APM - MTCP
Kinesin R APM - MTCD
Kinesin R APM - GCC
### molecules averaged over time
Time [min]

**A7**

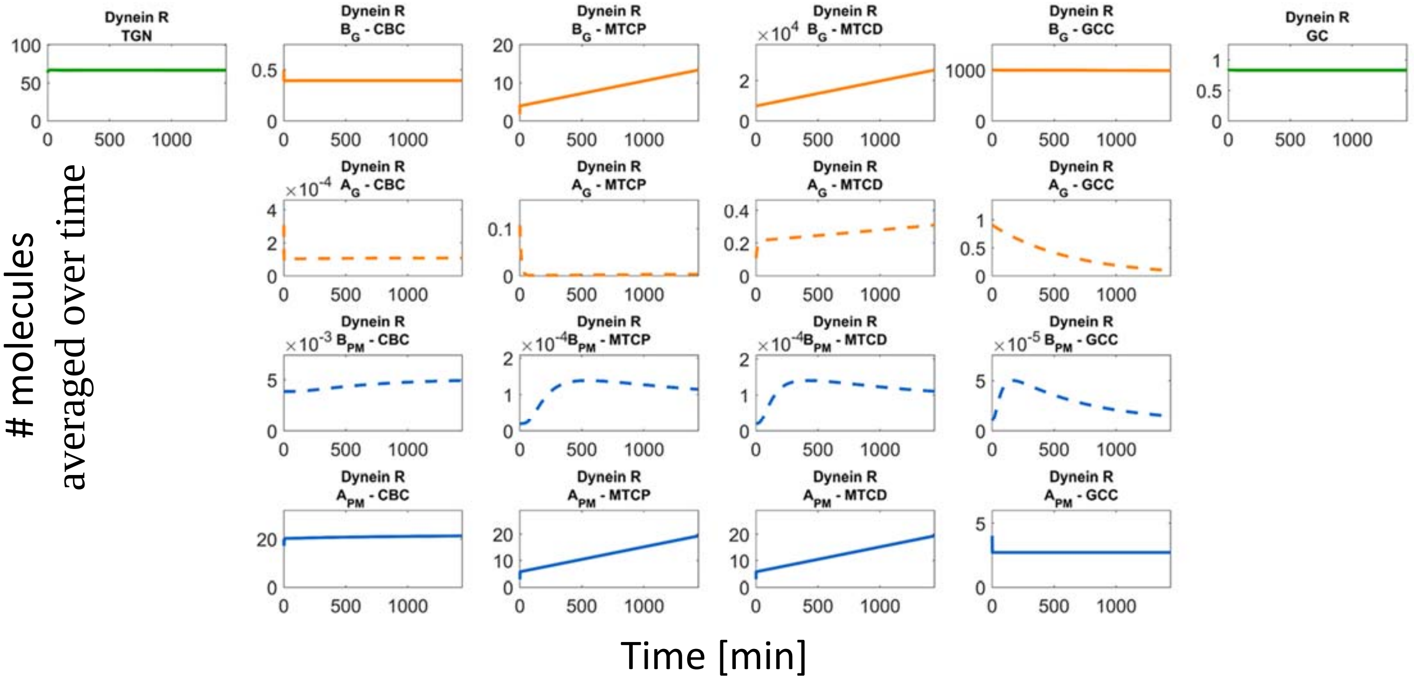

Dynein R TGN
Dynein R B_G - CBC
Dynein R B_G - MTCP
Dynein R B_G - MTCD
Dynein R B_G - GCC
Dynein R GC
Dynein R A_G - CBC
Dynein R A_G - MTCP
Dynein R A_G - MTCD
Dynein R A_G - GCC
Dynein R B_PM - CBC
Dynein R B_PM - MTCP
Dynein R B_PM - MTCD
Dynein R B_PM - GCC
Dynein R A_PM - CBC
Dynein R A_PM - MTCP
Dynein R A_PM - MTCD
Dynein R A_PM - GCC
### molecules averaged over time
Time [min]


**A8**

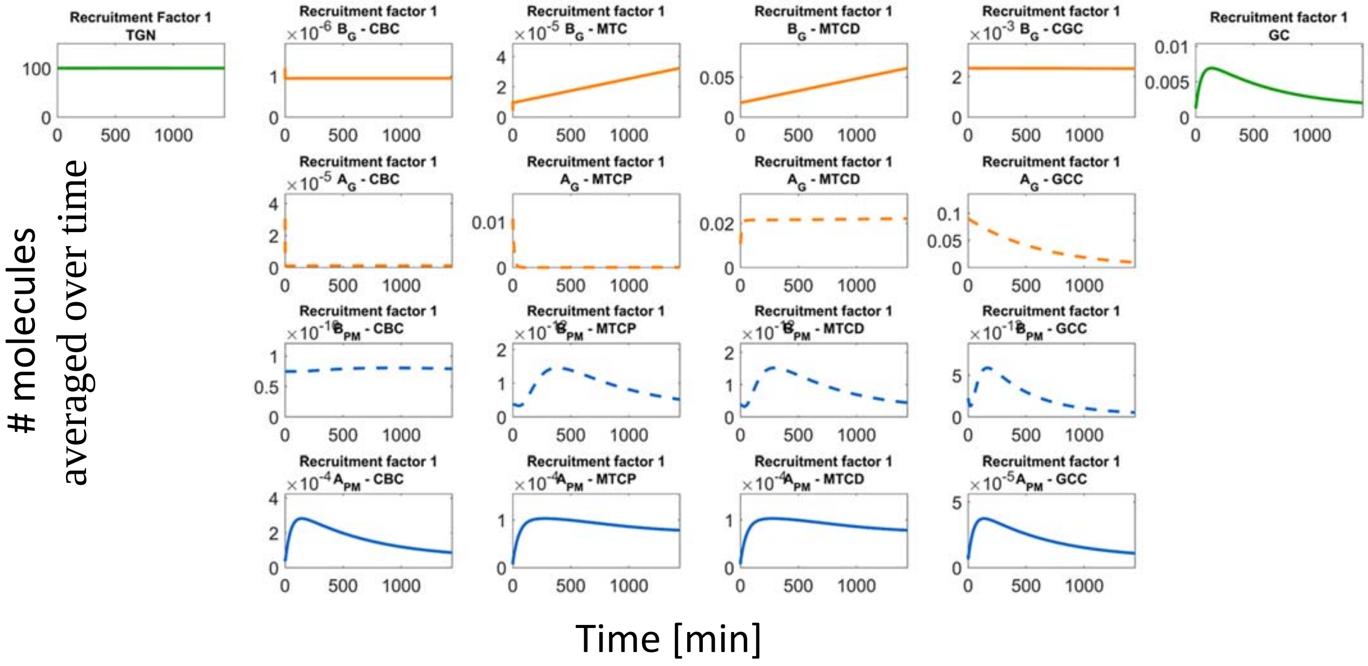

Recruitment Factor 1 TGN
Recruitment factor 1 B_G - CBC
Recruitment factor 1 B_G - MTC
Recruitment factor 1 B_G - MTCD
Recruitment factor 1 B_G - CGC
Recruitment factor 1 GC
Recruitment factor 1 A_G - CBC
Recruitment factor 1 A_G - MTCP
Recruitment factor 1 A_G - MTCD
Recruitment factor 1 A_G - GCC
Recruitment factor 1 B_PM - CBC
Recruitment factor 1 B_PM - MTCP
Recruitment factor 1 B_PM - MTCD
Recruitment factor 1 B_PM - GCC
Recruitment factor 1 A_PM - CBC
Recruitment factor 1 A_PM - MTCP
Recruitment factor 1 A_PM - MTCD
Recruitment factor 1 A_PM - GCC
### molecules averaged over time
Time [min]

A9

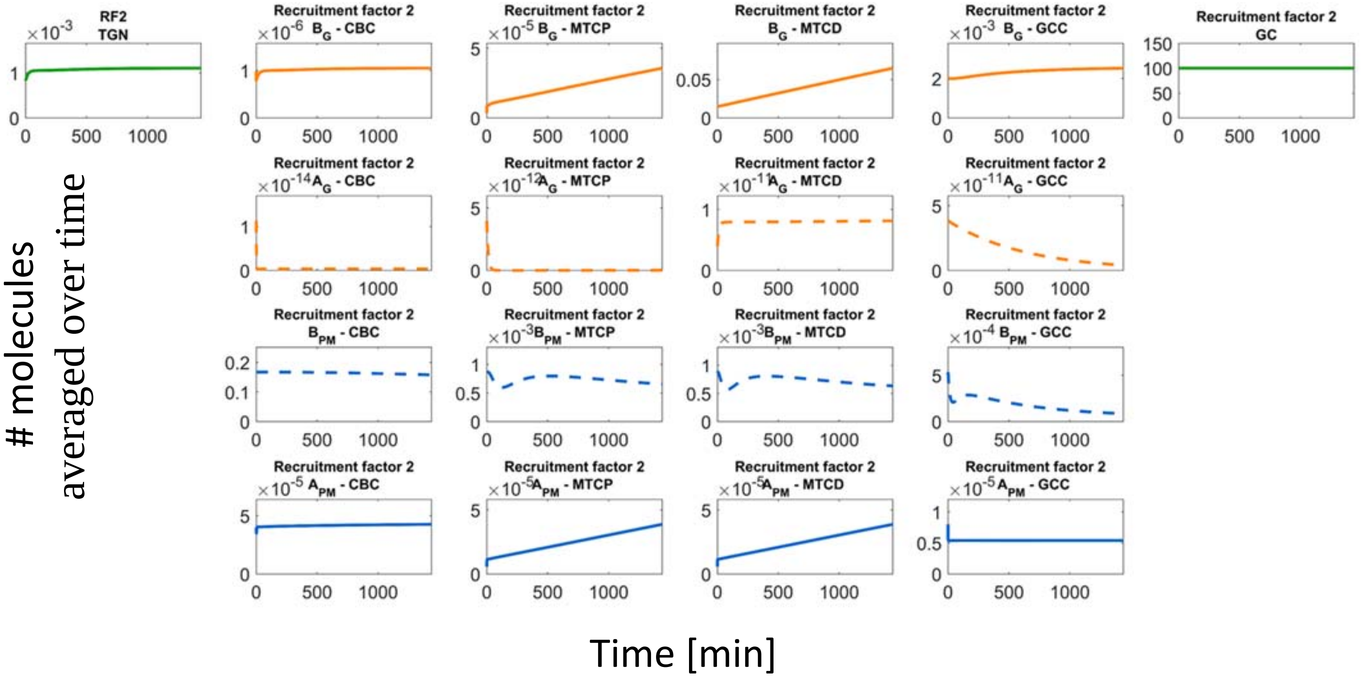
RF2
TGN
Recruitment factor 2
$B_G$ - CBC
Recruitment factor 2
$B_G$ - MTCP
Recruitment factor 2
$B_G$ - MTCD
Recruitment factor 2
$B_G$ - GCC
Recruitment factor 2
GC
Recruitment factor 2
$A_G$ - CBC
Recruitment factor 2
$A_G$ - MTCP
Recruitment factor 2
$A_G$ - MTCD
Recruitment factor 2
$A_G$ - GCC
Recruitment factor 2
$B_{PM}$ - CBC
Recruitment factor 2
$B_{PM}$ - MTCP
Recruitment factor 2
$B_{PM}$ - MTCD
Recruitment factor 2
$B_{PM}$ - GCC
Recruitment factor 2
$A_{PM}$ - CBC
Recruitment factor 2
$A_{PM}$ - MTCP
Recruitment factor 2
$A_{PM}$ - MTCD
Recruitment factor 2
$A_{PM}$ - GCC
### molecules
averaged over time
Time [min]

A10

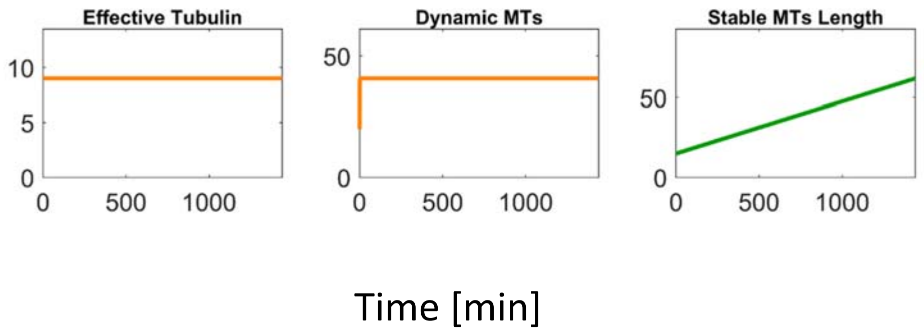
Effective Tubulin
Dynamic MTs
Stable MTs Length
Time [min]

B1

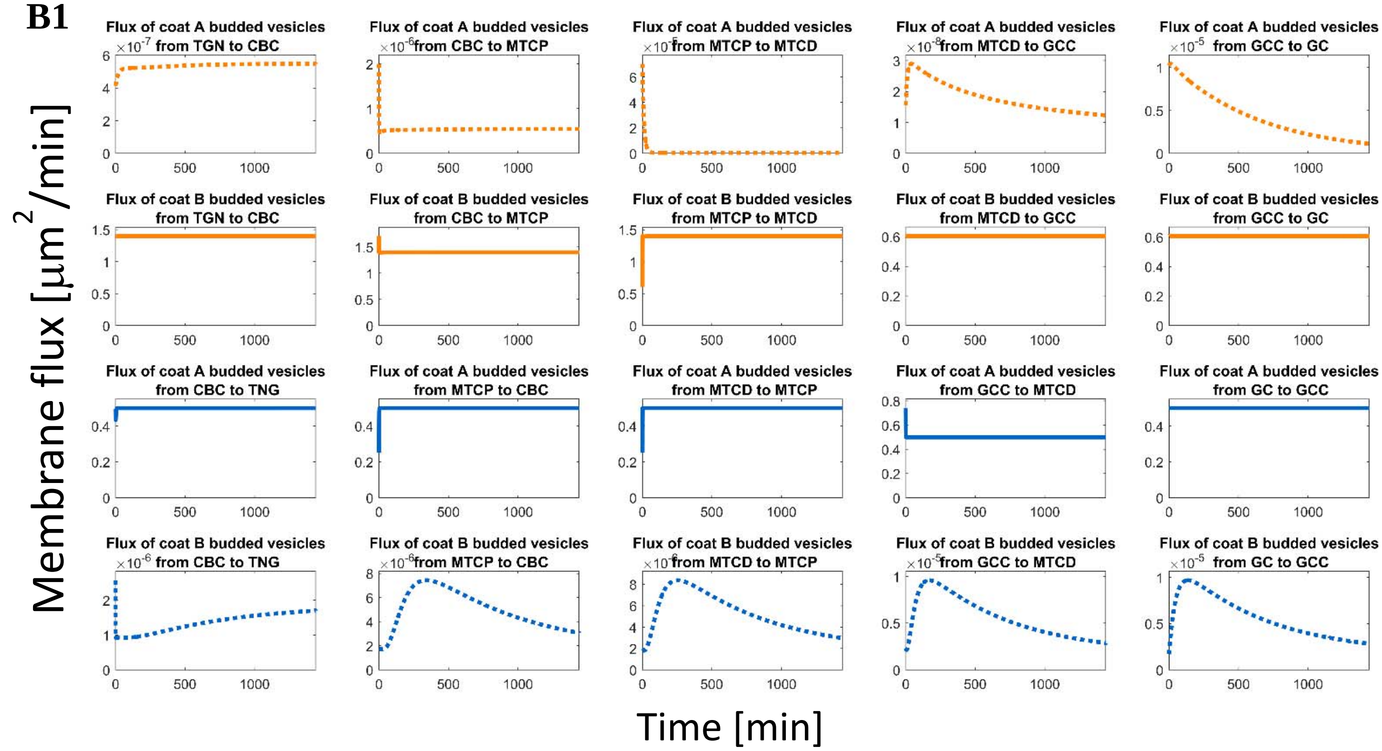
Flux of coat A budded vesicles from TGN to CBC
Flux of coat A budded vesicles from CBC to MTCP
Flux of coat A budded vesicles from MTCP to MTCD
Flux of coat A budded vesicles from MTCD to GCC
Flux of coat A budded vesicles from GCC to GC
Flux of coat B budded vesicles from TGN to CBC
Flux of coat B budded vesicles from CBC to MTCP
Flux of coat B budded vesicles from MTCP to MTCD
Flux of coat B budded vesicles from MTCD to GCC
Flux of coat B budded vesicles from GCC to GC
Flux of coat A budded vesicles from CBC to TNG
Flux of coat A budded vesicles from MTCP to CBC
Flux of coat A budded vesicles from MTCD to MTCP
Flux of coat A budded vesicles from GCC to MTCD
Flux of coat A budded vesicles from GC to GCC
Flux of coat B budded vesicles from CBC to TNG
Flux of coat B budded vesicles from MTCP to CBC
Flux of coat B budded vesicles from MTCD to MTCP
Flux of coat B budded vesicles from GCC to MTCD
Flux of coat B budded vesicles from GC to GCC
Membrane flux [μm²/min]
Time [min]

B2

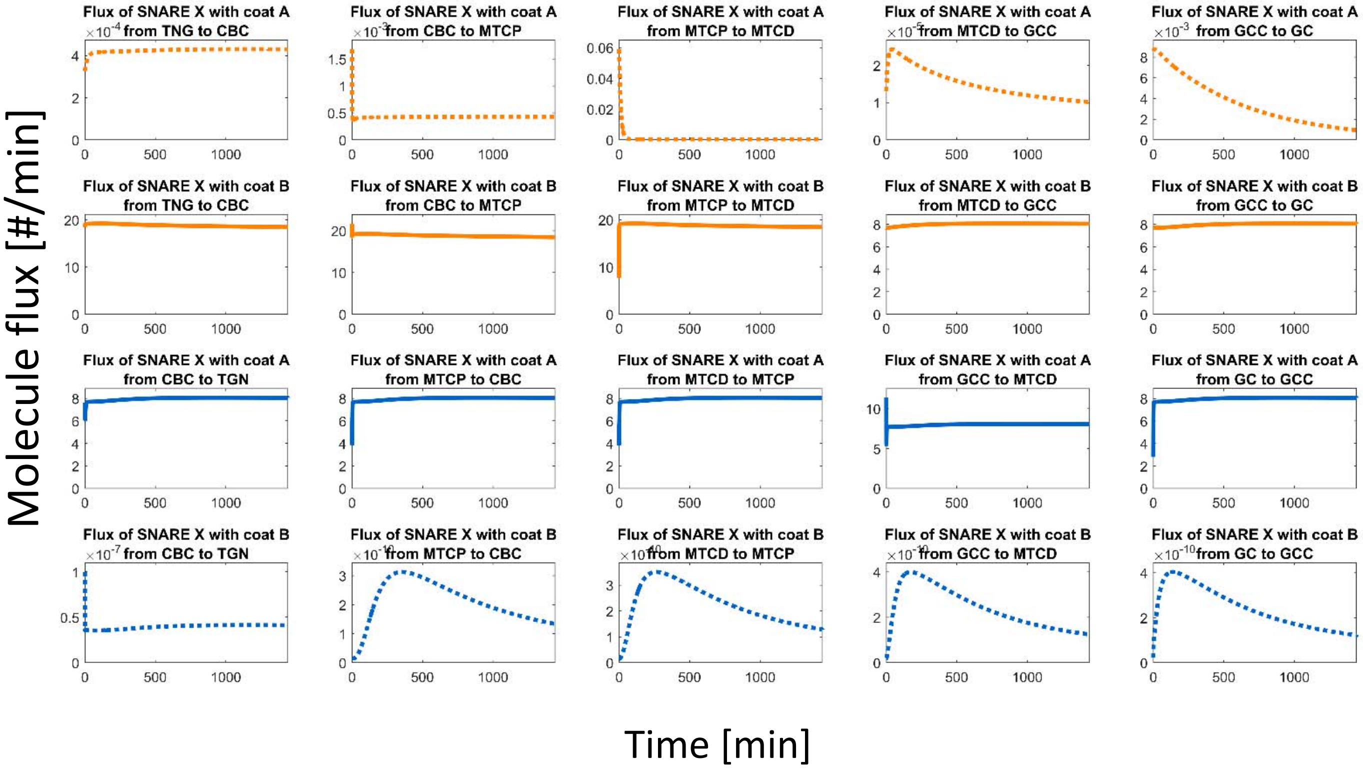
Flux of SNARE X with coat A from TNG to CBC
Flux of SNARE X with coat A from CBC to MTCP
Flux of SNARE X with coat A from MTCP to MTCD
Flux of SNARE X with coat A from MTCD to GCC
Flux of SNARE X with coat A from GCC to GC
Flux of SNARE X with coat B from TNG to CBC
Flux of SNARE X with coat B from CBC to MTCP
Flux of SNARE X with coat B from MTCP to MTCD
Flux of SNARE X with coat B from MTCD to GCC
Flux of SNARE X with coat B from GCC to GC
Flux of SNARE X with coat A from CBC to TGN
Flux of SNARE X with coat A from MTCP to CBC
Flux of SNARE X with coat A from MTCD to MTCP
Flux of SNARE X with coat A from GCC to MTCD
Flux of SNARE X with coat A from GC to GCC
Flux of SNARE X with coat B from CBC to TGN
Flux of SNARE X with coat B from MTCP to CBC
Flux of SNARE X with coat B from MTCD to MTCP
Flux of SNARE X with coat B from GCC to MTCD
Flux of SNARE X with coat B from GC to GCC
Molecule flux [#/min]
Time [min]

**B3**

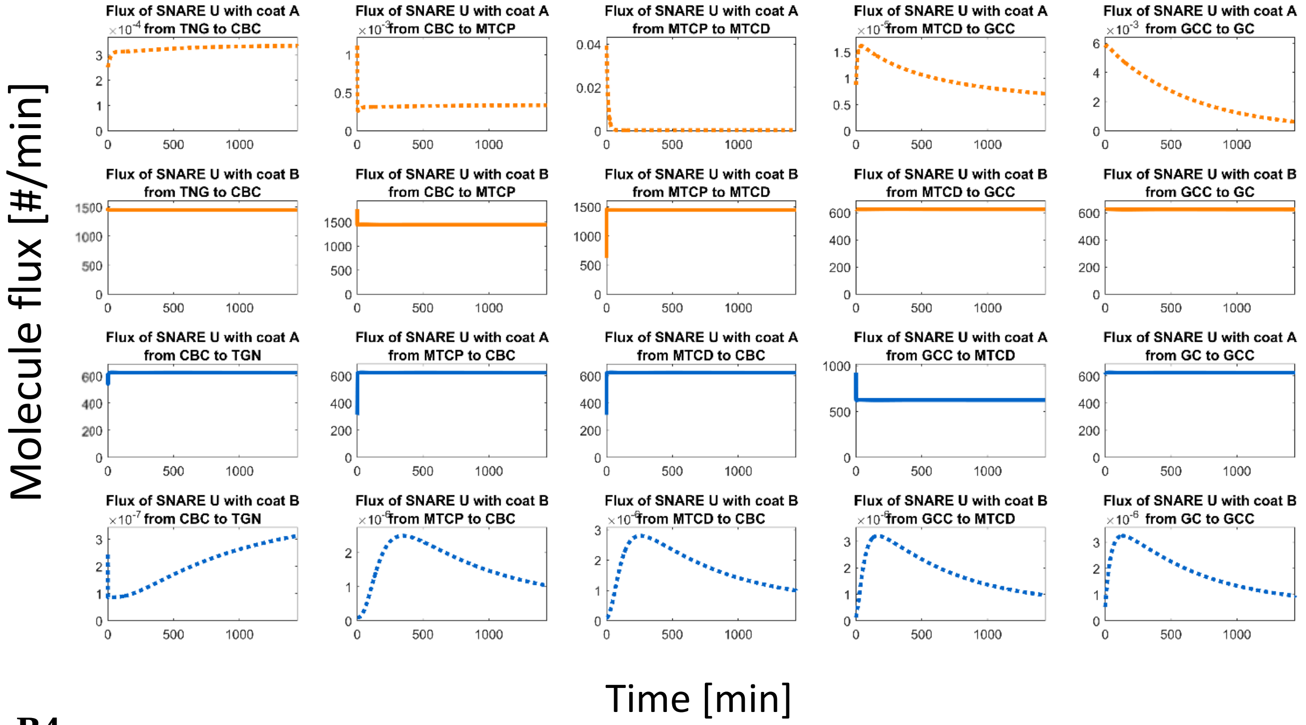

Flux of SNARE U with coat A from TNG to CBC
Flux of SNARE U with coat A from CBC to MTCP
Flux of SNARE U with coat A from MTCP to MTCD
Flux of SNARE U with coat A from MTCD to GCC
Flux of SNARE U with coat A from GCC to GC
Flux of SNARE U with coat B from TNG to CBC
Flux of SNARE U with coat B from CBC to MTCP
Flux of SNARE U with coat B from MTCP to MTCD
Flux of SNARE U with coat B from MTCD to GCC
Flux of SNARE U with coat B from GCC to GC
Flux of SNARE U with coat A from CBC to TGN
Flux of SNARE U with coat A from MTCP to CBC
Flux of SNARE U with coat A from MTCD to CBC
Flux of SNARE U with coat A from GCC to MTCD
Flux of SNARE U with coat A from GC to GCC
Flux of SNARE U with coat B from CBC to TGN
Flux of SNARE U with coat B from MTCP to CBC
Flux of SNARE U with coat B from MTCD to CBC
Flux of SNARE U with coat B from GCC to MTCD
Flux of SNARE U with coat B from GC to GCC
Molecule flux [#/min]
Time [min]


**B4**

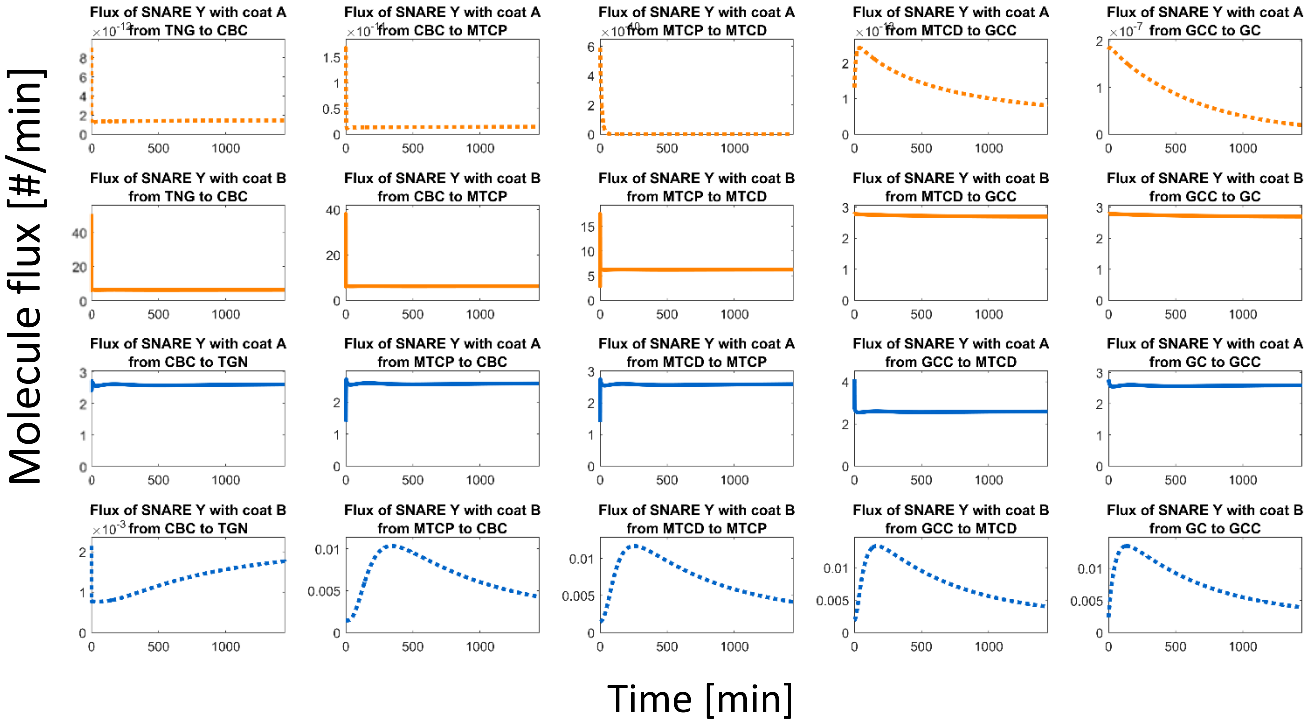

Flux of SNARE Y with coat A from TNG to CBC
Flux of SNARE Y with coat A from CBC to MTCP
Flux of SNARE Y with coat A from MTCP to MTCD
Flux of SNARE Y with coat A from MTCD to GCC
Flux of SNARE Y with coat A from GCC to GC
Flux of SNARE Y with coat B from TNG to CBC
Flux of SNARE Y with coat B from CBC to MTCP
Flux of SNARE Y with coat B from MTCP to MTCD
Flux of SNARE Y with coat B from MTCD to GCC
Flux of SNARE Y with coat B from GCC to GC
Flux of SNARE Y with coat A from CBC to TGN
Flux of SNARE Y with coat A from MTCP to CBC
Flux of SNARE Y with coat A from MTCD to MTCP
Flux of SNARE Y with coat A from GCC to MTCD
Flux of SNARE Y with coat A from GC to GCC
Flux of SNARE Y with coat B from CBC to TGN
Flux of SNARE Y with coat B from MTCP to CBC
Flux of SNARE Y with coat B from MTCD to MTCP
Flux of SNARE Y with coat B from GCC to MTCD
Flux of SNARE Y with coat B from GC to GCC
Molecule flux [#/min]
Time [min]

B5

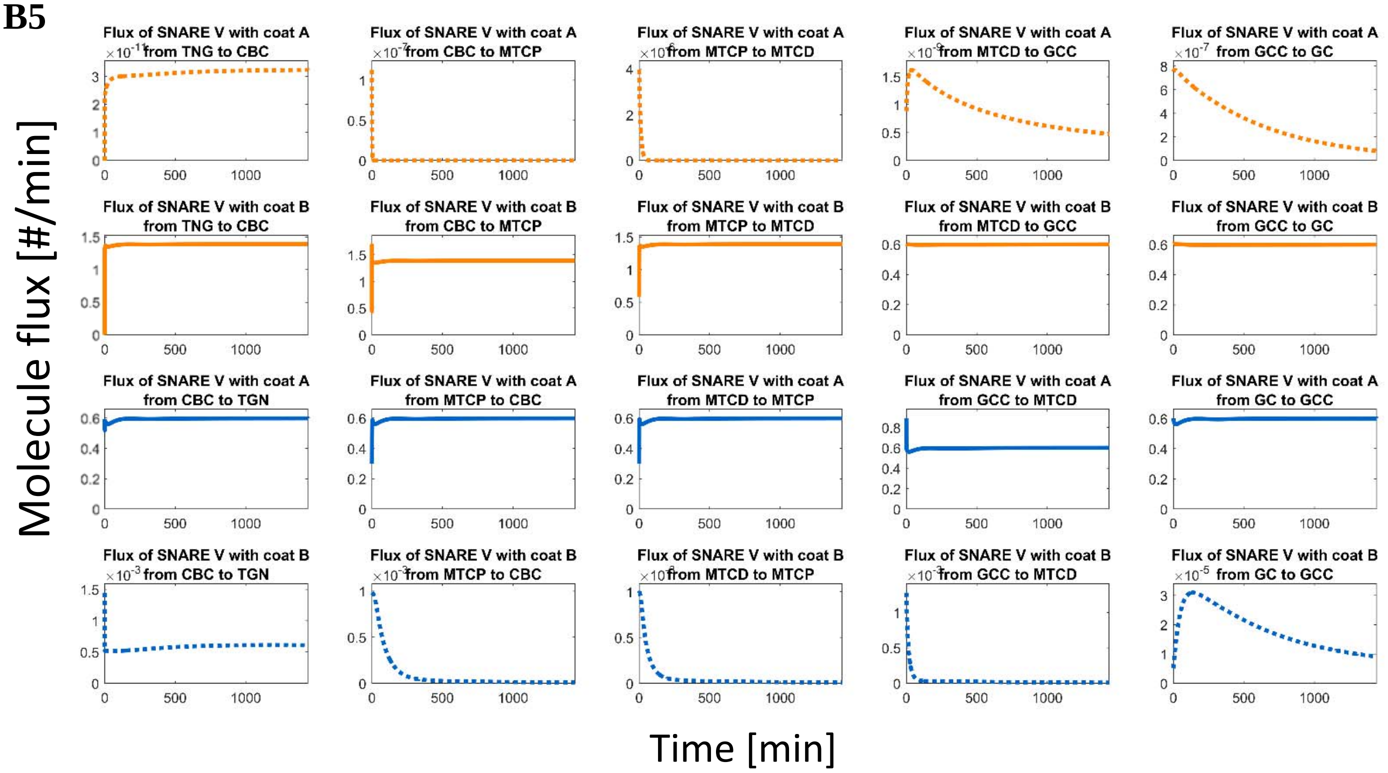

Flux of SNARE V with coat A from TNG to CBC
Flux of SNARE V with coat A from CBC to MTCP
Flux of SNARE V with coat A from MTCP to MTCD
Flux of SNARE V with coat A from MTCD to GCC
Flux of SNARE V with coat A from GCC to GC
Flux of SNARE V with coat B from TNG to CBC
Flux of SNARE V with coat B from CBC to MTCP
Flux of SNARE V with coat B from MTCP to MTCD
Flux of SNARE V with coat B from MTCD to GCC
Flux of SNARE V with coat B from GCC to GC
Flux of SNARE V with coat A from CBC to TGN
Flux of SNARE V with coat A from MTCP to CBC
Flux of SNARE V with coat A from MTCD to MTCP
Flux of SNARE V with coat A from GCC to MTCD
Flux of SNARE V with coat A from GC to GCC
Flux of SNARE V with coat B from CBC to TGN
Flux of SNARE V with coat B from MTCP to CBC
Flux of SNARE V with coat B from MTCD to MTCP
Flux of SNARE V with coat B from GCC to MTCD
Flux of SNARE V with coat B from GC to GCC
Molecule flux [#/min]
Time [min]


B6

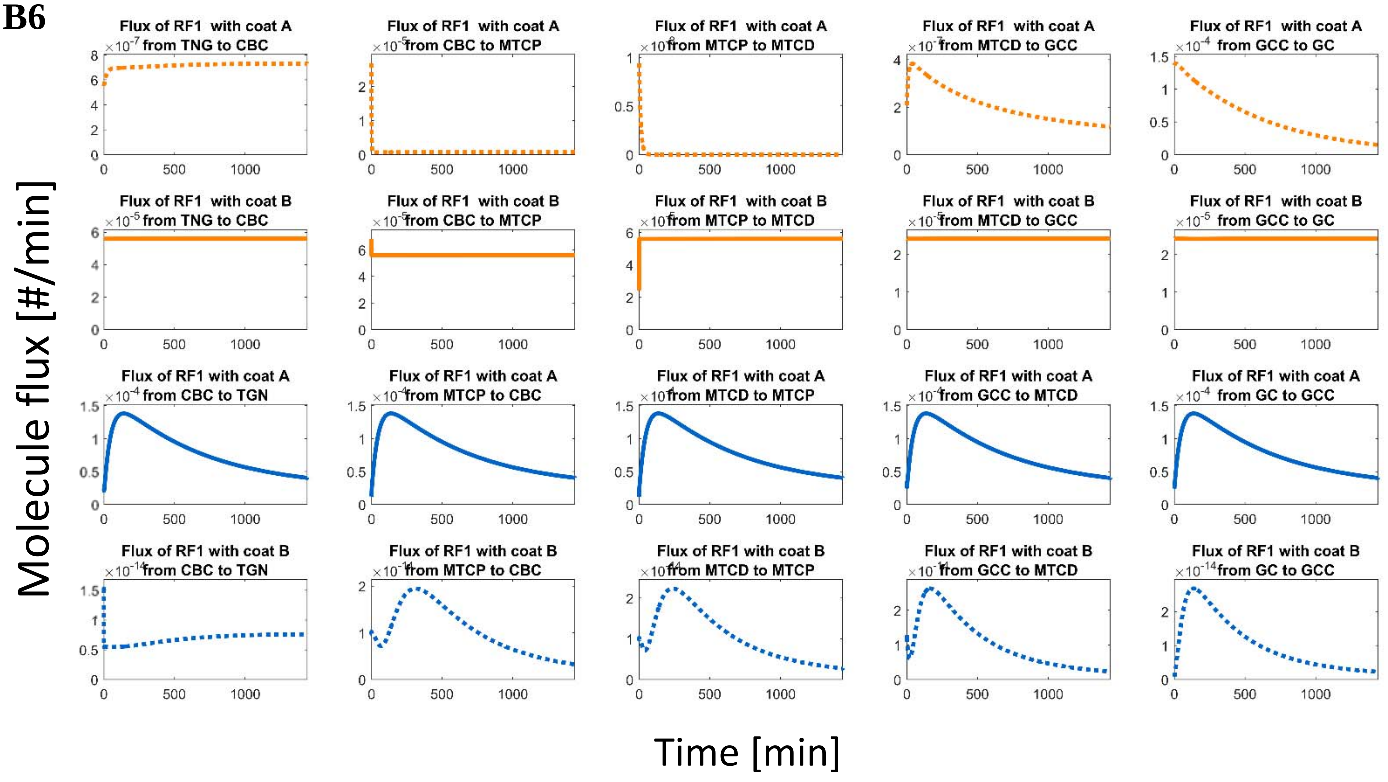

Flux of RF1 with coat A from TNG to CBC
Flux of RF1 with coat A from CBC to MTCP
Flux of RF1 with coat A from MTCP to MTCD
Flux of RF1 with coat A from MTCD to GCC
Flux of RF1 with coat A from GCC to GC
Flux of RF1 with coat B from TNG to CBC
Flux of RF1 with coat B from CBC to MTCP
Flux of RF1 with coat B from MTCP to MTCD
Flux of RF1 with coat B from MTCD to GCC
Flux of RF1 with coat B from GCC to GC
Flux of RF1 with coat A from CBC to TGN
Flux of RF1 with coat A from MTCP to CBC
Flux of RF1 with coat A from MTCD to MTCP
Flux of RF1 with coat A from GCC to MTCD
Flux of RF1 with coat A from GC to GCC
Flux of RF1 with coat B from CBC to TGN
Flux of RF1 with coat B from MTCP to CBC
Flux of RF1 with coat B from MTCD to MTCP
Flux of RF1 with coat B from GCC to MTCD
Flux of RF1 with coat B from GC to GCC
Molecule flux [#/min]
Time [min]

**B7**

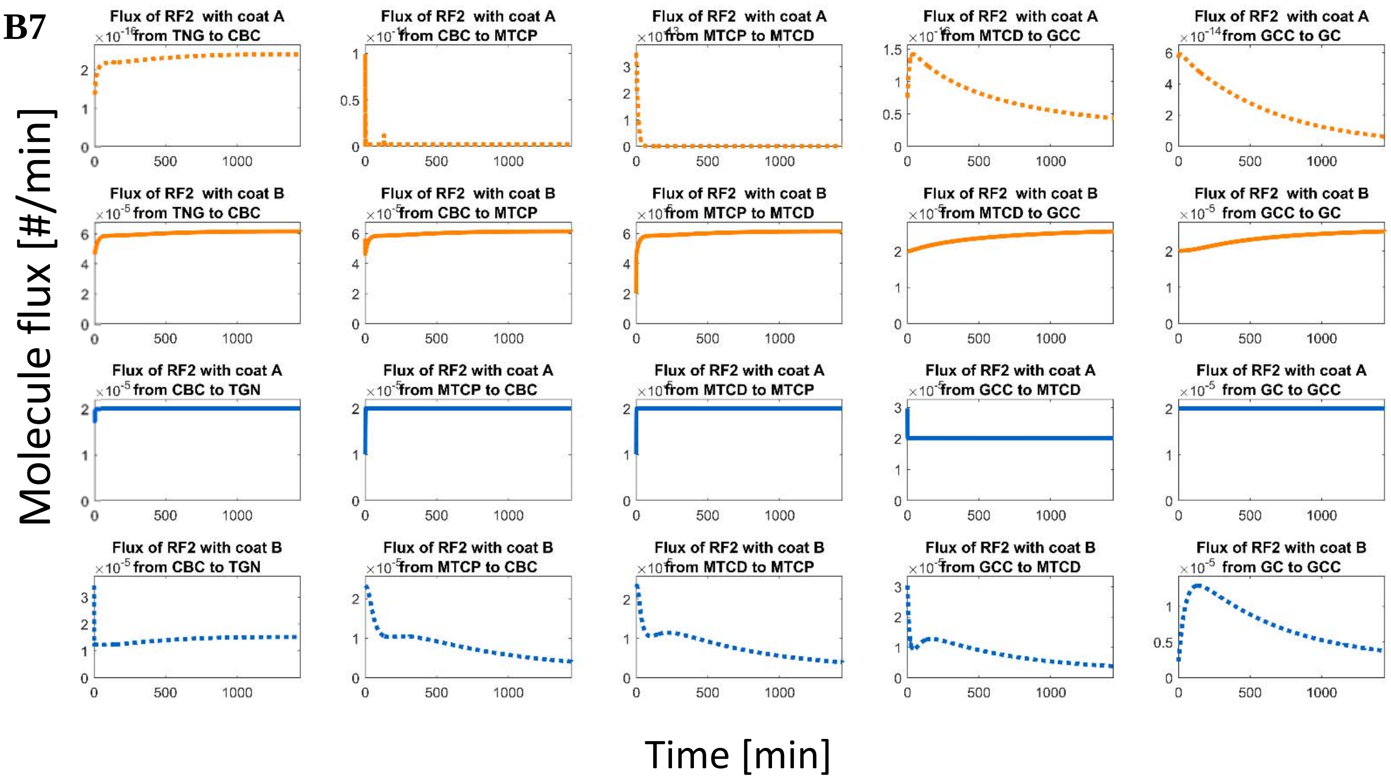


**Supplementary Figure 10: Steady state distribution and fluxes of proteins and membrane vesicles.** (**A**) At steady state all proteins and membrane amounts stay constant in the different compartments, except in the neurite shaft cytoplasm (NSC-P & NSC-D) and the neurite shaft. Neurite shaft growth is facilitated by continuous membrane addition, so the membrane surface area of the neurite shaft grows over the time. The NSC-D grows parallel to the neurite shaft and acts as a sink for transport vesicles, thereby accumulating vesicle membrane and vesicle membrane proteins. The figures show the membrane surface areas or protein amounts in each compartment within each set of vesicles. $B_G$ refers to anterograde vesicles that bud from the TGN with the coat protein B, $A_G$ refers to anterograde vesicles that bud from the TGN with the coat A, $B_{PM}$ refers to retrograde vesicles that bud from the GC-PM with coat protein B and $A_{PM}$ to retrograde vesicles that bud from the GC with coat protein A. Green lines refer to TGN or growth cone plasma membrane (GC-PM), orange lines refer to anterograde moving vesicles ($B_G$, $A_G$) and blue lines to retrograde moving vesicles ($B_{PM}$, $A_{PM}$). Solid lines represent those vesicles that are mainly responsible for membrane transport in the indicated direction (i.e. $B_G$ in the anterograde direction and $A_{PM}$ in the retrograde direction). Figure 3(A1-A9) shows distribution of anterograde and retrograde (vesicles, SNAREs-Y, SNAREs-V, SNAREs-X, SNAREs-U, Kinesin receptor-k, Dynein receptor-d, Recruitment factor-1, Recruitment factor-2) and A10 shows effective tubulin concentration of the model, number of dynamic microtubules and dynamics of stable microtubules. **(B)** Figure (B1-B11) shows all fluxes, i.e. membrane or protein transfers from one compartment into the other are shown. Colors and line styles are the same as in (A).

#### Supplementary Figure – 11

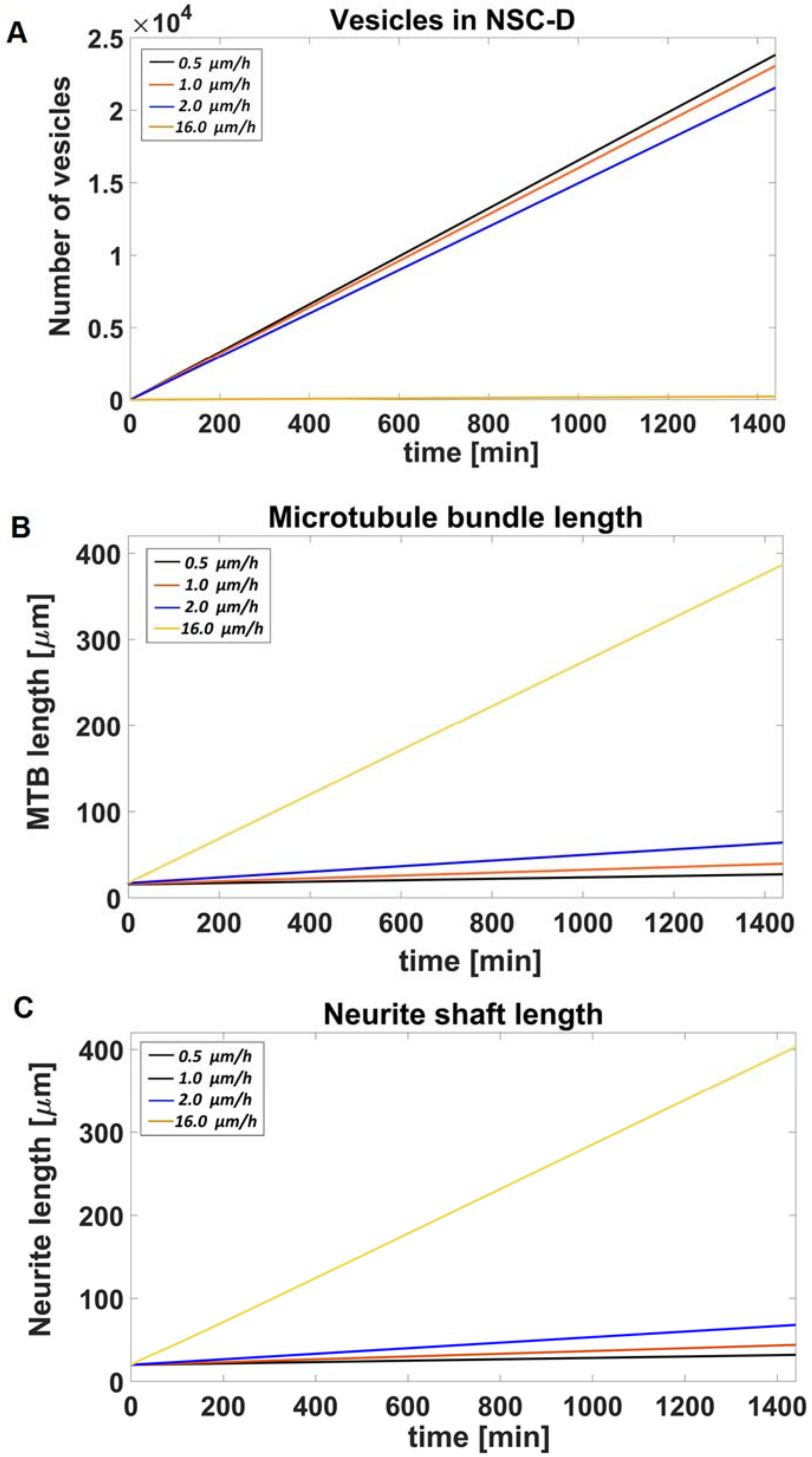

**Supplementary Figure 11: Multicompartmental ODE model for axonal outgrowth under different drug treatment conditions.** Axonal outgrowth velocities varies with different drug combination and shows synergistic effect for four drugs combinations (HU210, IL-6, APC and Taxol). HU210 and IL-6 applied at cell body which increase the capacity of membrane production at TGN, Taxol and APC applied at injury site where Taxol increase microtubule stabilization and APC clears inhibitory environment at injury site which increase vesicles mobility from vesicles reservoir (neurite shaft cytoplasm distal (NSC-D)) to growth cone plasma membrane (GC-PM). We built multi-compartmental ordinary differential equation based (ODEs) model by introducing biological roles of above mentioned drugs and validated with the growth velocities of neurite.

**(A)** Neurite shaft cytoplasm distal (NSC-D) compartment work as a reservoir for anterograde vesicles because Tau gradient increase from cell body cytoplasm to growth cone (Tau compete with kinesin to bind with microtubules in nerite shaft cytoplasm) and hence, binding rates of kinesin motor proteins decreases from NSC-P, NSC-D to GCC compartments. For combination of four drugs, neurite outgrow with 16 µm/h by mobilizing approximately 23,000 vesicles from NSC-D to GCC compartment.

**(B)** Axon outgrow with velocities V=0.5 µm/h, V=1 µm/h, V=2 µm/h, V=16 µm/h for control and different drug combinations. HU210 and IL-6 applied at cell body in all cases which increase the vesicles production rate at TGN. Axon outgrow with velocity V=1 µm/h when Taxol applied at injury site which stabilize microtubules, but overall growth of microtubule bundle inhibited by external debris. After applying APC gel foam at injury site, axon outgrow with velocity V=2 µm/h because it clears inhibitory environment and hence vesicle moves from reservoir NSC-D compartment and fuse with growth cone plasma membrane. Combination of four drugs (HU210, IL-6, APC and Taxol) shows synergistic effect because Taxol stabilizes microtubules and APC clears inhibitory environment and hence axon outgrow with velocity V=16 µm/h.

**(C)** To simulate microtubule growth, we consider the two different pools of microtubules, stable and dynamic microtubules. After nucleation new MTs are added to the pool of dynamic MTs that is characterized by alternating phases of growth and catastrophic breakdown. The frequency and duration of these phases depend on the tubulin concentration and the GTP hydrolysis rate. Taxol used at injury site which reduce the rate of hydrolysis, consequently microtubules stabilization rate increase.

*Materials*

**Antibodies**

| **Name** | **Company** | **Cat. Number** |
|---|---|---|
| Purified anti-tubulin Beta 3 (TUBB3) mouse | Biolegend | 801202 |
| STAT3 (124H6) mouse | Cell Signaling | 9139S |
| phospho YB1 (S102) (C34A2) Rabbit | Cell Signaling | 2900S |
| p35/25 (C64810) Rabbit | Cell Signaling | 2680S |
| Anti Albumin Chicken | Sigma | SAB3500217 |
| phospho STAT3 (TYR705) (D3A7) Rabbit | Cell Signaling | 9145S |
| Alexa Fluor 488 mouse | Invitrogen | A11029 |
| Alexa Fluor 568 mouse | Invitrogen | A11031 |
| Anti Cholera Toxin B Subunit Goat | List Biological Lab | 703 |
| GAP-43 Antibody | Novus Biologicals | NB300-143 |

**Media and Supplements**

| **Name** | **Company** | **Cat. Number** |
|---|---|---|
| Neurobasal | Thermo Fisher Scientific | 2103-049 |
| Pen-Strep | Thermo Fisher Scientific | 15140-122 |
| Feta bovine serum | Thermo Fisher Scientific | 10439-026 |
| L-Glutamine | Thermo Fisher Scientific | 25030-081 |
| B27 Supplement | Thermo Fisher Scientific | 17504044 |
| Papain from papaya extract | Sigma | P5306 |
| Deoxyribonuclease I from bovine pancreas | Sigma | DN25 |
| Poly-L-lysine hydrobromide | Sigma | P1274 |
| OptiPrep | Sigma | D1556 |

**Fixing and Staining**

| Name | Company | Cat. Number |
|---|---|---|
| Triton X 100 | Sigma | X100 |
| Phosphate Buffer Saline (10X) | Boston Bioproducts | BM-220 |
| Normal Donkey Serum | Jackson Immuno Research | 017-000-121 |
| Tetrahydrofuran | Sigma | 401757 |
| Sucrose | Sigma | S7905 |
| Paraformyldehyde 16% solution | Electron Microscopy Science | 15710-S |
| Hoeschst 33342 | Invitrogen | 953557 |
| Mountant- Permafluor | Thermo Fisher Scientific | TA-030-FM |
| OCT Compound | Sakura Tissue Tek | 4583 |
| Corning Cover Glass | Corning | 2980-245 |
| 50 mm Glass Bottom Dishes | Mattek Corp. | P50G-1.5-30-F |
| Standard neuron device | XONA | SND150 |

**Western Blot**

| Name | Company | Cat. Number |
|---|---|---|
| SDS 4X Buffer | Boston Bioproducts | BP-110R |
| Albumin Standard 2mg/ml | Thermo Fisher Scientific | 23210 |
| BioRad Protein Assay | BioRad | 5000006 |
| mini-protean tgx precast gels | BioRad | 456-1094 |
| 30% Acrylimade/Bis Solution | BioRad | 160158 |
| Odyssey Blocking Buffer (TBS) | Li-COR | 927-500000 |
| Halt Protease Phosphatase Inhibitors | Thermo Fisher Scientific | 78443 |
| IRDye® Goat anti-Rabbit 800CW Secondary | Li-COR | 925-32211 |
| IRDye® Goat anti-Mouse 680RD Secondary | Li-COR | 925-68070 |
| IRDye® 800CW Donkey anti-Chicken | Li-COR | 925-32218 |
| Tris-buffered saline (10X) | Boston Bioproducts | BM-300 |
| Tris-glycine SDS Running Buffer (10X) | Boston Bioproducts | BP-150 |
| TWEEN® 20 | Sigma | P9416 |
| Goat anti-Rabbit IgG (H+L) Secondary Antibody, HRP | Thermo Fisher Scientific | 31460 |
| Goat anti-Mouse IgG (H+L) Secondary Antibody, HRP | Thermo Fisher Scientific | 31430 |
| PMSF (Phenylmethylsulfonyl fluoride) | Sigma | 10837091001 |
| Pepstatin | Sigma | 10253286001 |

| | | |
|---|---|---|
| RIPA Buffer (10X) | Cell Signaling | 9806 |
| Ammonium Persulfate | Thermo Fisher Scientific | 17874 |
| Sodium dodecyl sulfate | Sigma | L3771 |
| Methanol | Sigma | 179337 |
| Transfer Buffer (10X) | Boston Bioproducts | BP-190 |
| Calcein, AM | Thermo Fisher Scientific | C3200MP |

**<u>Drugs</u>**

| **Name** | **Company** | **Cat. Number** |
|---|---|---|
| Human Activated Protein C | Haematologic Technologies | HCAPC-0080 |
| HU-210 | SIGMA | H7909 |
| Interleukin-6 from rat, recombinant | SIGMA | I0406 |
| Paclitaxel (Taxol) | SIGMA | T7402 |
| Dimethyl Sulphide, sterile | SIGMA | D2650 |

**<u>Surgeries</u>**

| **Name** | **Company** | **Cat. Number** |
|---|---|---|
| Forane Isoflurane | Baxter | 10019-360-40 |
| Buprenorphine HCL | ParPharmaceutical | 42023-179-01 |
| Vetasan Ointment | Kinetic Vet | 9006-06-00 |
| Alcohol Swabs | Becton Dickinson | 08290-3268-95-326895 |
| Vetbond | 3M | 14695B |
| Nanofil Syringe | World Precision Instruments | nanofil |
| Neomycin ointment | Bausch + Lomb | 24208-780-55 |
| Ketamin HCL | Vedco Inc | 50989-161-06 |
| AnaSed Xylazine | AKORN | 59399-111-50 |
